# Rare k-mers reveal centromere haplogroups underlying human diversity and cancer translocations

**DOI:** 10.1101/2025.07.26.666712

**Authors:** Yuichi Shiraishi, Yotaro Ochi, Masahiro Sugawa, Yoshitaka Sakamoto, Keisuke Kimura, Taro Tsujimura, Ai Okada, Rurika Okuda, Shinichi Namba, Tsubasa Miyauchi, Raul N. Mateos, Hajime Suzuki, Kenichi Chiba, Yu Ito, Wataru Nakamura, Fumiharu Ohka, Kazuya Motomura, Takuya Yamamoto, Yosuke Kawai, Yukinori Okada, Hiromichi Suzuki, Motohiro Kato, Ryuta Saito, Erik Garrison, Glennis A. Logsdon, Seishi Ogawa

## Abstract

Centromeres are among the most diverse and dynamically evolving regions of the human genome and are commonly affected in various human cancers. However, organized into highly repetitive α-satellite higher-order repeats (HORs), human centromere sequences have long resisted detailed genomic analysis. Although the development of long-read sequencing platforms has enabled the analysis of complete centromere sequences, their application to a large set of samples is still largely limited, preventing our understanding of centromere variation and haplotype structures across large human populations and the structural basis of centromere-involving translocations in cancer. Here we show that rare k-mers present in centromeric regions can serve as effective markers for dissecting the complexity of centromere structure, particularly that of active α-satellite HOR arrays (aHOR arrays), across human populations and for understanding centromere-involving abnormalities in cancer. Based on rare k-mer-based clustering, centromere aHOR arrays are clustered into discrete haplogroups (aHOR-HGs) with distinct structural features. These k-mers were also used to develop a framework that enables the inference of haplogroups in a given sample based on short-read whole genome sequencing (WGS) data (ascairn). By applying ascairn to large-scale human population datasets (*n* > 3,300), we revealed the diversity of aHOR-HGs and their geographic histories across populations. The rare k-mer-based approach was also applied to investigate the structure of 1p/19q co-deletion, a highly recurrent centromere-involving translocation in *IDH*-mutated oligodendrogliomas. Analyzing short-read WGS data from 142 cases with 1p/19q co-deletion using rare k-mers, we showed that breakpoints of 1p/19q co-deletion were mapped to aHOR arrays in chromosomes 1 (*D1Z7*) and 19 (*D19Z3*), which was validated by long-read sequencing of two 1p/19q co-deletion-positive cases. Notably, the translocation preferentially involved haplogroups composed of haplotypes containing larger regions susceptible to rearrangement. These results highlight the role of rare k-mers in dissecting the complexity of centromere sequences and their evolutionary history as well as understanding centromere-involving abnormalities associated with human diseases.

## Introduction

Centromeres are among the most diverse and rapidly evolving parts of the human genome. They are composed of a bulk of α-satellite DNA, consisting of ∼171 bp monomeric units, tandemly repeated to form higher-order repeat (HOR) arrays^1,2^. Among these arrays, those associated with CENP-A protein, the hallmark of active centromere^3^, are referred to as “active” α-satellite HOR arrays (aHOR arrays). Harboring a CENP-A-mediated kinetochore attachment site, these arrays play an essential role in centromere function and are implicated in centromere-involving translocations and aneuploidy^4–8^ frequently found in cancer. Unlike recombination-active regions, aHOR arrays are thought to retain lineage structure over long evolutionary time due to suppressed recombination^9^, which provides a unique opportunity to observe the cumulative impact of mutation, drift, and selection in an essential functional genomic compartment without the confounding effects of recombination^10^. However, their highly repetitive nature has long prevented detailed analysis of human centromeres^11^. Even with major advances in sequencing technologies over the past decades, centromeric regions have remained largely inaccessible to short-read platforms, persisting as “dark matter” regions in genome science^12,13^.

A major breakthrough has been made through recent advances in long-read sequencing technologies, which, coupled with the state-of-the-art genome assemblers, have enabled the reconstruction of complete sequences of the human genome, including centromeres^14–17^. Leveraging these advances, large-scale efforts by initiatives, such as the Human Pangenome Reference Consortium (HPRC)^18^ and the Human Genome Structural Variation Consortium (HGSVC)^19,20^, have generated a growing catalog of fully assembled centromere haplotypes, revealing large structural diversity of human centromeres across chromosomes and individuals^20,21^. Nevertheless, the evolutionarily structured variations of centromere haplotypes across populations and their relevance in human diseases, including cancer, remain poorly understood, because the time and cost of high-coverage long-read sequencing continue to preclude population-scale studies. The need for high-molecular-weight DNA also prevents large-scale analysis of clinical samples.

In this study, we identified rare k-mers as new polymorphic markers for a better understanding of the complexity of human centromere structure. We first demonstrated that haplotypes of aHOR arrays (aHOR-haps) were clustered into distinct haplogroups (aHOR-HGs) using short-nucleotide sequence markers, i.e., rare k-mers, and based on this, developed a probabilistic framework, “ascairn”, to assign aHOR-haps of a given individual to their haplogroups and to find their closest proxy within the reference haplotype set (reference aHOR-hap panel), using short-read whole-genome sequencing (WGS) data. ascairn was then applied to the WGS datasets from large-scale human genome projects to investigate the global structure of human centromeres across different populations with their evolutionary history. It was also used for the analysis of the class-defining centromere translocation, der(1;19)(q10;p10), or more commonly referred to as 1p/19q co-deletion^22–24^, found in *IDH*-mutated oligodendrogliomas (ODGs). Through the analysis of WGS data from a large cohort of ODG patients, we revealed the breakpoint structure of 1p/19q co-deletion within aHOR arrays in the relevant chromosomes, providing new insight into the mechanism of these translocations.

### Clustering aHOR-haps by rare k-mers

Although aHOR arrays are highly homogeneous, they still exhibit subtle sequence variations (Fig. 1a)^25^. To capture these subtle variations, we focused on rare k-mers: short subsequences that occur only infrequently within those aHOR arrays. We first extracted highly reliable aHOR-hap sequences including their flanking 100 kb sequences from publicly available datasets of chromosome-scale phased diploid assemblies^14,17,18,20,21^ derived from 118 individuals (Methods). In total, 2,930 aHOR-haps (20 to 165 per chromosome; median: 139) were obtained for each chromosome (ED_Fig. 1a and Supplementary Table 1), which we hereafter refer to as the “reference aHOR-hap panel”. Then, we identified a rare k-mer set (*k* = 27) for each chromosome that appeared only once or twice per aHOR-hap sequence. Redundant rare k-mers and those likely arising from sequencing errors were removed through rigorous filtering (Methods), yielding a total of 460,012 such rare k-mers across all chromosomes with a median of 19,213 (range: 5,673–45,407) per chromosome (ED_Fig. 1b). The median number of rare k-mers per aHOR-hap was 885.5 (range: 147–2,400) (Fig. 1b). In most aHOR arrays, the density of rare k-mers was higher at both ends compared to the array center (Fig. 1c and ED_Fig. 1c), consistent with the higher sequence diversity typically observed in array peripheries^15,26^. African populations exhibited significantly higher numbers of rare k-mers compared to other populations (Fig. 1d and ED_Fig. 2a), likely reflecting a higher diversity of African populations as suggested by studies on single nucleotide polymorphisms^27,28^.

**Figure 1:**
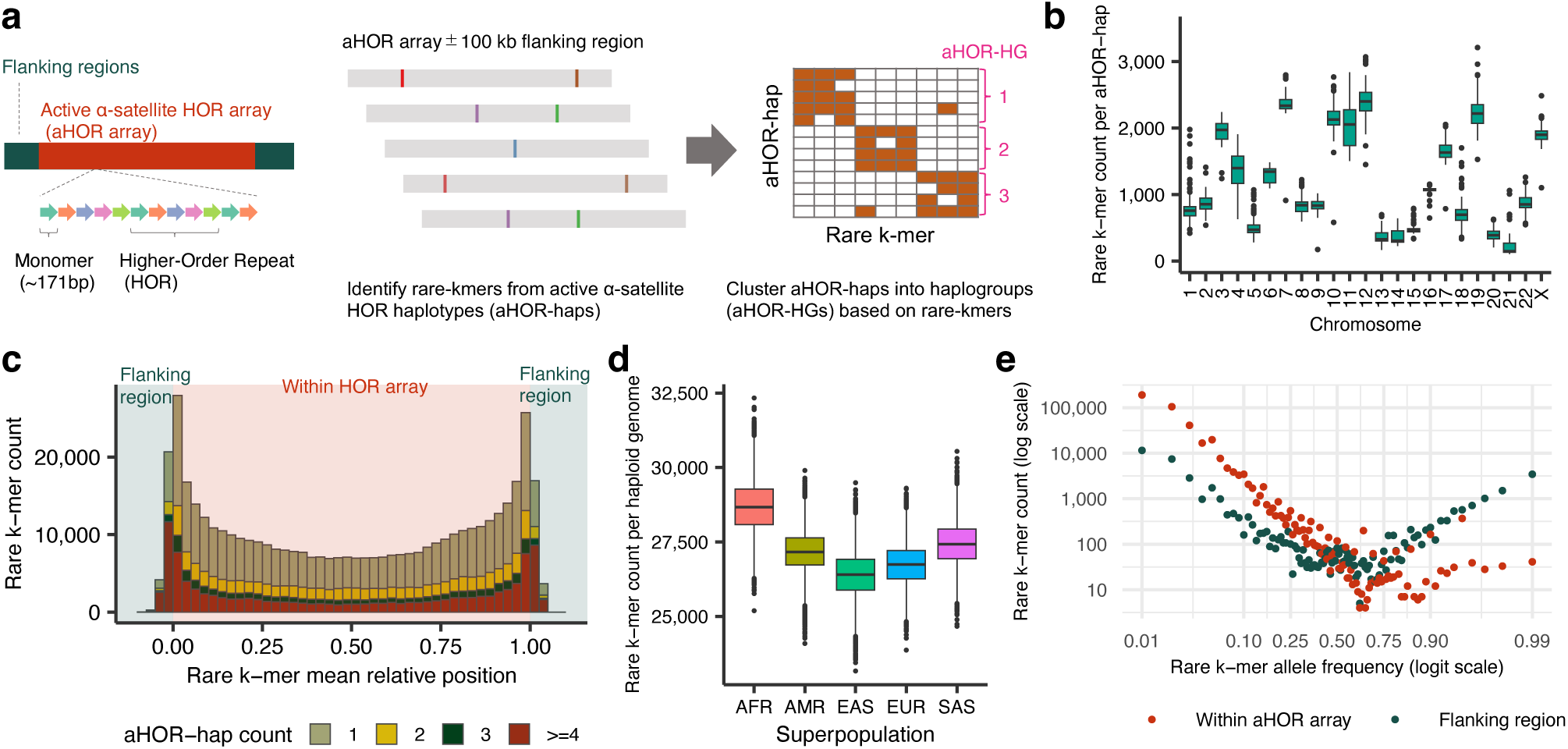
Study design and properties of rare k-mers from active α-satellite HOR haplotypes (aHOR-haps). (a) Schematic illustration of the active α-satellite higher-order repeat array (aHOR array), the identification of rare k-mers from aHOR-haps, and their clustering into haplogroups (aHOR-HGs) for downstream applications. (b) Distribution of rare k-mer frequencies per aHOR-hap across chromosomes. See also ED_Fig. 1b. (c) Number and mean relative position of rare k-mers aggregated across chromosomes, stratified by the number of aHOR-haps containing them. Mean relative position is defined as (rare k-mer position within aHOR array) / (aHOR array length); values between 0 and 1 indicate locations within aHOR arrays, whereas values outside this range correspond to flanking regions of the arrays. See also ED_Fig. 1c. (d) Estimated rare k-mer counts per haploid genome across superpopulations. Boxplots show the distribution of rare k-mer counts per haploid genome across individuals from five superpopulations in the 1000 Genomes Project: African (AFR), American (AMR), East Asian (EAS), European (EUR), and South Asian (SAS). For each superpopulation, 10,000 Monte Carlo iterations were performed. In each iteration, rare k-mer counts were sampled from the empirical allele frequency distribution of each chromosome and summed across all chromosomes to estimate the total count per haploid genome. See also ED_Fig. 2a. (e) Rare k-mer allele frequency spectra, aggregated across all chromosomes. Log-scaled scatter plots showing the number of rare k-mers (y-axis) as a function of their allele frequency (x-axis), stratified by the location of the rare k-mers (within the aHOR array or the flanking region). Allele frequencies were binned from 0.01 to 1.00 in 0.01 increments using ceiling-based binning. Red and gray points represent rare k-mers in aHOR arrays and flanking regions, respectively. See also ED_Fig. 2b.

Most rare k-mers within aHOR arrays had low allele frequency in the reference aHOR-hap panel, rarely exceeding 0.5. By contrast, many k-mers in the flanking regions exhibited very high allele frequencies (>0.90). These results suggest that rare k-mers in the flanking regions generally have evolutionarily older ancestral origins compared to those within aHOR arrays, which emerged in the human population more recently (Fig. 1e and ED_Fig. 2b). Some haplotype pairs shared many rare k-mers, while others had only a few or no such rare k-mers, suggesting that these aHOR-haps could be segregated into distinct groups according to their similarities (ED_Fig. 3a). In fact, using the hierarchical clustering based on the rare k-mer profiles, aHOR-haps in each chromosome were clustered into 3 to 16 (median: 8) haplogroups (aHOR-HGs) (Fig. 2a, b, ED_Fig. 3b-d, Supplementary Fig. 1, Supplementary Table 2 and Methods), where each haplogroup is denoted like *Cn*-aHOR-HG by assigning chromosome number (*n*).

**Figure 2:**
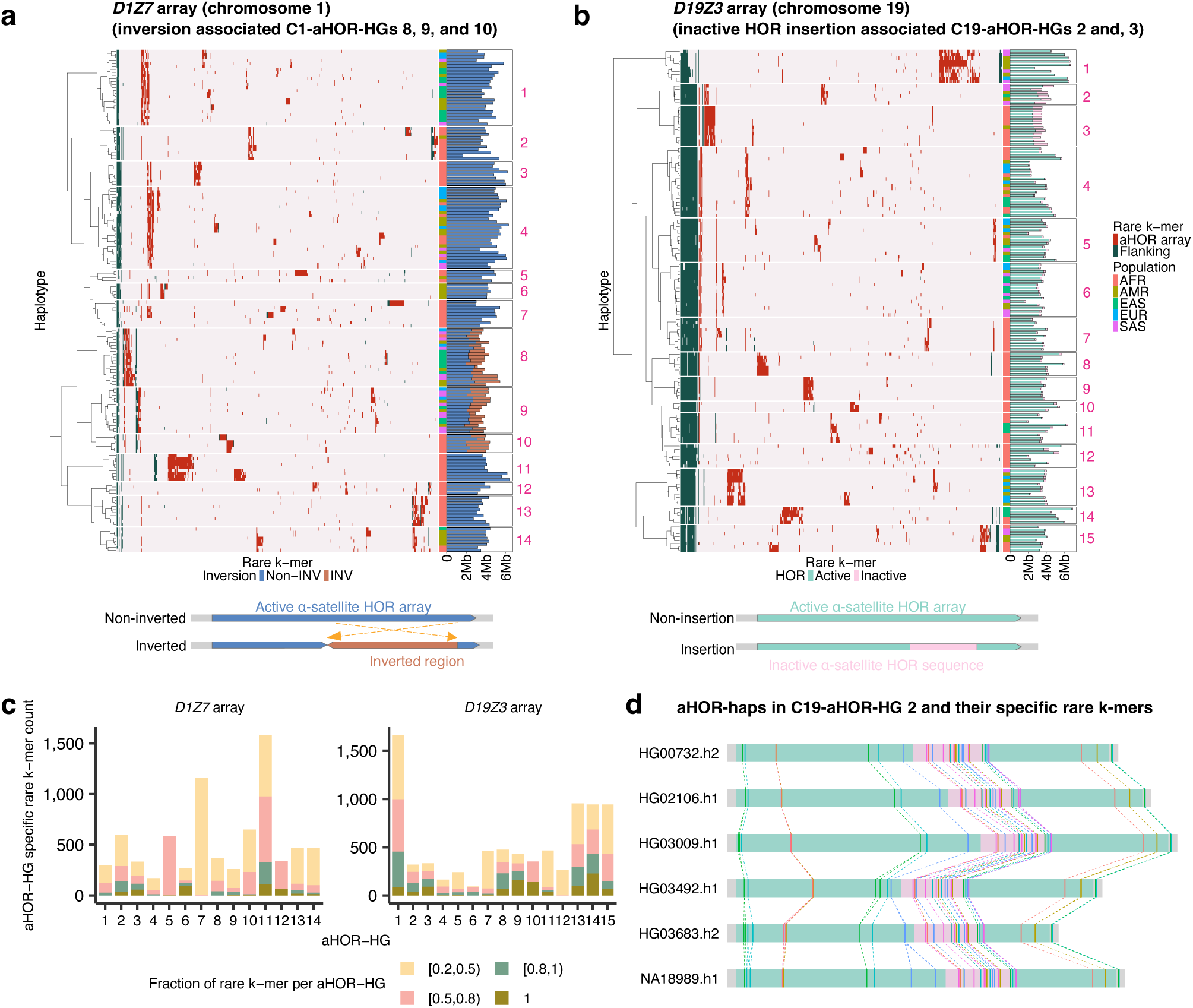
Clustering of active α-satellite HOR haplotypes (aHOR-haps) into haplogroups based on rare k-mers. (a, b) Heatmaps showing presence or absence profiles of rare k-mers across aHOR-haps for the *D1Z7* array (a) and *D19Z3* array (b). Haplotypes were clustered into haplogroups (aHOR-HGs) using hierarchical clustering. Rows represent individual aHOR-haplotypes, with superpopulation annotations shown as color-coded bars on the right. The adjacent bar plots indicate haplotype sizes, color-coded according to structural characteristics (e.g., inverted configurations or insertions of inactive α-satellite HOR sequences). Each heatmap column represents a rare k-mer marker, with colored tiles indicating presence in each haplotype. Rare k-mers located within the active α-satellite HOR array are shown in red (aHOR array), while those located in the flanking regions are shown in green (Flanking). Below each heatmap, schematic diagrams illustrate representative structural variations: inversions within *D1Z7* array and insertion of inactive α-satellite HOR sequences within *D19Z3* array. See also ED_Fig. 3d and Supplementary Fig. 1. (c) aHOR-HG-specific rare k-mer counts per haplogroup. Bar plots show the number of rare k-mers specifically observed in each aHOR haplogroup (aHOR-HG) on chromosome 1 (left) and chromosome 19 (right). Only rare k-mers detected in at least two aHOR-haps were included. Bars are stratified according to the fraction of aHOR-haps within each aHOR-HG that carry each rare k-mer: [0.2, 0.5) (yellow), [0.5, 0.8) (pink), [0.8, 1) (green), and 1 (brown). See also Supplementary Figs. 2 and 3. (d) aHOR-haps in C19-aHOR-HG 2 and their specific rare k-mers. Each horizontal track represents one of the five aHOR-haps assigned to the C19-aHOR-HG 2 haplogroup. Colored lines indicate the presence of rare k-mers uniquely associated with and shared by all aHOR-haps within this haplogroup. Each distinct color corresponds to a different rare k-mer, and connecting lines trace their presence across haplotypes. The shaded pink region marks inserted inactive α-satellite HOR sequences, which are a hallmark of the C19-aHOR-HG 2 haplogroup.

In accordance with their allele frequency distributions (Fig. 1e and ED_Fig. 2b), rare k-mers from the flanking regions were broadly shared by many aHOR-HGs in each chromosome, whereas those within the aHOR arrays were largely restricted to one or a few aHOR-HGs, which shared sets of unique rare k-mers in a hierarchical manner, reflecting their evolutionary history. Notably, the majority of aHOR-HGs (155 /207) were associated with unique sets of rare k-mers that were consistently shared by all haplotypes within the haplogroup, but undetected in any other haplogroups, suggesting that each aHOR-HG represents an evolutionarily distinct set of haplotypes (Fig. 2c, d and Supplementary Figs. 2 and 3). Supporting this is the observation that several haplotypes with distinct structural variations (SVs) or HOR profiles were co-segregated into particular aHOR-HGs, as exemplified in the aHOR arrays on chromosomes 1 (*D1Z7*), 7 (*D7Z1*), 11 (*D11Z1*), 17 (*D17Z1*), and 19 (*D19Z3*). A subset of *D1Z7* aHOR-haps is known to contain a large inversion in their long-arm side on the array^15^. We confirmed that 40 aHOR-haps with the inversion (range: 0.28–2.70 Mb; median: 1.44 Mb) in the reference aHOR-hap panel share identical junction sequences, indicating a common ancestral origin (ED_Fig. 4a). Consistent with this, these aHOR-haps clustered into three closely related haplogroups (C1-aHOR-HGs 8–10; Fig. 2a and ED_Fig. 4c, e, f). Similarly, 18 aHOR-haps in the *D19Z3* HOR array were assigned into two closely related aHOR-HGs (C19-aHOR-HGs 2 and 3; Fig. 2b and ED_Fig. 4d), all harboring large insertions (range: 0.40–1.06 Mb; median: 0.54 Mb) in the p-arm side of *D19Z3*^20^, primarily consisting of the sequences derived from *D19Z1*, an inactive α-satellite HOR array (Fig. 2b). All insertions in these aHOR-haps share identical junction sequences, indicating their common ancestral origin (ED_Fig. 4b). Furthermore, haplotypes within these two C19-aHOR-HGs were distinguished by the C19-aHOR-HG 2-specific monomeric HOR units adjacent to the *D19Z1*-derived sequences (ED_Fig. 4g, h). aHOR-haps in another aHOR array (*D17Z1*) contained multiple major HOR patterns, including 13-mer and two distinct 16-mer HORs (A and B)^29–31^, which were largely grouped into two sets of haplogroups, C17-aHOR-HG 1 and C17-aHOR-HGs 2–5. Haplotypes with 16-mer A HOR were mostly clustered into C17-aHOR-HG 1, whereas those in the remaining four C17-aHOR-HGs were characterized by 13-mer and the 16-mer B HORs (ED_Fig. 3d). Rare k-mer-based clustering also delineated additional characteristic HOR types, such as the atypical 12-mer HOR specific to C7-aHOR-HG 4 within *D7Z1*, where the predominant HOR pattern is the 6-mer HOR, and the atypical 6-mer HOR specific to C11-aHOR-HGs 6 and 7 within *D11Z1*, where the typical HOR pattern is the 5-mer^21^ (ED_Fig. 3d). Despite clustering based solely on the profiles of rare k-mers, without using information on SVs or HORs, we were able to distinguish the presence or absence of SVs and HORs, indicating that rare k-mer profiles alone effectively capture global structures and evolutionary lineages of aHOR-haps.

### Haplogroup typing using short-read sequencing

Based on the observation that aHOR-haps can be grouped into discrete haplogroups (aHOR-HGs) according to their rare k-mer profiles, we developed ascairn, a probabilistic framework that enables the inference of diploid aHOR-HG pairs (except for sex chromosomes in males) from short-read whole-genome sequencing (WGS) data for individuals outside the reference aHOR-hap panel. For each chromosome, we selected the aHOR-HG pair with the highest likelihood, where we assumed a two-step generative model for rare k-mer counts observed from short-read WGS data (Fig. 3a). To calculate this likelihood for each aHOR-HG pair, we first derived probabilities for each rare k-mer’s allele count (from 0 to 4) by convoluting empirical allele frequencies observed in individual aHOR-HGs. Next, for each rare k-mer, we computed the probabilities of observed counts in the short-read WGS by summing the products of negative binomial probabilities parameterized according to each allele count and their corresponding allele count probabilities obtained in the first step. Finally, the total likelihood was calculated by multiplying these probabilities across all rare k-mers. Furthermore, after determining the most probable aHOR-HG pairs, ascairn selects proxy aHOR-hap pairs from the reference aHOR-hap panel. These proxy pairs best approximate the given individual’s diploid aHOR-hap sequences and are chosen using a similar probabilistic model (Methods).

**Figure 3:**
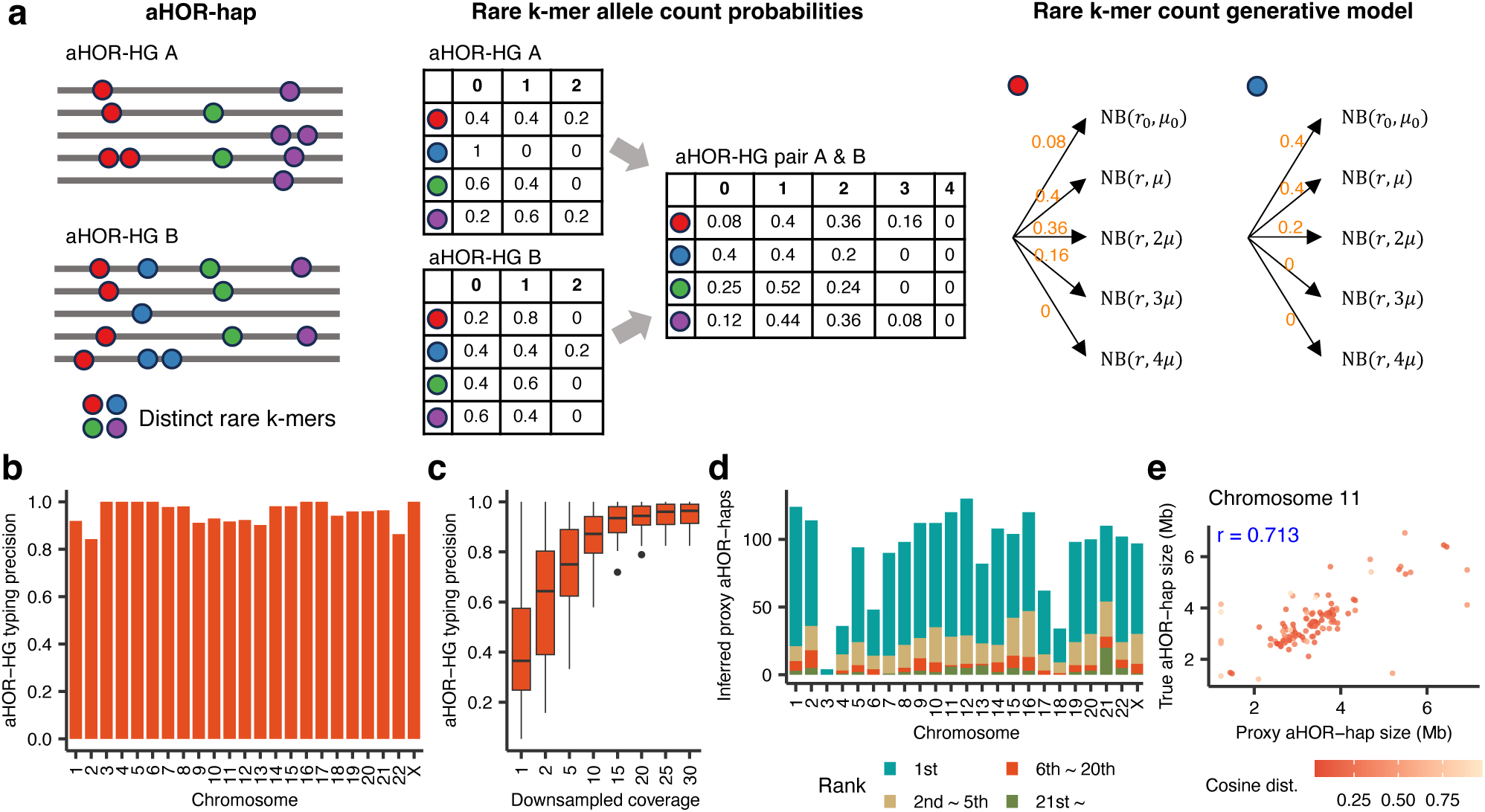
Probabilistic model underlying ascairn and its performance evaluation. (a) A schematic representation of the generative modeling process employed in ascairn for generating rare k-mer count profiles from short-read sequencing data. Rare k-mer frequencies were initially assessed within aHOR-haps for each haplogroup, and empirical probabilities for allele count distributions of each haplogroup were derived. These empirical probabilities were used to generate rare k-mer allele counts given specific haplogroup combinations. Rare k-mer counts in the short-read sequence data were assumed to be generated through a two-step process: first, allele counts of each rare k-mer in the corresponding individual were generated based on combined empirical probabilities, and second, observed counts of rare k-mers in the short-read sequencing data were generated according to the corresponding negative binomial distribution model. (b) Precision of ascairn haplogroup typing evaluated through leave-one-individual-out cross-validation for each chromosome. Clustering required a minimum of three haplotypes per haplogroup (see ED_Fig. 3b). See also ED_Fig. 5a. (c) Precision analysis at varying sequence coverage levels, illustrated by boxplots showing precision variability per chromosome following downsampling. See also ED_Fig. 5b. (d) Counts of haplotypes selected by ascairn via leave-one-individual-out cross-validation for each chromosome, stratified according to cosine distance ranking relative to the true haplotype. See also ED_Fig. 5c. (e) Accuracy of aHOR-hap size estimation from proxy aHOR-haps. Scatter plot showing the correlation between proxy-derived and true aHOR-hap sizes for chromosome 11, based on leave-one-individual-out cross-validation. Each point represents an individual aHOR-haplotype; point color denotes the cosine distance between the proxy and true aHOR-haps. Pearson’s correlation coefficient (*r*) is shown in the top left. See also ED_Fig. 5d.

The performance of ascairn was evaluated by a “leave-one-individual-out” cross-validation using haplotypes from the reference aHOR-hap panel in which fully phased diploid aHOR-hap data and matched short-read WGS data were available (Methods). The accuracy of aHOR-HG pair assignment ranged from 84.2% to 100% across chromosomes (median: 96.4%), and exceeded 90% for most chromosomes (Fig. 3b and Supplementary Table 3). Accuracy was lower for chromosomes having a large number of aHOR-HGs, such as chromosome 2, which was substantially improved when small aHOR-HGs were merged (ED_Fig. 3c and ED_Fig. 5a). Downsampling analysis suggested that ≥15× coverage was sufficient for reliable haplogroup typing (Fig. 3c and ED_Fig. 5b). We also validated the accuracy of proxy aHOR-hap pair inference using “leave-one-individual-out” cross-validation. The proportion of cases in which the proxy aHOR-hap closest to the actual haplotype (in terms of cosine distance) was correctly selected ranged from 50.9% to 84.4% (median: 74.0%). When considering the top five closest proxy aHOR-haps, this proportion increased, ranging from 74.6% to 98.9% (median: 91.9%) (Fig. 3d, ED_Fig. 5c, and Supplementary Table 4). We evaluated how well proxy aHOR-haps approximate true array sizes and observed a correlation coefficient ranging from 0.227 to 0.835 (median: 0.596) (Fig. 3e and ED_Fig. 5d). Restricting the analysis to proxy aHOR-haps with low cosine distance (≤ 0.3) to the true haplotypes (accounting for 49.7% of proxy aHOR-haps) markedly improved accuracy (correlation coefficient from 0.501 to 0.974 (median: 0.868)). Thus, array size can be reliably estimated using proxy haplotypes, particularly when closely related sequences (cosine distance ≤ 0.3) are available in the reference aHOR-hap panel.

In the following sections, we applied ascairn to large-scale publicly available short-read WGS datasets to investigate the variation of aHOR-HGs within human populations and its impact on the propensity for centromere-involving structural variants (SVs) in human cancers.

### aHOR-HG variations in large human populations

We first applied ascairn to two sets of short-read WGS data from the 1000 Genomes Project (*n* = 2,502) and the Human Genome Diversity Project (HGDP) (*n* = 828), enabling us to assess aHOR-HG variation across globally diverse human populations (Supplementary Table 5). The median number of individuals per population was 99 (range: 61–113) for the 1000 Genomes Project and 10.5 (range: 2–44) for the HGDP, across 26 and 54 populations, respectively. We observed marked differences in haplogroup frequencies within individual aHOR arrays (ED_Fig. 6a, and Supplementary Fig. 4). As suggested from the observations in the reference aHOR-hap panel, the inferred distributions of aHOR-HG substantially differed across populations (Fig. 4a and ED_Fig. 7). Many haplogroups exhibited pronounced population specificity. In particular, African populations notably displayed distinct haplogroup patterns compared to non-African populations: many haplogroups were found almost exclusively in African populations, with minimal or no representation in non-African groups (e.g., C1-aHOR-HG 10), while less common haplogroups were widely shared across all non-African populations but were present only at low frequencies in Africa (e.g. C1-aHOR-HGs 8 and 9). This asymmetry in haplogroup distribution, where African populations harbor many haplogroups not observed elsewhere, is in agreement with the Out-of-Africa model for the origin of modern human populations^32,33^. This interpretation is further supported by increased Shannon and Simpson diversity index (Fig. 4b and ED_Fig. 6b), and consistent with elevated rare k-mer complexity observed in African-derived haplotypes, pointing to both sequence-level and evolutionary diversity within these groups.

**Figure 4:**
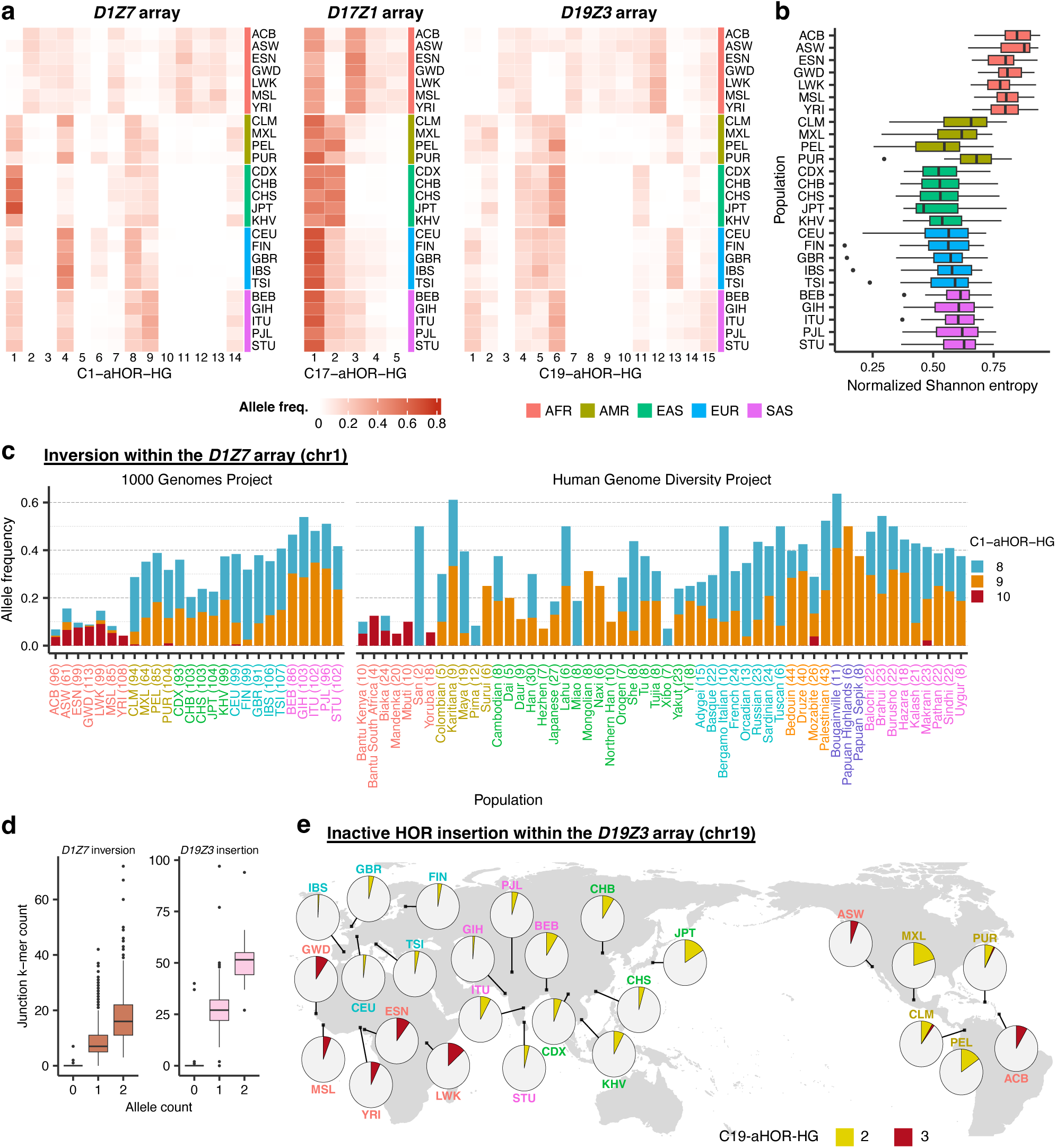
Variation of α-satellite HOR haplogroups (aHOR-HGs) across populations. (a) Allele frequencies of aHOR-HGs on chromosomes 1, 17 and 19 among populations included in the 1000 Genomes Project. Heatmap cell color intensity indicates allele frequency for each haplogroup within each population. Superpopulation annotations are displayed as colored bars on the right side of the heatmap. See also ED_Fig. 7. (b) Chromosome-wise haplogroup diversity per population measured by normalized Shannon entropy. Boxplots show the distribution of normalized Shannon entropy across chromosomes for each population in the 1000 Genomes Project. Entropy was calculated per chromosome and per population based on allele frequency distributions of haplogroup assignments estimated by ascairn. Only chromosomes with five or more haplogroups were included in the analysis. See also ED_Fig. 6b. (c) Allele frequencies of aHOR-HGs specific to the inversion within the *D1Z7* array across global populations from the 1000 Genomes Project and the Human Genome Diversity Project. Frequencies were calculated by summing the corresponding haplogroup allele frequencies, with bar colors indicating superpopulations. See also ED_Fig. 8a-c. (d) Boxplots showing the relationship between allele counts and junction k-mer counts for *D1Z7* inversions and *D19Z3* insertions. For *D1Z7* inversion events (left), junction k-mer counts were quantified by summing the occurrences of the 30 bp start and end junction sequences. For *D19Z3* insertions (right), junction k-mer counts were similarly derived from the number of occurrences of the 30 bp start and end junction sequences. The x-axis represents the number of alleles assigned to specific aHOR-HGs inferred by ascairn: for inversions, alleles assigned to C1-aHOR-HG 8, 9, or 10; for insertions, alleles assigned to C19-aHOR-HG 2 or 3. Boxplots display the distribution of total junction k-mer counts (y-axis) across samples, stratified by these allele counts. (e) Allele frequencies of aHOR-HGs specific to the inactive α-satellite HOR insertion within the *D19Z3* array (chromosome 19). Pie charts indicate proportions mapped to the geographic locations corresponding to each population.

The distribution of aHOR-HGs with common ancestors among different populations may provide insight into human evolutionary history (ED_Fig. 8a-c). To illustrate this, we first focused on the haplogroups, C1-aHOR-HGs 8-10 (Fig. 4c). As seen above, they are characterized in the reference aHOR-hap panel by large inversions with identical junction sequences, confirming that each of these haplogroups derived from a common ancestral haplotype. These junction sequences were highly specific to individuals assigned to C1-aHOR-HGs 8–10, detected in 1,649 of 1,660 such cases but in only 22/1,670 cases (mostly observed in only one read) without these haplogroups, supporting the high accuracy of ascairn inference (Fig. 4d). Despite their common ancestral origins, these haplogroups showed distinct distributions across different populations. C1-aHOR-HG 10 was almost exclusively found in African populations. By contrast, C1-aHOR-HGs 8 and 9 were also present in African populations but at very low frequency, whereas they were common in non-African populations. Notably, C1-aHOR-HG 8 was more prevalent in Europeans, whereas C1-aHOR-HG 9 predominated among South Asians. These findings suggest that the founder haplotype carrying the original inversion likely emerged and propagated among African populations, from which two groups of haplotypes (C1-aHOR-HGs 8 and 9) evolved and spread primarily outside the continent, while C1-aHOR-HG 10 was confined to African populations. Another set of haplogroups of interest was C19-aHOR-HGs 2 and 3. They were characterized by shared insertions of an inactive α-satellite HOR sequence, which was confirmed by the presence of unique junction sequences in all but four individuals assigned to these haplogroups (Fig. 4d). These aHOR-HGs also showed striking differences in population distributions. Of these haplogroups, C19-aHOR-HG 2 was most prevalent in Central and South American populations (including indigenous groups), followed by East Asian populations, whereas its frequency was notably low in Europeans and nearly absent in Africans (Fig. 4e and ED_Fig. 8a). In contrast, C19-aHOR-HG 3 was almost exclusively observed in African populations. These patterns support a model in which C19-aHOR-HG 2 diverged from C19-aHOR-HG 3 in the context of the Out-of-Africa migration and subsequently spread through East Asia and into the Americas via the Bering Strait.

### Analysis of 1p/19q co-deletions in gliomas

Translocations involving centromere regions are common in human cancer. Unlike other cancer translocations, they are poorly characterized at the molecular level primarily due to the highly repetitive nature of the centromere regions, although involvement of aHOR arrays has been implicated^4–7^. Among these is the unbalanced translocation der(1;19)(q10;p10), a class-defining abnormality in *IDH*-mutated oligodendroglioma (ODG) characterized by distinct clinicopathological and genetic features, resulting in 1p/19q co-deletion^22–24,34^. To investigate the molecular features of this translocation, we analyzed 142 ODG cases with der(1;19) from The Cancer Genome Atlas (TCGA), most of whom were of European ancestry. The *D1Z7* and *D19Z3* HOR arrays both belong to suprachromosomal family 1 (SF1), characterized by an almost perfectly alternating arrangement of two monomer types (J1 and J2) (Supplementary Fig. 5), and exhibit extremely high nucleotide sequence similarity, sharing largely identical monomers^2,15^. This may provide a structural basis for non-allelic homologous recombination, motivating us to explore putative breakpoints within these regions. To this end, we first applied ascairn to short-read WGS data from normal DNA samples to infer aHOR-HG and proxy aHOR-hap pairs for each patient, using the reference aHOR-hap panel for chromosomes 1 and 19. Proxy haplotype selection was restricted to haplotypes with available methylation data (Methods and Supplementary Table 6). Then, we inferred the proxy aHOR-hap corresponding to the rearranged allele and the putative breakpoint region. In this analysis, we first extracted allele-specific rare k-mers distinguishing the two proxy aHOR-haps. We then calculated copy numbers of these k-mers from tumor and normal short-read sequencing data. This approach enabled us to identify rearranged alleles exhibiting significant copy number shifts at the breakpoint (Fig. 5a; Methods). We could not apply this approach to cases in which the two proxy aHOR-haps belonged to the same aHOR-HG, as in that case, high similarity of both haplotypes precluded the identification of sufficient allele-specific rare k-mers for copy number analysis. Consequently, 104 and 119 cases were evaluable for copy number analysis of *D1Z7* and *D19Z3*, respectively. As expected, after excluding cases with suspected poor sequencing quality, breakpoints were mapped to the *D1Z7* and *D19Z3* arrays in all evaluable samples (ED_Fig. 9, Supplementary Table 7 and Supplementary Figs. 6 and 7). These findings strongly suggest that the breakpoints of 1p/19q co-deletion are located within active aHOR arrays on both participating chromosomes.

We next investigated the positions of rearrangement breakpoints relative to the functional centromeres, i.e., the kinetochore attachment sites, which correspond to locally hypomethylated regions known as centromere dip regions (CDRs) within the active aHOR arrays^15,35^. In this analysis, functional centromeres were inferred based on the CDRs of the proxy aHOR-haps. Given that the 1p/19q co-deletion is the characteristic consequence of this translocation, breakpoints are expected to occur on the p-arm side of the CDR in *D1Z7* and on the q-arm side in *D19Z3* (Fig. 5b). Although the expected pattern was observed in most cases (90/103 for *D1Z7* and 84/118 for *D19Z3*), a substantial fraction deviated from this trend (Fig. 5c). Specifically, the breakpoints were located on the q-arm side of the CDR in *D1Z7* in 7/103 cases and to the p-arm side of the CDR in *D19Z3* in 27/118 cases, suggesting that the CDR and/or breakpoint positions were inaccurately estimated in these instances. Notably, these cases were significantly enriched in C19-aHOR-HGs 1, 14, and 15 (Fig. 5c), which lack haplotypes from individuals of European ancestry. This observation suggests that the misestimation of breakpoint locations relative to the CDRs in these TCGA cases may be due to poor representation of population-specific haplotypes in these aHOR-HGs in terms of CDR locations and that these proxy aHOR-haps from these aHOR-HGs should be excluded from the further analyses involving CDR position estimation.

**Figure 5:**
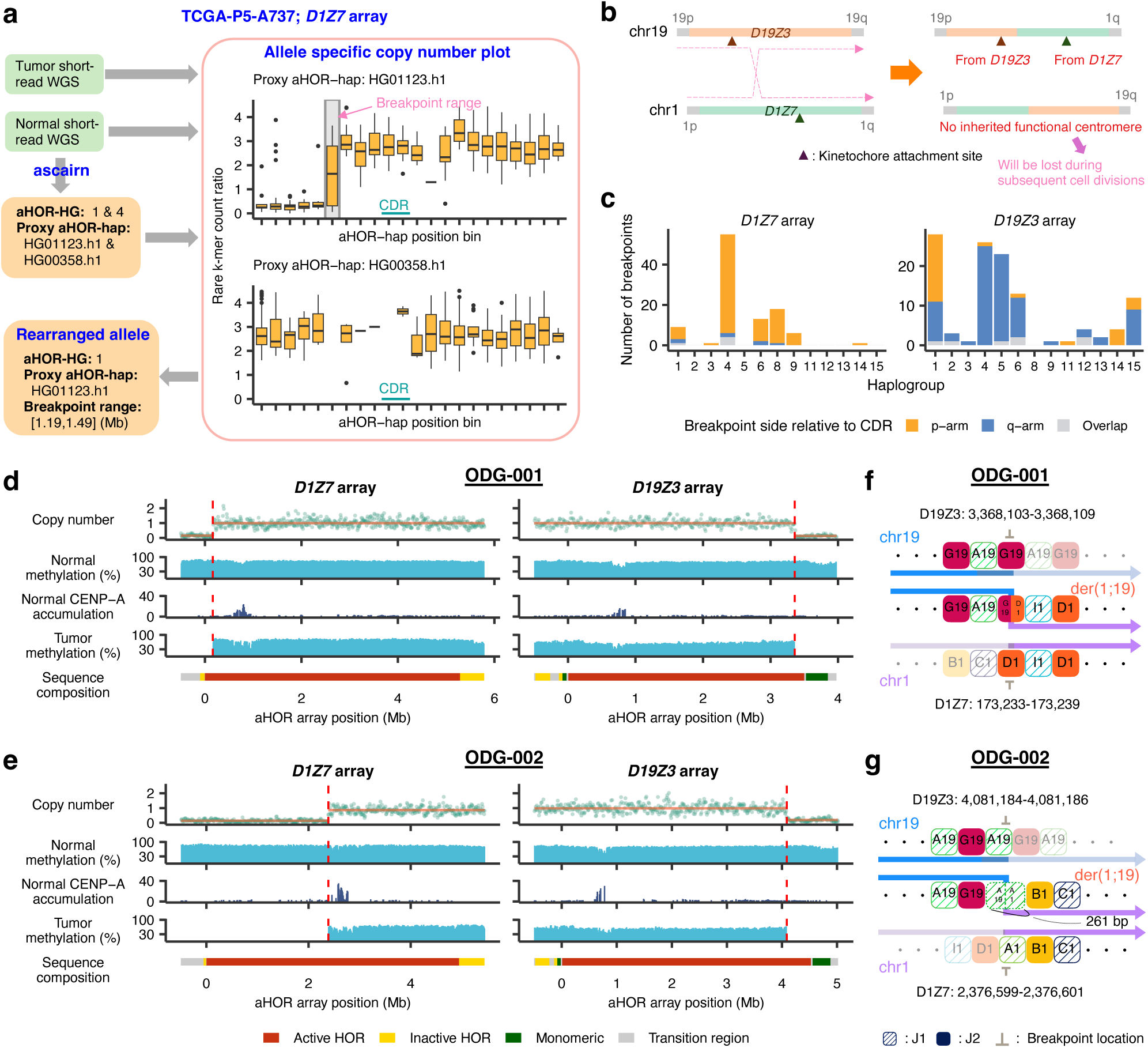
Rearrangement breakpoint mapping using short-read and long-read oligodendroglioma samples. (a) Illustrative example for inference of rearranged aHOR-haps (and –HGs) and breakpoint regions using TCGA-P5-A737, focusing on the *D19Z3* array. Using short-read whole-genome sequencing (WGS) data from normal DNA, ascairn was applied to infer the diploid C19-aHOR-HG pair (1 and 4) and to select two proxy aHOR-haps (HG01123.h1 and HG00358.h1). Based on allele-specific rare k-mers distinguishing the two proxy aHOR-haps, copy number profiles were computed from tumor and matched normal WGS data. Boxplots show the rare k-mer count ratio for each position bin along the proxy aHOR-haps. A distinct copy number gain was observed around the region (1.19–1.49 Mb) for HG01123.h1, indicating that it corresponds to the rearranged allele. (b) Schematic of the rearrangement leading to 1p/19q co-deletion. Following homologous recombination, two derivative chromosomes are transiently generated: one dicentric chromosome (a fusion of 1q and 19p) that retains functional centromeres from both parental chromosomes, and one acentric chromosome (a fusion of 1p and 19q) that is subsequently lost during cell division. (c) Distribution of inferred breakpoint regions relative to centromere dip regions (CDRs) across aHOR-HGs. Bar plots show the number of inferred breakpoint regions detected in each aHOR-HG (x-axis) for the *D1Z7* (top) and *D19Z3* (bottom) HOR arrays. Breakpoints are categorized based on the location of their inferred regions relative to the CDR: falling on the p-arm side (orange), the q-arm side (blue), or spanning both sides (overlap, grey). (d, e) Copy number profiles (normalized by ploidy), DNA methylation levels, and CENP-A concentrations at the *D1Z7* and *D19Z3* arrays in alleles harboring translocation breakpoints in (d) ODG-001 and (e) ODG-002. DNA methylation is presented separately for normal and tumor tissues. CENP-A concentrations are shown in normal tissues. Sequence compositions indicating active α-satellite HOR arrays (Active HOR; red), inactive α-satellite HOR array (Inactive HOR; yellow), monomeric α-satellite (Monomeric; green) and transition region (Transition region; gray) are provided below. See also Supplementary Fig. 10. (f, g) Structural characteristics of monomer sequences surrounding breakpoint regions in (f) ODG-001 and (g) ODG-002. Sequence homology between chromosomes at the breakpoints introduced ambiguity in determining precise breakpoint locations. The *D1Z7* and *D19Z3* α-satellite HOR arrays of chromosomes 1 and 19 belong to the same suprachromosomal family 1 (SF1), comprising distinct J1 and J2 monomer groups. Breakpoints occurring within the same monomer groups resulted in fusion monomers of typical size (∼170 bp) in ODG-001 and atypical size (261 bp) in ODG-002.

### Validation of breakpoint location using long-read sequencing

The findings regarding the der(1;19) breakpoints in the above section were directly validated using long read sequencing of matched tumor/normal DNA from two surgical specimens (ODG-001 and ODG-002) using Oxford Nanopore Technologies (ONT) and PacBio HiFi platforms supplemented with Hi-C sequencing (Supplementary Table 8). To identify the breakpoint sequences, we first performed *de novo* diploid assembly of sequence reads from normal samples using hifiasm, generating four centromere-containing contigs for chromosomes 1 and 19 in each of two cases, which was used as normal references (Supplementary Table 9). All contigs contained the active α-satellite HOR array sequences of the corresponding centromeres, including *D1Z7* and *D19Z3* sequences (Supplementary Table 10). These assemblies were validated by two methods.

First, they were compared for concordance with assemblies generated by another assembler, Verkko^16^. Second, uniform sequence coverage was confirmed by aligning raw sequence reads to the corresponding contigs (Methods and Supplementary Figs. 8 and 9). Based on the alignment of tumor and normal reads to the corresponding contigs, we evaluated genomic copy numbers on each contig in each tumor sample. This revealed the expected copy number shift (namely 1p/19q co-deletion) in one of the diploid aHOR arrays on each relevant chromosome, indicating that the translocation breakpoints occurred within these aHOR arrays (Fig. 5d, e and Supplementary Fig. 10). Finally, by aligning tumor reads containing specific sequence markers near breakpoint regions to each active α-satellite HOR sequence of corresponding normal samples (Methods), we successfully determined the breakpoint sequences at single-nucleotide resolution. In both cases examined, rearrangements occurred between two monomers from the same group (J2 in ODG-001 and J1 in ODG-002). Moreover, the rearrangement produced a fusion monomer of canonical size (∼171 bp) in ODG-001, whereas in ODG-002, it resulted in a non-canonical fusion monomer (261 bp) (Fig. 5f, g). Finally, we confirmed that the rearrangements actually occurred on the p-arm side of the CDR in *D1Z7* and the q-arm side of the CDR in *D19Z3*, generating a dicentric chromosome^36^ retaining two functional centromeres from chromosomes 1 and 19 (i.e., dic(1;19)(p10;q10)) (Fig. 5b). These observations collectively suggest that the rearrangements were triggered by high sequence homology between the two SF1-derived α-satellite arrays, which share similar monomer organization.

### Impact of centromere haplotypes on translocation

aHOR-haps displayed large structural diversity in array length and CDR composition (ED_Fig. 9). For example, in *D19Z3*, the CDRs of C19-aHOR-HG 13 haplotypes were highly polarized toward the q-arm side (ED_Fig. 10a). Notably, among 30 der(1;19)(+) tumors carrying these haplotypes, only two cases (6.7%) exhibited rearrangements involving C19-aHOR-HG 13, where the breakpoints were located within a very narrow segment between the CDR and the q-arm end of the array. Based on this observation, we hypothesized that aHOR-HGs influence the risk of translocation events through the size of breakpoint-susceptible regions (BSRs; regions defined as extending from the kinetochore toward the p-arm in *D1Z7* and toward the q-arm in *D19Z3*) (ED_Fig. 10b). We observed that BSR size displayed a substantial variability among aHOR-HGs (Fig. 6a and ED_Fig. 10c). We calculated the BSR size estimates for ODG samples using the array length of proxy aHOR-haps and the mean relative CDR position (range: 0–1.0) averaged across all aHOR-haps within the same haplogroup (aHOR-HG), thereby accounting for variability of CDR positions (Methods). In cross-validation, we confirmed that this provided robust estimates of the actual BSR size (R = 0.58 for *D1Z7* and 0.68 for *D19Z3*) (ED_Fig. 10d and Methods).

**Figure 6:**
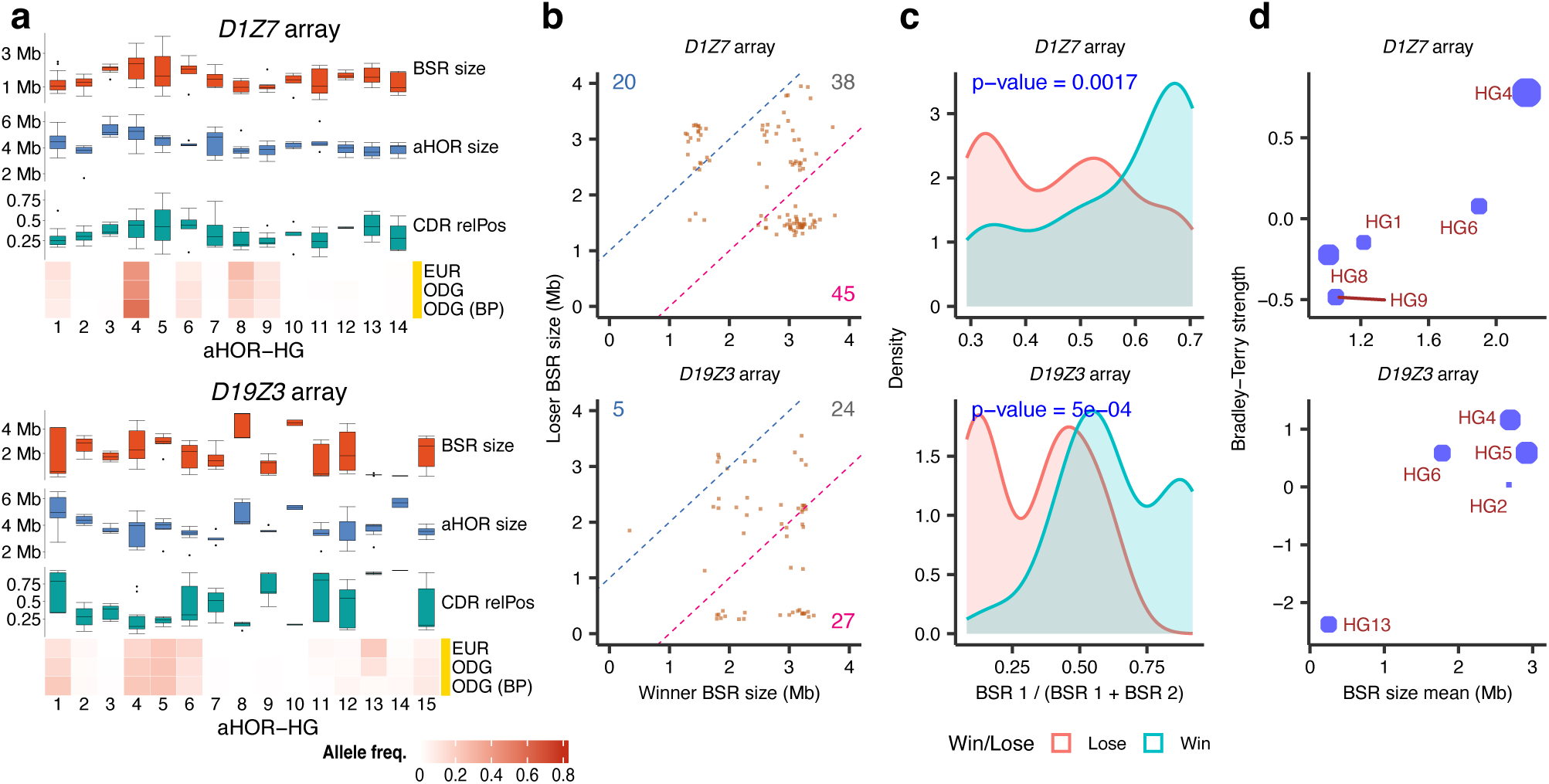
Variation in breakpoint-susceptible region (BSR) sizes in oligodendrogliomas. (a) Boxplots showing the distribution of breakpoint-susceptible region (BSR)-related statistics among aHOR-haps belonging to each haplogroup. These statistics include BSR sizes, total active α-satellite HOR array sizes, and relative positions of centromere dip regions (CDR). Data are presented separately for (top) D1Z7 α-satellite HOR arrays (chromosome 1), and (bottom) D19Z3 α-satellite HOR arrays (chromosome 19). Below each set of boxplots, heatmaps depict allele frequencies of aHOR-HGs for all analyzed ODG cases, ODG alleles restricted to those containing breakpoints, and the European population (EUR) from the 1000 Genomes Project, serving as a control group for ODG cases. (b) Scatter plots comparing the BSR sizes of aHOR-hap pairs in tumors with 1p/19q co-deletion, separately shown for (top) D1Z7 and (bottom) D19Z3 α-satellite arrays. Each point represents a tumor sample, with the x-axis and y-axis indicating the BSR sizes of the haplotype involved in the rearrangement (Winner) and the other parental haplotype (Loser), respectively. Dashed lines represent ±1 Mb difference between the two haplotypes. Numbers indicate sample counts in each region. (c) Density plots illustrating the distributions of breakpoint-susceptible region (BSR) ratios for D1Z7 (top) and D19Z3 (bottom). The BSR ratio is defined as BSR1 divided by the sum of BSR1 (first allele) and BSR2 (second allele). Samples are stratified into “Win” (blue) or “Lose” (red) groups based on the outcome for the first allele. Statistical significance of the association between BSR ratio and group outcomes was assessed by logistic regression, and corresponding p-values are shown in blue. (d) Scatter plots presenting the relationship between haplogroup mean BSR size (in megabases, Mb) and rearrangement propensity (estimated as selection strength) using the Bradley-Terry model based on pairwise comparisons between aHOR-HG pairs. Each circle represents aHOR-HG, with circle size proportional for allele frequency for haplogroup. Data are presented separately for (top) D1Z7 and (bottom) D19Z3 α-satellite HOR arrays.

We then assessed whether larger BSR sizes were associated with increased rearrangement frequency. Within each patient, the parental haplotype with the larger BSR size was significantly more likely to undergo rearrangement than the haplotype with the smaller BSR size for both *D1Z7* (*P-*value = 0.002) and *D19Z3* (*P-*value = 1 × 10⁻⁴) (Fig. 6b, c). Next, we estimated the relative propensity for rearrangement across aHOR-HGs using the Bradley-Terry model^37^ and examined its correlation with the mean BSR size (Methods). This analysis revealed substantial differences in rearrangement propensity among aHOR-HGs, which significantly correlated with their average BSR sizes in *D1Z7* and *D19Z3* (Fig. 6d). As expected, C19-aHOR-HG 13 in *D19Z3*, which has CDRs strongly polarized toward the q-arm side and therefore short BSRs, exhibited the lowest rearrangement propensity. In contrast, C1-aHOR-HG 4 in *D1Z7*, with CDRs polarized toward the q-arm side and larger BSRs, showed a high propensity for rearrangement. These results suggest that the propensity of rearrangement between *D1Z7* and *D19Z3* varied across aHOR-HGs depending on their average BSR size.

## Discussion

We showed that rare k-mers can serve as effective markers for dissecting the complexity of centromere structure, particularly that of aHOR arrays, across human populations and for understanding centromere-involving abnormalities commonly found in human cancer^38^. The extensive repetitive nature of centromere sequences, especially of aHOR arrays, has long prevented identification of such markers through pairwise alignment of haplotype sequences. Thus, in previous studies, analyses of centromere haplotype variation have primarily relied on pericentromeric sequences flanking these arrays, which are more amenable to alignment and marker identification^10,15,21^. In this study, by contrast, we identified rare k-mer markers from within aHOR arrays using a fully alignment-free approach, enabling the analysis of centromere diversity from standard short-read sequencing data.

One of the major findings enabled by rare k-mer markers is the identification of distinct haplogroups of aHOR-haps. They were defined based on similarity in rare k-mer profiles, were organized into a hierarchical structure, were often associated with distinct SVs and HORs, were maintained by the non-recombining nature of centromeric arrays, and exhibited population-specific distributions. Thus, aHOR-HGs capture the lineage structure of aHOR-haps in a manner analogous to mitochondrial and Y-chromosome haplogroups^39–41^, offering a coherent framework for investigating centromere diversity and its biological relevance. Although the existence of centromeric haplogroups was previously inferred from variation in flanking pericentromeric regions, such inference remained indirect and lacked resolution within the core α-satellite HOR arrays themselves. In contrast, our rare k-mer-based approach would enable direct haplogroup definition within these otherwise inaccessible regions, providing a more accurate and sequence-resolved view of centromere diversity.

Another key finding was the significant correlation between der(1;19) translocations and the estimated size of the BSR within the *D1Z7* and *D19Z3* arrays, with longer BSRs conferring higher rearrangement risk. Centromeres are recurrent hotspots for structural abnormalities in cancer and other diseases^5^, implicated in aneuploidy and allelic imbalance. However, the role of centromere structure in their pathogenesis has remained largely unknown. Using rare k-mer-based analysis, the current study therefore provides, to our knowledge, the first evidence that structural polymorphisms of aHOR arrays can modulate susceptibility to disease-specific centromeric abnormalities.

Finally, we note an important limitation of the current rare k-mer-based framework. In some cases, ascairn failed to correctly assign aHOR-HG types or identify sufficiently close proxy aHOR-haps. This limitation is likely due to the relatively small number of reference aHOR-haps (∼100 per chromosome) currently available for rare k-mer extraction. As the number of fully resolved centromere haplotypes continues to grow through advances in long-read sequencing^18,20,42,43^, incorporating these additional sequences will be critical for improving the resolution and accuracy of haplogroup classification. This expansion will enhance our understanding of population-level centromere diversity, its evolutionary dynamics, and its contribution to human disease.

## Methods

### Collection of active α-satellite HOR haplotypes and rare k-mers

#### Extracting active α-satellite HOR haplotypes from genome assemblies

We downloaded chromosome-scale genome assemblies from several pangenome consortia. For the Human Pangenome Reference Consortium (HPRC)^18^, we downloaded Year 1 data from the following URL: s3://human-pangenomics/working/HPRC/{sample}/assemblies/year1_f1_assembly_v2_genbank/{sample}.{pater nal|maternal}.f1_assembly_v2_genbank.fa.gz. We also downloaded assemblies obtained in the study of hifiasm (UL)^17^, available at: s3://human-pangenomics/submissions/53FEE631-4264-4627-8FB6-09D7364F4D3B--ASM-COMP/{sample}/assemblies/hifiasm_v0.19.5/trio/{sample}.{pat or mat}.fa.gz. From the Human Genome Structural Variation Consortium (HGSVC) Phase 3^20^, we downloaded genome assemblies generated via Verkko from the following FTP site: https://ftp.1000genomes.ebi.ac.uk/vol1/ftp/data_collections/HGSVC3/working/. From the T2T Consortium, we used CHM13 v2.0^14^ and HG002 v1.1 (available at https://s3-us-west-2.amazonaws.com/human-pangenomics/T2T/HG002/assemblies/hg002v1.1.fasta.gz). We also used the CHM1 genome assembly^20^, available at https://ftp.ncbi.nlm.nih.gov/genomes/all/GCA/037/575/895/GCA_037575895.1_UW_CHM1_v1.0/GCA_037575895.1_UW_CHM1_v1.0_genomic.fna.gz. If there were duplicated samples, we retained only one assembly, prioritizing them in the order T2T > HGSVC > hifiasm (UL) study > HPRC. Additionally, for trio samples, we excluded the child’s assembly if assemblies for both parental samples (paternal and maternal) were available.

We aligned these curated assemblies to the CHM13 v2.0 genome using minimap2^44^ with the “-x asm5” option and selected the contig sequences covering the active α-satellite HOR array (aHOR array) of each chromosome. The definition of the aHOR array for each chromosome was obtained from s3://human-pangenomics/T2T/CHM13/assemblies/annotation/chm13v2.0_censat_v2.1.bed, from which we selected regions whose HOR names included ‘H1L’.

Next, to define the aHOR array regions within each contig sequence, we applied StringDecomposer^45^ to each contig using monomer sequences corresponding to each chromosome, as defined in the HORmon^46^ paper (the Monomers Final directory at https://figshare.com/articles/dataset/HORmon/16755097/1). We first identified partial aHOR array regions, defined as regions composed of monomers with ≥90% sequence identity that extended continuously for at least 1,000 bp. The aHOR array region for each contig was then defined by merging these partial regions into a single genomic span, extending from the start of the first partial region to the end of the last partial region.

Next, we extracted assembly contigs that fully encompassed the aHOR arrays defined above, extending each extracted region by a margin of 500 kbp upstream and downstream. For chromosomes 3 and 4, in which active α-satellite HOR regions typically span two distinct areas^15,21^, the margin size was extended to 5 Mbp. Finally, for each extracted contig, the sequence of the aHOR array region, along with 100 kbp of upstream and downstream flanking sequences, was designated as the active α-satellite HOR haplotypes (aHOR-haps).

aHOR-haps were excluded when both paternal and maternal aHOR-haps were available for the same sample and exhibited high similarity in size, with a normalized length difference below 0.5%, to avoid potential assembly artifacts. Furthermore, when aHOR-haps from samples with familial relationships exhibited high size similarity, one of the samples was excluded to avoid redundancy.

#### QC check of active α-satellite HOR haplotypes

To identify mis-assemblies within the aHOR-haps, we used NucFlag (v0.2.4) (https://github.com/logsdon-lab/NucFlag), a fork of NucFreq^47^. PacBio HiFi sequencing data corresponding to each assembly were downloaded and aligned to their respective genome assemblies using minimap2 with the “-ax map-hifi” option. The alignments were then sorted, indexed, and filtered to remove secondary and supplementary alignments. Next, for each aHOR array region corresponding to each aHOR-hap, NucFlag was executed with the default configuration. aHOR-haps meeting either of the following criteria were considered likely mis-assemblies and excluded from further analysis: (1) at least one flagged region (COLLAPSE, COLLAPSE_VAR, or MIS_JOIN) exceeding 100 kb in size was present, or (2) four or more flagged regions ranging from 100 bp to 100 kb in size were detected. The remaining aHOR-haps constitute the reference aHOR-hap panel. We also performed HumAS-HMMER (https://github.com/fedorrik/HumAS-HMMER_for_AnVIL) for a more detailed investigation of centromere sequence composition where necessary.

#### Gathering of rare k-mers

Considering robustness against sequencing errors, adequate discriminatory power, and observability in short-read sequencing, we employed a k-mer size of 27. Rare k-mers were extracted separately for each chromosome. First, for each aHOR-hap on a chromosome, we enumerated all canonical k-mers and stored them in a dictionary. Here, a canonical k-mer was defined as the lexicographically smaller sequence between each k-mer and its reverse complement. We then merged the lists of k-mers and their occurrence counts for each aHOR-hap, and extracted those occurring at most twice within an aHOR-hap of a given chromosome as rare k-mers. We referred to this initial collection as the “raw rare k-mer set”.

We observed that rare k-mers often occurred at consecutive genomic positions within an aHOR-hap. These consecutive rare k-mers typically represent the same underlying variant, as sliding windows of size k overlapping the variant each tend to be identified as a rare k-mer. We aggregated these essentially redundant rare k-mers. Specifically, we scanned each aHOR-hap sequence again, merging consecutive rare k-mers into groups if they were registered in the raw rare k-mer set extracted above. Then, for each group, we calculated hash values for rare k-mers within the group, selected the k-mer with the smallest hash as the representative, and removed all others. Hash values were computed using the built-in hash function of Python 3, with the environment variable PYTHONHASHSEED set to 42. Furthermore, rare k-mers appearing across multiple chromosomes were removed.

As a final filtering step, we compared the allele counts of rare k-mers in aHOR-haps against their actual observation counts from short-read sequencing, and removed rare k-mers exhibiting substantial discrepancies. For each chromosome, we first selected samples in which both paternal and maternal aHOR-haps were available. Among these, we identified samples with short-read sequencing data available from the 1000 Genomes Project (s3://1000genomes/1000G_2504_high_coverage). For each sample, sequencing reads mapped to the active α-satellite HOR array (defined in the “Definition of active α-satellite HOR arrays of GRCh38” section) were extracted, and occurrences of rare k-mers within these sequence reads were counted.

For each rare k-mer, we first classified samples based on the allele count (AC) of the rare k-mer in their aHOR-haps:

● AC = 0 group: samples in which neither the paternal nor maternal aHOR-haps contained the given rare k-mer.
● AC ≥ 1 group: samples in which at least one of the paternal or maternal aHOR-haps contained one or more occurrences of the rare k-mer.

Then, rare k-mers were filtered out if they failed to meet either of the following criteria:

● Among samples in the AC = 0 group, the proportion of samples containing 0, 1, or 2 occurrences in short-read sequencing was 90% or more.
● Among samples in the AC ≥ 1 group, the proportion of samples containing three or more occurrences in short-read sequencing was 90% or more.

#### Estimating rare k-mer counts per haploid genome

To estimate the number of rare k-mers per haploid genome (from chromosome 1 to chromosome X), we performed Monte Carlo simulations. First, based on the number of rare k-mers observed in each aHOR-hap within the reference panel (each annotated with the superpopulation of the individual from whom it was derived), we constructed empirical distributions of rare k-mer counts for each chromosome and superpopulation (AFR [African], AMR [American], EAS [East Asian], EUR [European], SAS [South Asian]). Then, for each superpopulation, we sampled rare k-mer counts from the corresponding chromosome-specific empirical distributions (chromosomes 1 through X) and summed them to obtain the total rare k-mer count per haploid genome. This procedure was repeated 10,000 times for each superpopulation.

### Haplogrouping of active α-satellite HOR haplotypes

First, we constructed a data matrix in which the rows represent aHOR-haps and the columns represent rare k-mers, with each element indicating the occurrence frequency (0, 1, or 2) of a given rare k-mer in a specific aHOR-hap. Rare k-mers observed in only a single aHOR-hap were excluded. Next, we computed a similarity matrix (S) from this matrix using cosine similarity, which was then converted into a distance matrix (D) using the formula D = 1 − S.

Hierarchical clustering was performed using Ward’s method (“ward.D2”), which minimizes within-haplogroup variance. To determine the optimal number of clusters (haplogroups), we used the silhouette score as the evaluation criterion. Specifically, the number of haplogroups was varied from 3 to 20, and the average silhouette score was calculated for each clustering result. Additionally, to ensure that each haplogroup contained a sufficient number of aHOR-haps, we imposed a requirement that each haplogroup should contain at least three aHOR-haps (or five for evaluating the coarse classification). The final optimal number of haplogroups was determined as the value that maximized the average silhouette score while satisfying the minimum haplogroup size requirement.

### Inference of diploid active α-satellite HOR haplogroups and haplotypes using short-read sequencing data (ascairn)

#### Definition of active α-satellite HOR arrays of GRCh38

We first downloaded RepeatMasker annotations (rmsk.txt.gz) for the human genome assembly hg38 from the UCSC Genome Browser database (https://hgdownload.soe.ucsc.edu/goldenPath/hg38/database/rmsk.txt.gz). Alpha-satellite repeat regions were extracted from these annotations by selecting repeats annotated as “ALR/Alpha” with lengths greater than 1,000 bp. Overlapping or adjacent regions separated by less than 500,000 bp were merged using BEDTools (bedtools merge). Regions located on unplaced contigs, random sequences, fixed patches, or chromosome Y were excluded. Only merged active α-satellite HOR arrays exceeding 1 Mb in length were retained. Additionally, to define extended α-satellite HOR arrays, the boundaries of these curated regions were expanded by 500,000 bp both upstream and downstream.

#### Sequence coverage estimation for each sample

In this study, we selected the q-arm of chromosome 22 (the region starting from q11.21; chr22:17,400,001–50,818,468) as the reference region for evaluating sequence coverage. Sequence reads mapped to this selected region were extracted using the samtools view command (v1.21), after which mosdepth (v0.3.9) was executed to calculate coverage. Finally, from the mosdepth output, we extracted the row where the “chrom” column was labeled “total_region” and defined the corresponding value in the “mean” as the coverage.

#### Rare k-mer count profile generation

First, sequence reads aligned to the predefined active α-satellite HOR array regions of GRCh38 were extracted using the samtools view command, and then converted to FASTA format with samtools fasta. Next, rare k-mers were counted using the jellyfish (v2.3.0) “count” command with the options “-s 100M –C –m 27 –t 8 –-if rare_kmer_file”, where rare_kmer_file is the file path which contains the list of rare k-mers. Finally, the resulting counts were converted into TSV format, with the first column representing the rare k-mer nucleotides and the second column indicating their corresponding counts.

#### Typing active α-satellite HOR haplogroup pairs

For each haplogroup, we calculated the relative frequency of each rare k-mer (0, 1, or 2) within the set of aHOR-haps belonging to that haplogroup, adding a pseudo-count of 0.1. For each pair of haplogroups (including pairs consisting of the same haplogroup), we computed the convolution of their relative frequencies (0, 1, 2, 3 or 4).

We assumed the generative probabilities of rare k-mers follow a two-step model: first, the allele count (AC) of each rare k-mer is generated based on the convoluted relative frequencies; second, given this allele count, the observed count of each rare k-mer is generated from a negative binomial distribution (NBin) as follows:

– AC = 0: NBin(*r*_0_, μ_0_),

– AC = 1: NBin(*r*, μ),

– AC = 2: NBin(*r*, 2μ),

– AC = 3: NBin(*r*, 3μ),

– AC = 4: NBin(*r*, 4μ).

Here, the size parameters *r*_!_, *r* were set to 0.5 and 8, respectively. The mean parameter μ_!_ was set to 0.8, and the unit of mean parameter μ was set to 0.4 × *d* where *d* is the sequence coverage estimated in the previous subsection. Finally, we calculated the log-likelihood for each haplogroup pair, defined as the sum of the log probabilities across all rare k-mer occurrences, and selected the pair that maximized this log-likelihood.

#### Typing active α-satellite HOR haplotype pairs

Next, we performed typing of aHOR-hap pair. The process is fundamentally similar to the haplogroup pair typing. We first calculated the relative frequencies of rare k-mer allele counts for each aHOR-hap. Then, we computed the convoluted relative frequency for each aHOR-hap pair. Following the same two-step modeling approach used for haplogroup pairs, we selected the aHOR-hap pair that maximized the log-likelihood.

One point to note is that, for calculating the relative frequencies of allele counts for each aHOR-hap, we used a mixture of the relative frequencies from each aHOR-hap and their respective haplogroups, with mixing proportions of 0.9 and 0.1, respectively. All subsequent procedures followed the methodology previously described for haplogroup pairs.

### Validation of ascairn via cross-validation

For each chromosome, we selected individuals for whom both paternal and maternal centromeres were represented in the reference aHOR-hap panel and for whom corresponding short-read sequencing data were available at s3://1000genomes/1000G_2504_high_coverage/. To evaluate the performance of our method, we conducted leave-one-individual-out cross-validation, where for validation of each individual, we excluded its paternal and maternal aHOR-haps from the reference aHOR-hap panel and constructed a generative model of the rare k-mer count profile. Based on this model, ascairn was performed to infer the diploid aHOR-HG and aHOR-hap pair. For aHOR-HG, a prediction was considered correct only when both inferred haplogroups matched the true pair.

For downsampling experiments, we first calculated the average read depth on the q-arm of chromosome 22 from the sample’s short-read data. We then randomly subsampled reads via the “samtools view –-subsample” command to reach the target sequence coverage, and applied ascairn to the downsampled data.

For evaluating proxy aHOR-hap inference accuracy, we leveraged the fact that the ground-truth aHOR-haps for each individual were known. We computed cosine distances based on the rare k-mer count profile between the true aHOR-haps and all other aHOR-haps in the reference aHOR-hap panel, and assessed (i) how closely the predicted aHOR-hap ranked in proximity to the true one within the reference aHOR-hap panel, and (ii) the difference in aHOR-hap size.

### Analysis of α-satellite HOR haplogroup variations across human populations

#### Performing ascairn to short-read whole-genome sequencing data from large human populations

We applied ascairn to the short-read whole-genome sequencing data from the 1000 Genomes Project (available at s3://1000genomes/1000G_2504_high_coverage/data), and the Human Genome Diversity Project (HGDP) (available at https://ddbj.nig.ac.jp/public/public-human-genomes/GRCh38/HGDP/CRAM/). Two samples in the 1000 Genomes Project were excluded due to suspected file corruption. Population metadata were downloaded from https://www.internationalgenome.org/data-portal/data-collection/30x-grch38 and https://www.internationalgenome.org/data-portal/data-collection/hgdp, respectively. For each population, we calculated allele frequencies of aHOR-HGs.

#### Quantifying variation in α-satellite HOR haplogroups

For each chromosome and each population, we calculated the normalized Shannon entropy (to account for differences in aHOR-HG counts across chromosomes) and Simpson’s index based on the allele frequencies of aHOR-HGs. Chromosomes with four or fewer aHOR-HGs were excluded from the analysis. We then plotted the distributions of normalized Shannon entropy and Simpson’s index values.

### Generation of sequencing data for centromere rearrangement analysis

#### ONT Sequencing (ultralong and standard)

Genomic DNA was extracted from normal blood samples using the Monarch® HMW DNA Extraction Kit for Tissue (NEB, T3060) starting with 5 or 6 million cells. Libraries for these samples were then directly prepared using the ONT Ultra-Long DNA Sequencing Kit V14 (Oxford Nanopore Technologies, SQK-ULK114) following the standard protocol. For brain tumor samples, genomic DNA was extracted from tissue pieces using the same kit. Libraries for brain tumor samples were prepared from 100∼200 ng of the extracted genomic DNA using the ONT Rapid Sequencing Kit (SQK-RAD114).

Sequencing was performed on PromethION24 (PRO-SEQ024) using PromethION R10.4.1 flow cells (FLO-PRO114M) for both sample types. For normal blood samples, each library was typically loaded in quarters across two or three flow cells, with a flow cell wash step using the Flow Cell Wash Kit (Oxford Nanopore Technologies, EXP-WSH004) performed between loadings on the same flow cell. For brain tumor samples, a total of four libraries were analyzed using three flow cells, with one flow cell being used twice with a wash step between the loading of the two libraries.

Basecalling was performed using Guppy (v6.4.8) or Dorado (v0.6.0) with 5mC CpG methylation in super-accuracy mode, using the configuration file specific to each flow cell.

#### PacBio HiFi sequencing

Genomic DNA was fragmented using the Megaruptor 3 shearing kit (Diagenode E07010003) with a shear speed of 31. Fragment size distribution was assessed using the FemtoPulse System (Agilent Technologies, M5330AA) with the Genomic DNA 165 kb Kit (Agilent Technologies, FP-1002-0275), and the fragmentation step was repeated if necessary to achieve the desired size range. Fragmented DNA was then used for library preparation with the SMRTbell prep kit 3.0 (PacBio 102-182-700). This process involved purification with SMRTbell cleanup beads, followed by DNA damage repair, end repair, and dA-tailing. SMRTbell adapters were then ligated to the DNA ends, and the resulting constructs were cleaned up using SMRTbell cleanup beads. DNA molecules without SMRTbell adapters were digested with nuclease, and the remaining DNA was purified with AMPure PB beads (PacBio 102-182-500), targeting the elimination of fragments shorter than 5 kb. Sequencing primers were annealed, and polymerase was bound to the SMRTbell templates using the Sequel II Binding Kit 3.2.

The prepared libraries were sequenced on the Sequel IIe System (PacBio 101-986-400) using the Sequel II sequencing kit 2.0 (PacBio 101-802-200 or 102-194-400) and the SMRT Cell 8M tray (PacBio 101-389-001 or 102-281-700). Sequencing runs were managed using the instrument’s control software SMRT-Link v11.0.0 or v11.0.1. All PacBio HiFi data were generated by CCS v6.4.0 with the “--hifi-kinetics” option except for the ODG-002 sample, which was generated using CCS v6.3.0, and 5mC of CpGs in each HiFi read was subsequently predicted by Primrose (v1.3.0).

#### Hi-C sequencing

Hi-C experiments were performed using the MboI restriction enzyme, as previously described^48,49^. Briefly, two million cells were crosslinked with 1% formaldehyde for 10 minutes at room temperature. Cells were then permeabilized, and chromatin was digested with MboI. The resulting DNA fragment ends were labeled with biotinylated nucleotides and subsequently ligated. After reversing the crosslinks, DNA was purified and sheared using Covaris M220. Ligation junctions were enriched using streptavidin beads. Sequencing libraries were prepared using the NEBNext Ultra DNA Library Prep Kit for Illumina according to the manufacturer’s instructions and sequenced on an Illumina NovaSeq 6000 using standard 150-bp paired-end reads.

### Somatic translocation analysis on fully assembled α-satellite HOR sequences

#### Phased diploid genome assembly of matched-control samples

We generated phased diploid genome assemblies using hifiasm (version v0.19.6-r595) with the –-ul, –-h1, and –-h2 options enabled to incorporate ONT-UL and Hi-C sequencing data. Additionally, we used Verkko (version 2.0) for comparative validation. We then extracted aHOR-haps following the procedure described in the subsection “Extracting active α-satellite HOR haplotypes from genome assemblies” for chromosomes 1 and 19.

We also aligned the tumor/normal ONT and PacBio HiFi sequence data (unaligned BAM files with MM and ML tags) to the diploid genome assembly using the samtools fastq –T MM,ML command, piped into minimap2 with the option “-ax map-ont –y” (for ONT) or “-ax map-hifi –y” (for PacBio HiFi), to ensure that the MM and ML tags were retained in the aligned BAM files.

#### Somatic copy number analysis on active α-satellite HOR arrays

To perform copy number analysis, we first computed the sequencing depth of PacBio HiFi tumor and normal samples within aHOR arrays using the samtools depth command with the options “-F 2308 –q 1”. The depth values were segmented into 10,000 bp bins, and tumor/normal coverage ratios were calculated. Change-point detection was then applied to these ratios using the “cpt.mean” function from the “changepoint” R package (with the arguments method = “PELT”, penalty = “MBIC”, and minseglen = 20), and bins showing significant changes were designated as changepoint bins. We subsequently estimated ploidy within the aHOR arrays and corrected the tumor/normal coverage ratios based on the total number of sequenced bases in each sample. The resulting normalized coverage profiles were then visualized.

#### Methylation analysis on active α-satellite HOR arrays

For ONT tumor and normal sequencing data, we used modbam2bed (https://github.com/epi2me-labs/modbam2bed) with the options “--aggregate–-cpg–e” to obtain aggregated counts of methylated and unmethylated bases at each CpG site. The genome was then partitioned into 10,000 bp bins. For each bin, the total counts of methylated and unmethylated CpG bases were summed, and the methylation ratio was calculated and plotted on the aHOR arrays.

#### CENP-A binding region analysis on active α-satellite HOR arrays

CENP-A ChIP and input control paired-end sequencing data were independently aligned using BWA-MEM (v0.7.18) with the phased diploid genome assemblies as reference sequences. The resulting BAM files were filtered using samtools view with the options “-F 3332 –q 1”. The ratio of CENP-A ChIP to input control depth in the aHOR arrays was calculated using deepTools (v3.5.6) with the options “--smoothLength 30000 –-skipNAs –-binSize 10000”.

#### Identification of single-base resolution rearrangement breakpoints

We used the approximate breakpoint locations on *D1Z7* and *D19Z3* arrays inferred from somatic copy number analysis on aHOR arrays. First, unique 27-mer sequences, defined as sequences occurring exactly once within the four aHOR haplotypes of chromosomes 1 and 19, were identified. Next, chromosome-specific “breakpoint-associated unique 27-mer” sequences were extracted from genomic regions spanning 0.1 Mb upstream and downstream of the preliminary approximate positions. This process yielded two distinct sets of breakpoint-associated unique 27-mers for *D1Z7* and *D19Z3* arrays.

Tumor-derived PacBio HiFi reads containing breakpoint-associated unique 27-mer sequences from both *D1Z7* and *D19Z3* sets were selected. Within each of these reads, the unique 27-mer closest to the estimated breakpoint region (the most distal toward the p-arm side for *D1Z7* arrays and toward the long-arm side for *D19Z3* arrays) was designated as the breakpoint marker. For refined breakpoint localization, subsequences of the HiFi read between the two breakpoint markers (length L bp) were extracted. Artificial chimeric reference sequences were created, each consisting of two segments: (a) i bp extending toward the p-arm side from the *D1Z7* arrays breakpoint marker on the corresponding aHOR-hap, and (b) (L – i) bp extending toward the long-arm side from the *D19Z3* breakpoint marker on its respective haplotype. By systematically varying i from 1 bp to L bp and performing pairwise alignments with the extracted HiFi subsequences using edlib^50^, we defined the optimal breakpoint position as the one yielding the smallest total edit distance across the two alignments. Finally, consistency and reproducibility of the breakpoint positions were confirmed through repeated analyses across multiple independent tumor-derived HiFi reads meeting the above criteria.

### Localization of centromere dip region (CDR)

We collected BAM files with MM and ML tags from the Human Pangenome Reference Consortium (basecalled using Guppy versions 6.3.7–6.5.7, available at s3://human-pangenomics) and from the Human Genome Structural Variation Consortium (available from the 20240816_JAX_ONT_guppy6_Rebasecalled directory at ftp.1000genomes.ebi.ac.uk). These BAM files were first converted to FASTQ format using the samtools fastq command with “-T MM,ML” option. Subsequently, the FASTQ files were aligned to their respective genome assemblies for each sample using minimap2 with “-ax map-ont –y” option. The resulting alignment files were converted to BAM format (via samtools view with “-F 2304 –q 1” option), sorted, and indexed. Finally, BAM files restricted to regions defined as aHOR haplotypes were extracted for further analysis.

We then performed modbam2bed (https://github.com/epi2me-labs/modbam2bed) with the options “--aggregate –-cpg –e” to obtain aggregated counts of methylated and non-methylated bases at each CpG site. These regions were subsequently divided into bins of 10,000 bp. For each bin, total counts of methylated and non-methylated bases at CpG sites were summed, and the methylation ratio was calculated. A rolling mean was obtained using the function “rollapply” of the R package “zoo” with the arguments “width = 3L, FUN = mean, align = ‘center’, partial = TRUE”.

Finally, the mean and standard deviation of methylation ratios were calculated, and regions exhibiting methylation ratios lower than two standard deviations below the mean were designated as CDR regions.

### Detecting rearrangement breakpoint using tumor and matched control short-read sequencing data

We ran ascairn on the matched control samples to determine haplogroup pairs as well as the proxy aHOR-hap pairs. The selection of proxy aHOR-haps was restricted to those with CDR information, due to the absence of methylation information in some ONT sequencing data. Based on the hierarchical model used in this analysis, we calculated posterior probabilities for the allele counts for the two haplogroups for each rare k-mer. Next, we defined allele-specific rare k-mers as follows:

– Allele 1-specific rare k-mers: rare k-mers with posterior probability ≥ 0.8 when summing over the allele count combinations (1, 0) and (2, 0).
– Allele 2-specific rare k-mers: rare k-mers with posterior probability ≥ 0.8 when summing over the allele count combinations (0, 1) and (0, 2).

For each allele-specific rare k-mer, we tallied the observed counts in tumor and normal samples and calculated their tumor/normal ratios, limiting to k-mers with ≥8 counts in the normal sample. To identify breakpoints at the haplogroup level, we divided each rare k-mer into 20 bins across the aHOR array, as well as upstream and downstream bins, based on their positions on the proxy aHOR-hap. We then generated boxplots of tumor/normal count ratios of rare k-mers within each bin, and regions exhibiting abrupt changes were manually curated.

#### Sample selection for oligodendroglioma

We retrieved clinical data for the Brain Lower Grade Glioma TCGA PanCancer study from cBioPortal (https://www.cbioportal.org/) and identified patient IDs where the “Subtype” column was set to “LCC_IDHmut-cdel”. For these patients, we downloaded the corresponding tumor and matched control BAM files from the GDC Data Portal (https://portal.gdc.cancer.gov/), selecting those with “Experimental Strategy” set to “WGS” and “WGS Coverage” set to either “10x–25x”, “25x–150x”, or “unknown”.

### Inference of breakpoint-susceptible region size for each α-satellite haplotype

The breakpoint-susceptible region (BSR) was defined as the region extending from the left boundary of the CDR toward the p-arm end in *D1Z7* arrays, and from the right boundary of the CDR toward the q-arm end in *D19Z3* arrays. We first calculated the BSR size for each aHOR-hap in the reference aHOR-hap panel of chromosomes 1 and 19. We also calculated the average BSR size for each C1-aHOR-HG and C19-aHOR-HG.

To infer BSR size outside the reference aHOR-hap panel, we first applied ascairn to obtain the aHOR-HG and proxy aHOR-hap pairs. To reduce the estimation variance, we used a shrinkage-like approach^51^. Instead of using the BSR size of the proxy aHOR-hap directly, we estimated the final BSR size as the average of the proxy aHOR-hap BSR size and the mean BSR size of the corresponding aHOR-HG, in order to stabilize individual estimates.

### Evaluation of susceptibility to rearrangement acquisition for each haplogroup using Bradley-Terry model

For chromosomes 1 and 19, we constructed a star chart comparing rearrangement breakpoint susceptibility among aHOR-HGs, using results from the “Detecting rearrangement breakpoint using tumor and matched-control short-read sequencing data” section. For each sample, we scored two distinct aHOR-HGs (pairs with identical aHOR-HGs were excluded): the aHOR-HG with a breakpoint was labeled as the “winner”, the other as the “loser”. Results from all samples were combined into a comprehensive star chart. Then, we estimated the allele-selection strength of each haplogroup for the rearrangement using the “BTm” function in BradleyTerry2 R package^52^.

## Data availability

The raw sequence data acquired in this study will be available through public sequence repository service.

## Code availability

The software of ascairn is available at GitHub (https://github.com/friend1ws/ascairn).

## Acknowledgment

This work was supported by Grants-in-Aid from the Japan Agency for Medical Research and Development (AMED; JP24ama221538 to Y.Sh., JP24tk0124003h0002 and JP25ama221530h0002 to S.O., JP25ck0106019h0001 to Y.Oc.); National Cancer Center Research and Development Funds (2025-A-03 to Y.Sh.); the Moonshot Research and Development Program (24zf0127009h0003 to S.O.); the Ministry of Education, Culture, Sports, Science and Technology of Japan (MEXT; hp200138, hp210167, and JPMXP1020200102 to S.O.); Japan Society for the Promotion of Science (JSPS) KAKENHI (JP24H00009 to S.O. and JP24K19223 to Y.Oc.); the Research Support Project for Life Science and Drug Discovery (Basis for Supporting Innovative Drug Discovery and Life Science Research (BINDS)) from AMED (JP24ama121021 to T.T. and T.Y.); Japan Science and Technology Agency (JST) Fusion Oriented REsearch for disruptive Science and Technology (FOREST) Program (JPMJFR220L to Y.Oc.); and the National Institutes of Health (NIH; GM147352 to G.A.L.). We used the NIG supercomputer (provided by ROIS National Institute of Genetics) and Shirokane supercomputer (provided by Human Genome Center, the Institute of Medical Science, The University of Tokyo). The results shown here are partly based on data generated by TCGA Research Network (https://cancergenome.nih.gov/). The authors thank S.Tarumoto, M. Shinagawa, N. Sakamoto and staff at the Single-Cell Genome Information Analysis Core (SignAC) at WPI-ASHBi, Kyoto University for the long-read sequence analyses. The authors also thank Dr. Kozo Tanaka and Dr. Tatsuo Fukagawa for helpful discussions. We used ChatGPT (OpenAI) to assist in language editing and proofreading of the manuscript. We would like to acknowledge the Human Pangenome Reference Consortium (BioProject ID: PRJNA730823) and its funder, the National Human Genome Research Institute (NHGRI), and also acknowledge the Human Genome Structural Variation Consortium, funded by NHGRI in 2019.

## Author contributions

Y.Sh. and S.O. conceived the study and jointly led the project. Y.Sh., with assistance from M.S., Y.K., E.G., G.A.L. and S.O., designed and implemented ascairn and performed centromere variation analyses. Y.Sh., M.S. and Y.Sa., with support from A.O., Ha.S., K.C., Y.I., W.N., R.N.M. and M.K., performed cancer rearrangement analyses using long-read sequencing data. Y.Oc., K.K., T.T., and R.O., with assistance from F.O., K.M., T.Y., R.S. and S.O., collected samples, performed sequencing, and conducted wet-lab experiments. Y.Sh., M.S. and S.N. with assistance from A.O, T.M. and Hi.S. and Y.Ok. performed cancer rearrangement analyses using short-read sequencing data with ascairn. Y.Sh. and M.S. generated figures. Y.Sh. and S.O. wrote the manuscript.

---

**Extended Data Figure 1:**
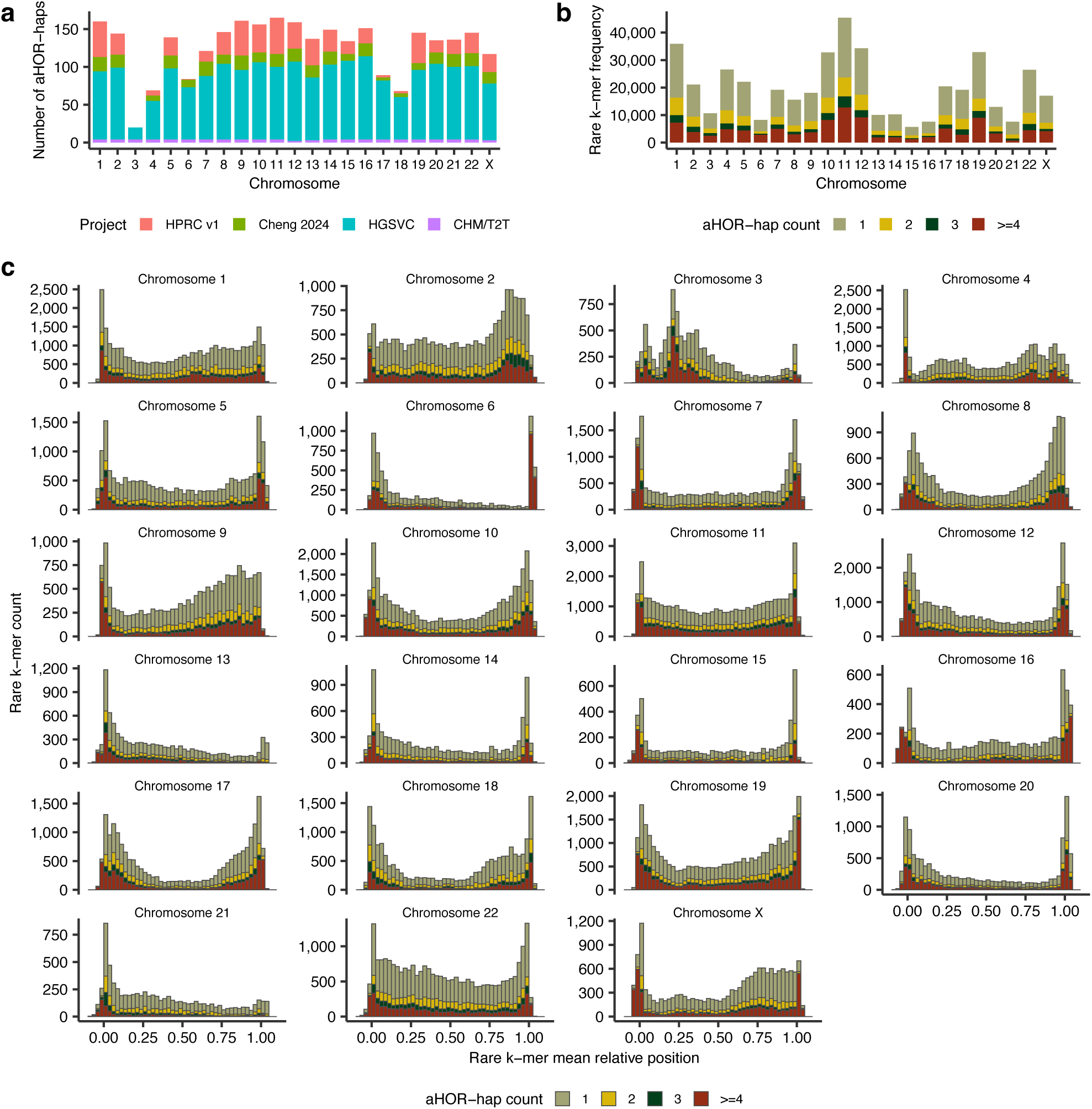
Statistics of active α-satellite HOR haplotypes (aHOR-haps) and rare k-mers. (a) Number of aHOR-haps obtained across chromosomes, stratified by the source genome project (HPRC v1, Cheng et al., 2024, HGSVC and CHM/T2T). (b) The number of rare k-mers detected across chromosomes, stratified by the number of aHOR-haps containing those k-mers. (c) The number and mean relative position of rare k-mers across chromosomes, stratified by the number of aHOR-haps in which they are present. The relative position is defined as (rare k-mer position within the aHOR array) / (aHOR array length); values between 0 and 1 indicate locations within the array, whereas values outside this range correspond to positions outside the array. See also Fig. 1c.

**Extended Data Figure 2:**
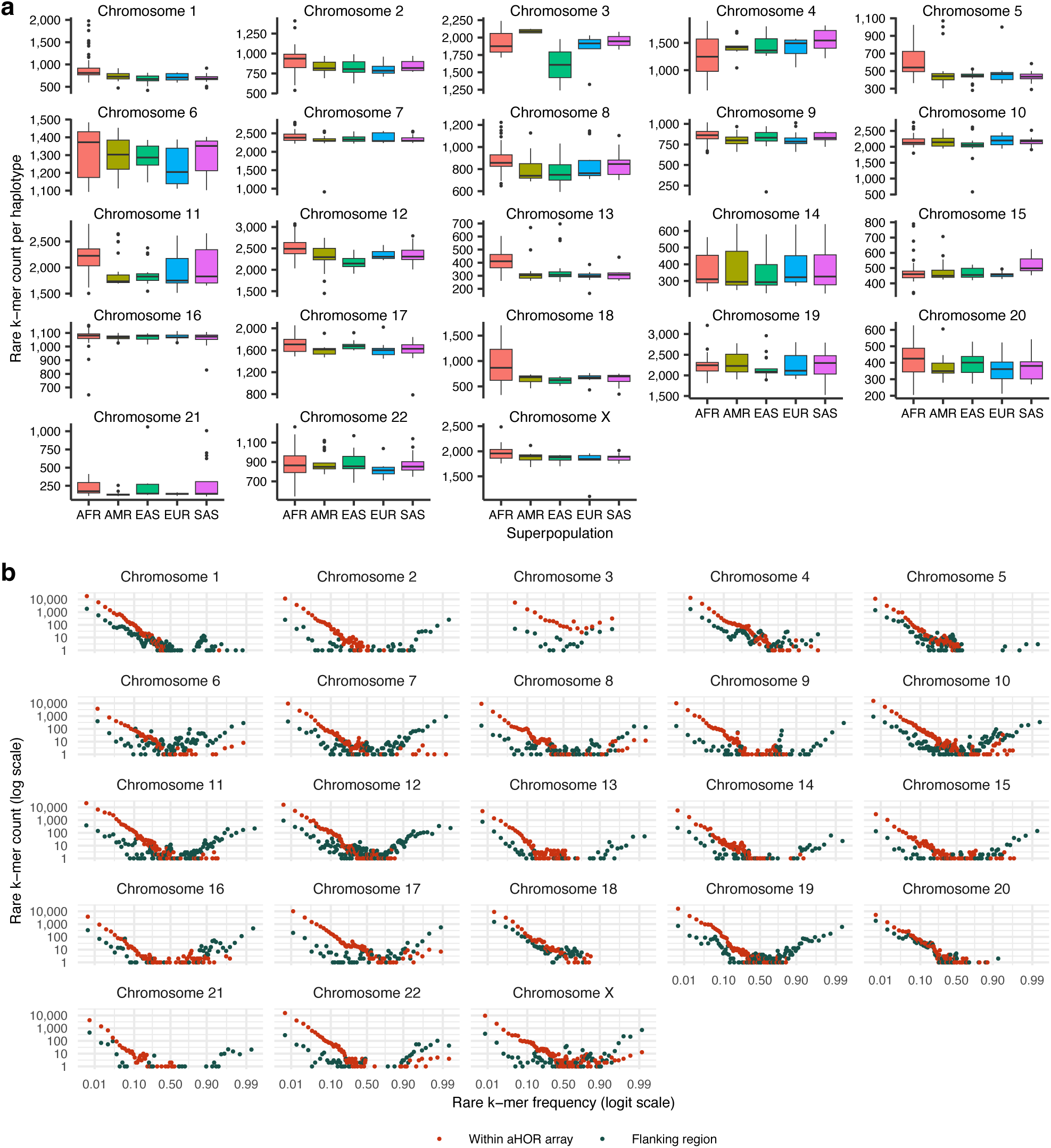
Rare k-mer counts across superpopulations and rare k-mer allele frequency spectra across all chromosomes. (a) Chromosome-wise distribution of observed rare k-mer counts per haplotype across superpopulations. Boxplots show the distribution of rare k-mer counts per chromosomal haplotype across five superpopulations from the 1000 Genomes Project: African (AFR), American (AMR), East Asian (EAS), European (EUR), and South Asian (SAS). Rare k-mer counts were directly calculated for each haplotype based on the observed presence or absence of rare k-mers on each chromosome. See also Fig. 1d. (b) Rare k-mer allele frequency spectra across all chromosomes. Log-scaled scatter plots show the number of rare k-mers (y-axis) as a function of their allele frequency (x-axis), stratified by the location of the rare k-mers (within the aHOR array or the flanking region). Red and gray points represent rare k-mers in aHOR arrays and flanking regions, respectively. See also Fig. 1e.

**Extended Data Figure 3:**
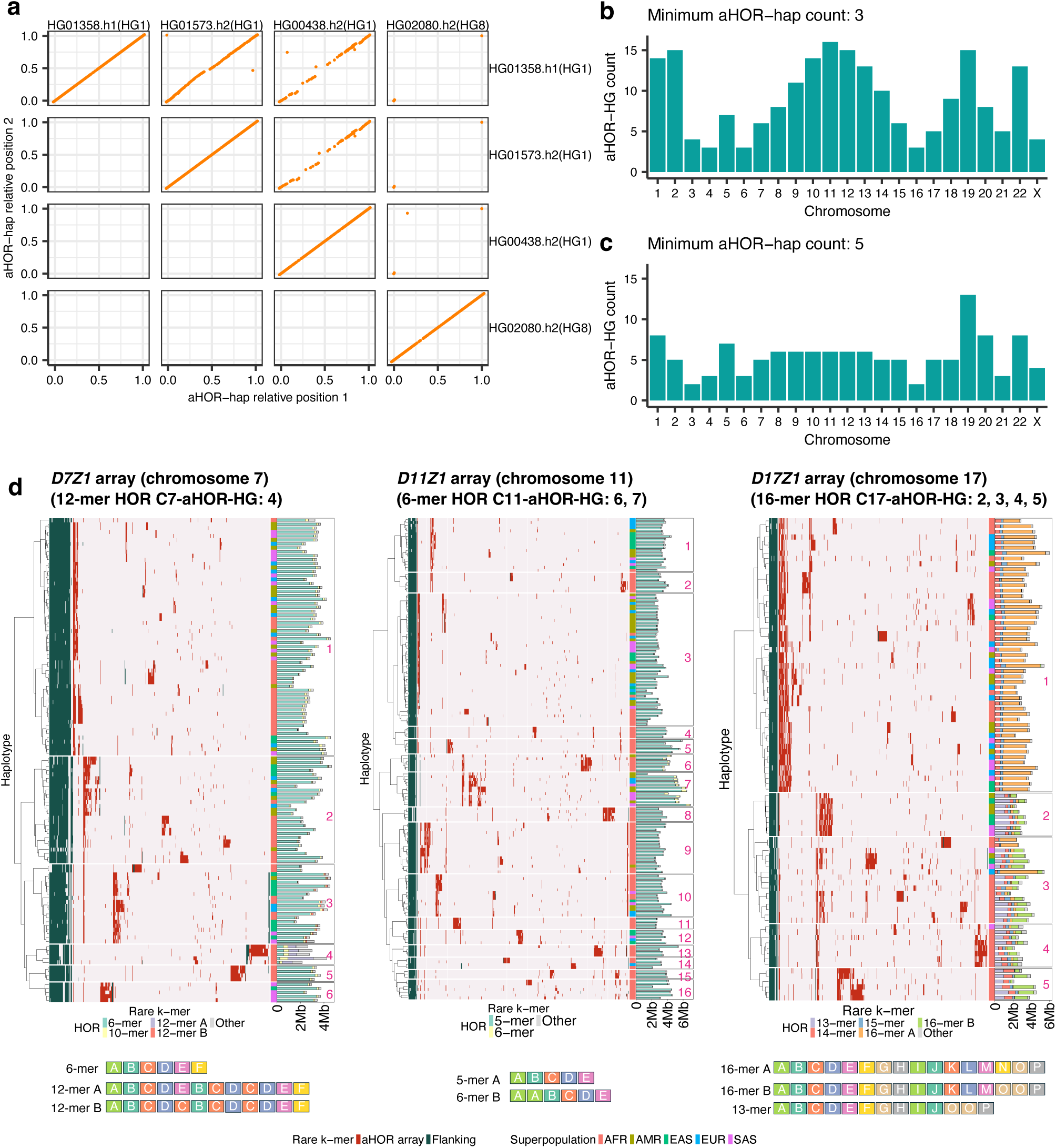
Haplogroup clustering and representative structural features for aHOR arrays at chromosomes 7, 11, and 17. (a) Relative positions of rare k-mers shared between pairs of *D1Z7* aHOR-haps. HG01358.h1, HG01573.h2, and HG00438.h2 (classified into C1-aHOR-HG 1), and HG02080.h2 (classified into C1-aHOR-HG 8) are plotted. Closely related aHOR-hap pairs shared numerous rare k-mers aligned along the diagonal, whereas distantly related pairs shared very few. (b) Number of aHOR-HGs in each chromosome, in which the constraint that each aHOR-HG contains at least three aHOR-haps was imposed. This clustering is mainly used in this study. (c) Number of aHOR-HGs in a different setting, in which the constraint that each aHOR-HG contains at least five aHOR-haps was imposed. (d) Heatmaps showing rare k-mer presence (red: within aHOR array; dark green: flanking region) profiles across aHOR-haps of the *D7Z1*, *D11Z1* and *D17Z1* arrays (chromosomes 7, 11, and 17, respectively). aHOR-haps were grouped into aHOR-HGs using hierarchical clustering. Each row represents an aHOR-hap, with the rightmost column indicating the superpopulation (AFR: African; AMR: American; EAS: East Asian; EUR: European; SAS: South Asian). Accompanying bar plots display the size of each aHOR-hap, color-coded by distinct HOR patterns. Schematic representations below each heatmap illustrate representative HOR structures for each haplogroup. See also Supplementary Fig. 1.

**Extended Data Figure 4:**
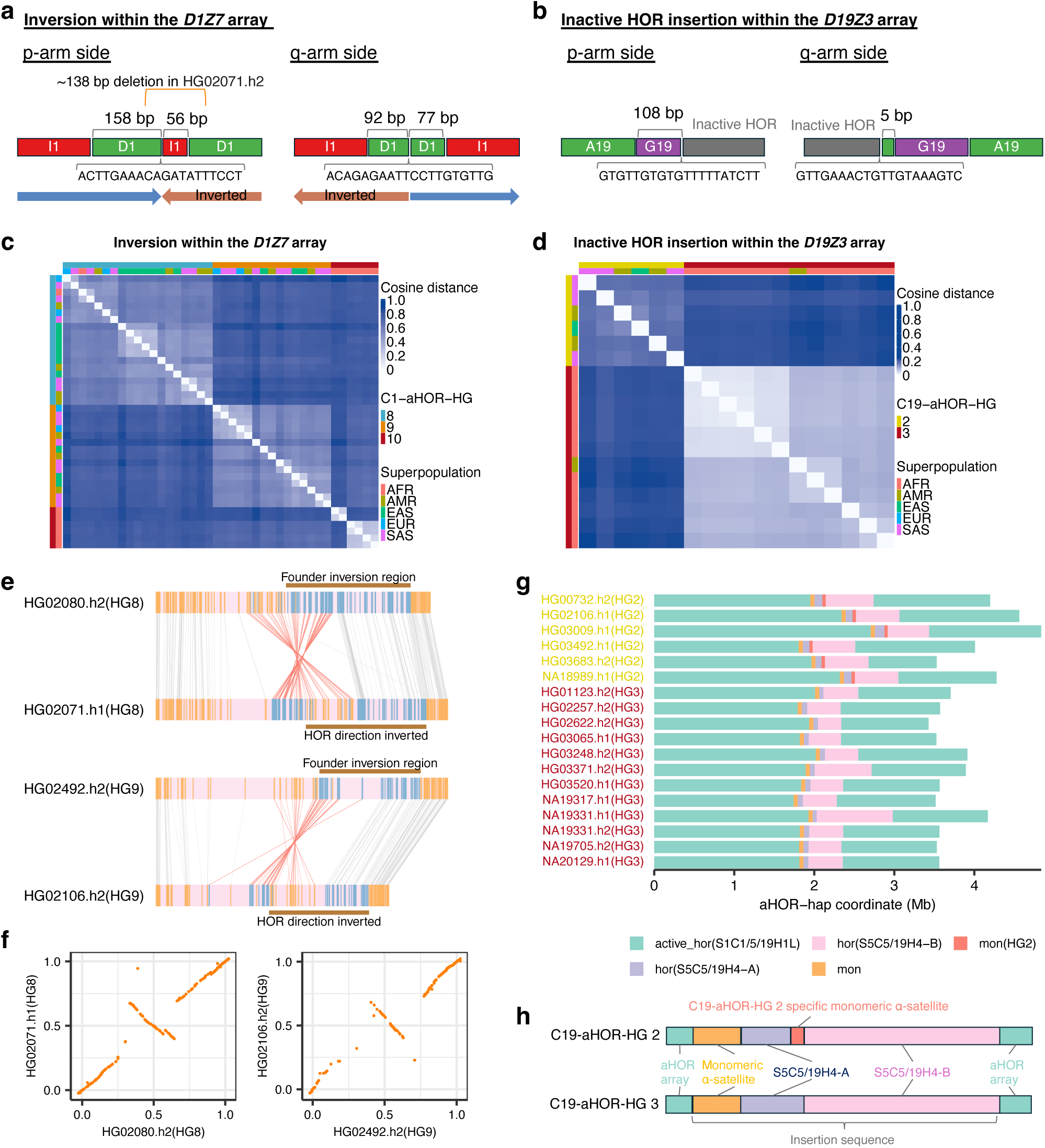
Structural characteristics of the chromosome 1 inversion and chromosome 19 inactive HOR insertion. (a) Detailed structural depiction of chromosome 1 inversion boundaries on the p-arm and q-arm sides. On the p-arm side, all aHOR-haps have a junction connecting a 158 bp forward-oriented D1 monomer to a 56 bp inverted I1 monomer, except for the HG02071.h2 aHOR-hap, which appears to have a deletion at this junction. On the q-arm side, all aHOR-haps exhibit a junction connecting a 92 bp inverted D1 monomer to a 77 bp forward-oriented D1 monomer. (b) Detailed structural depiction of chromosome 19 inactive HOR insertion boundaries on the p-arm and q-arm sides. On the p-arm side, all aHOR-haps contain a junction connecting a 108 bp G19 monomer to inactive HOR sequences. On the q-arm side, all aHOR-haps exhibit a junction connecting inactive HOR sequences to a 5 bp A19 monomer. (c, d) Heatmaps illustrating cosine distances of rare k-mer profiles among aHOR-haps carrying the (c) chromosome 1 inversion and (d) chromosome 19 inactive HOR insertion, validating the clustering results shown in Fig. 2a, b through an alternative visualization method. Color bars at the top and left of each heatmap indicate stratifications by aHOR-HGs and superpopulation. (e) Examples of aHOR-haps (HG02071.h1 and HG02106.h2) that exhibit secondary inversions occurring on top of the primary inversion. Comparisons are shown with closely related aHOR-haps without those secondary inversions (HG02080.h2 and HG02492.h2, respectively). Blue lines indicate rare k-mers specific to the inverted region in aHOR-haps with the inversion, whereas orange lines indicate other rare k-mers. Rare k-mers shared between target and proxy aHOR-haps in the same orientation (indicating no inversion) are connected by gray lines, while those shared but oriented in opposite directions (indicating inversion) are connected by red lines. (f) Relative positions of rare k-mers shared between aHOR-haps with secondary inversions (HG02071.h1 and HG02106.h2) and with closely related aHOR-haps without those secondary inversions (HG02080.h2 and HG02492.h2, respectively) are plotted. (g) Composition of individual aHOR-haps of C19-aHOR-HGs 2 and 3. Each row corresponds to an aHOR-hap (C19-aHOR-HG 2 [yellow] or 3 [red]). Horizontal bars indicate the composition of repeat structures across the aHOR array coordinate (Mb). Color codes distinguish active HOR arrays [active_hor(S1C1/5/19H1L)], inactive HOR arrays [hor(S5C5/19H4-A/B)], and monomeric α-satellite segments common [mon] and specific to C19-aHOR-HG 2 [mon(HG2)]. The inactive α-satellite HOR array S5C5/19H4−B, which comprises the majority of the inserted sequence, is also known as the *D19Z1* array. (h) Schematic comparison of the aHOR array structure between C19-aHOR-HGs 2 and 3. C19-aHOR-HG 2 aHOR-haps contain specific monomeric α-satellite segments, which are absent in C19-aHOR-HG 3.

**Extended Data Figure 5:**
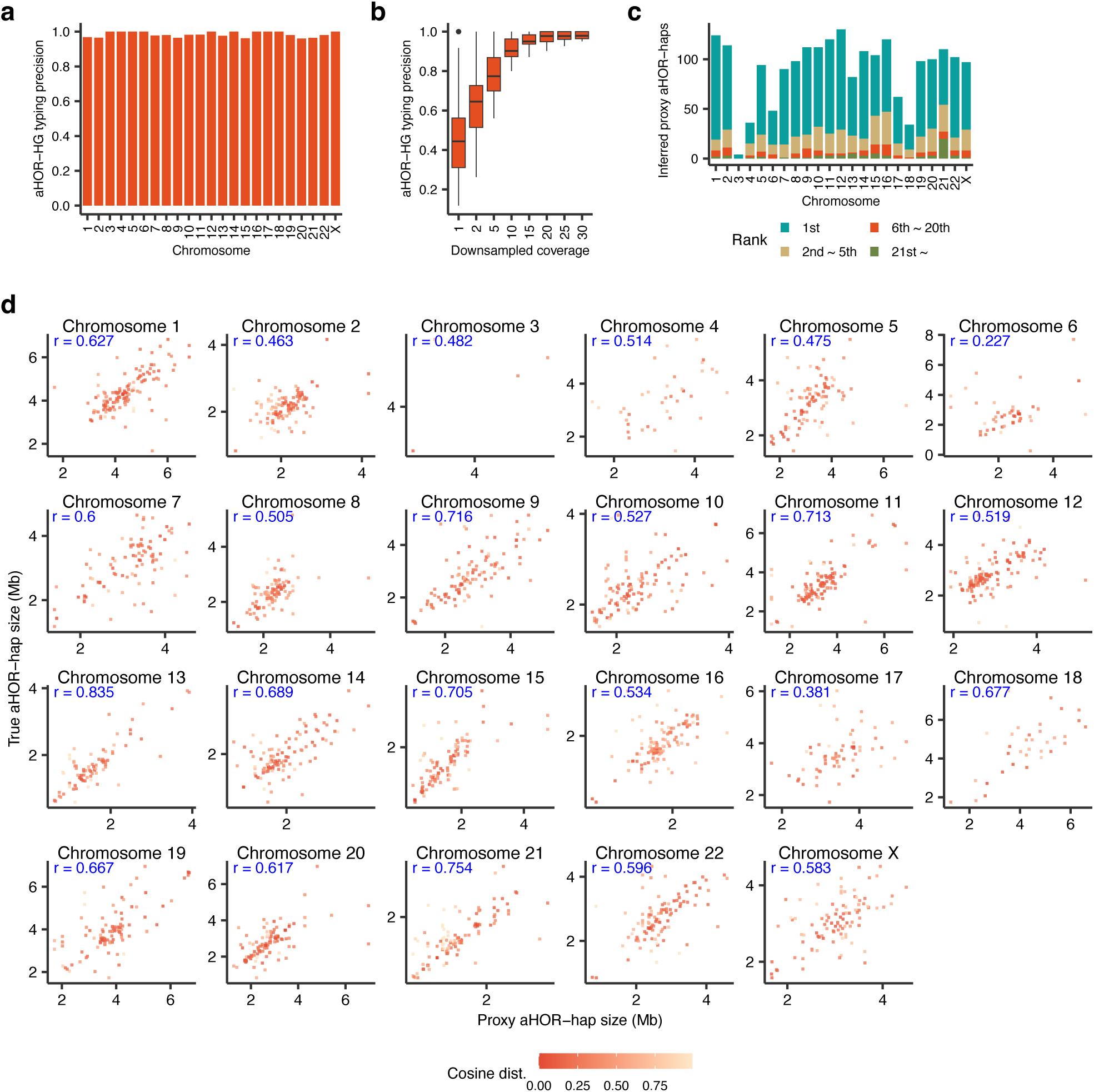
Performance evaluation of the ascairn algorithm. (a, b, c) Evaluations of ascairn performance for clustering with the constraint that each haplogroup contains at least five aHOR-haps. (a) Precision of the ascairn haplogroup typing evaluated by “leave-one-individual-out” cross-validation. See also Fig. 3b. (b) Precision analysis across varying sequence coverage levels, illustrated by boxplots depicting precision variability per chromosome after downsampling. See also Fig. 3c. (c) Counts of haplotypes selected by ascairn via “leave-one-individual-out” cross-validation for each chromosome, stratified according to cosine distance ranking relative to the true haplotype (1st, 2nd– 5th, 6th–20th, and 21st or greater). See also Fig. 3d. (d) Accuracy of aHOR-hap size estimation from proxy aHOR-haps. Each panel shows the correlation between predicted and true aHOR-hap sizes for a given chromosome, evaluated via leave-one-individual-out cross-validation. Colors represent cosine distances between the proxy and true aHOR-haps. Pearson’s *r* is indicated in each panel.

**Extended Data Figure 6:**
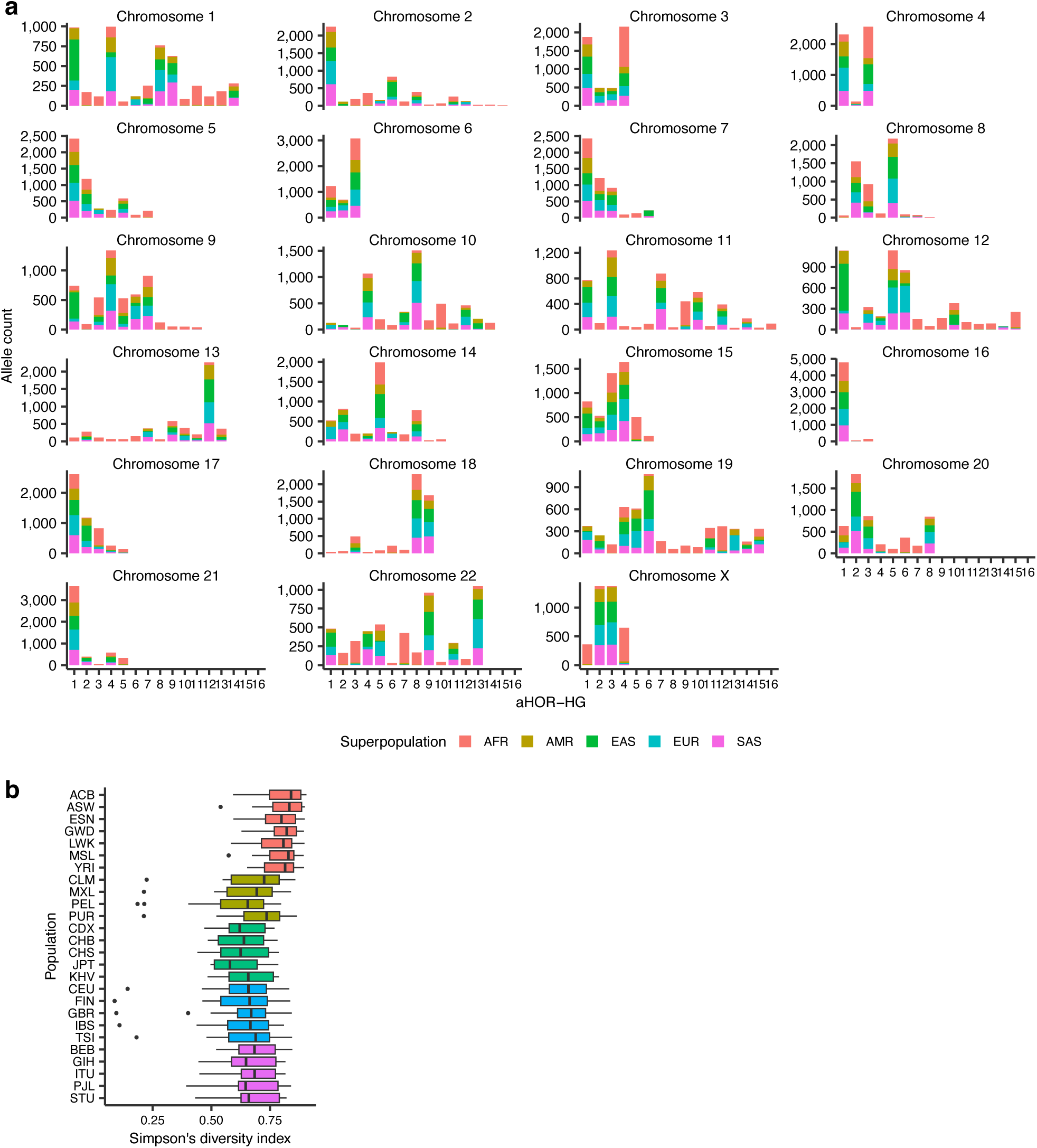
Population distribution and diversity of aHOR-HGs. (a) Allele counts of aHOR-HGs across chromosomes in the 1000 Genomes Project populations. Stacked bar plots represent the total number of alleles assigned to each aHOR-HG (x-axis) for each chromosome, stratified by superpopulation (AFR, African; AMR, American; EAS, East Asian; EUR, European; SAS, South Asian). Each bar reflects the combined allele count across individuals within each superpopulation. (b) Chromosome-wise haplogroup diversity per population, measured by Simpson’s diversity index. Box plots show the distribution of the index values across chromosomes for each population, calculated from allele frequency distributions of aHOR-HG assignments inferred by ascairn. Only chromosomes with five or more haplogroups were included in the analysis. Higher index values indicate greater haplogroup diversity within each population. See also Fig. 4b.

**Extended Data Figure 7:**
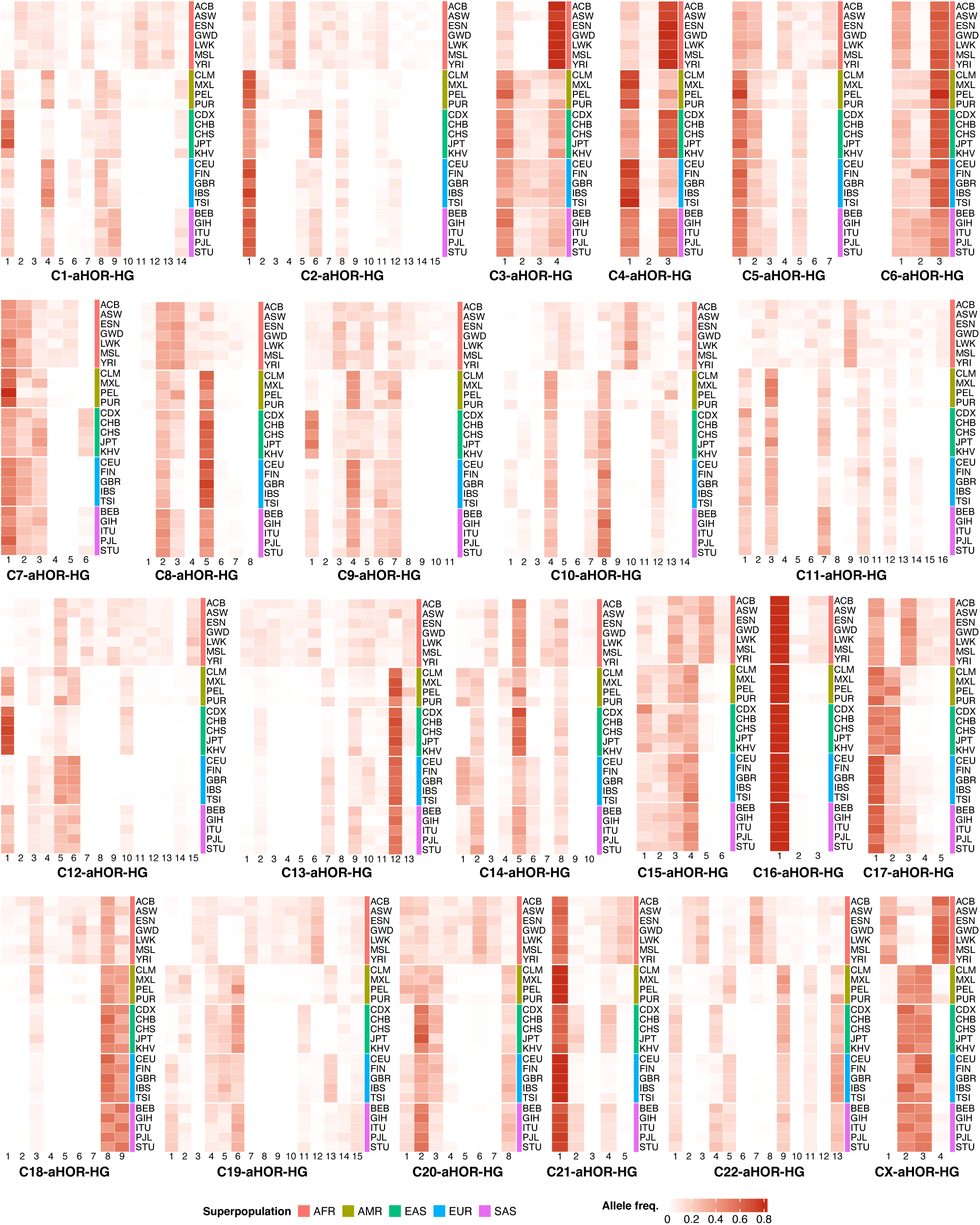
Variation of aHOR-HGs across populations. Allele frequencies of aHOR-HGs across chromosomes in populations from the 1000 Genomes Project. Heatmap cell color intensity indicates the allele frequency for each haplogroup within each population. Superpopulation annotations are displayed as colored bars on the right side of the heatmap (AFR, African; AMR, American; EAS, East Asian; EUR, European; SAS, South Asian). See also Fig. 4a.

**Extended Data Figure 8:**
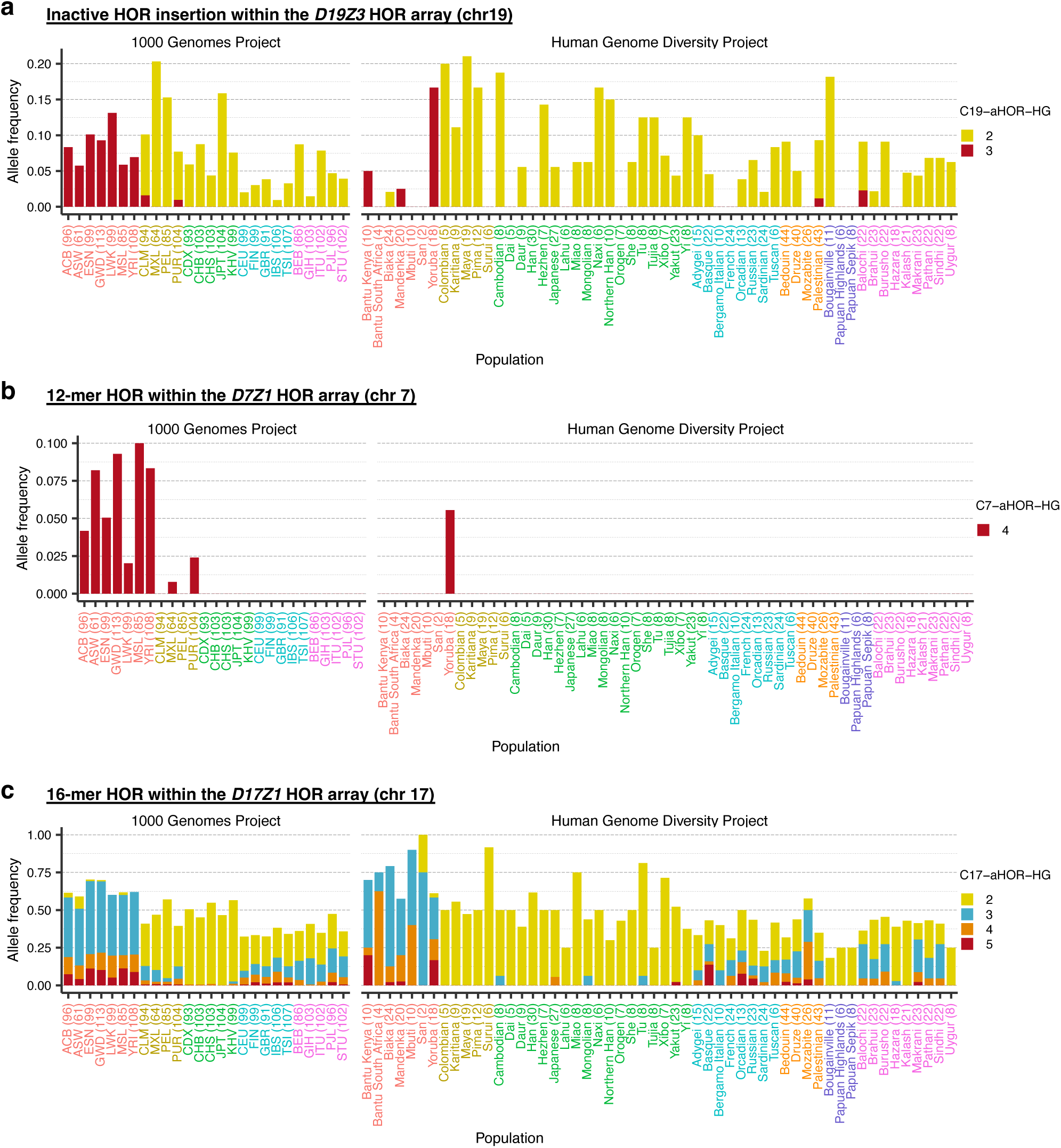
Allele frequencies of aHOR-HGs and associated structural variations across populations. (a, b, c) Allele frequencies of aHOR-HGs specific to (a) inactive α-satellite HOR sequence insertion within the *D19Z3* array (chromosome 19), (b) 12-mer HOR within the *D7Z1* array (chromosome 7) and (c) 16-mer HOR within the *D17Z1* array (chromosome 17) across global populations from the 1000 Genomes Project and the Human Genome Diversity Project. Frequencies were calculated by summing the corresponding haplogroup allele frequencies, with bar colors indicating superpopulations.

**Extended Data Figure 9:**
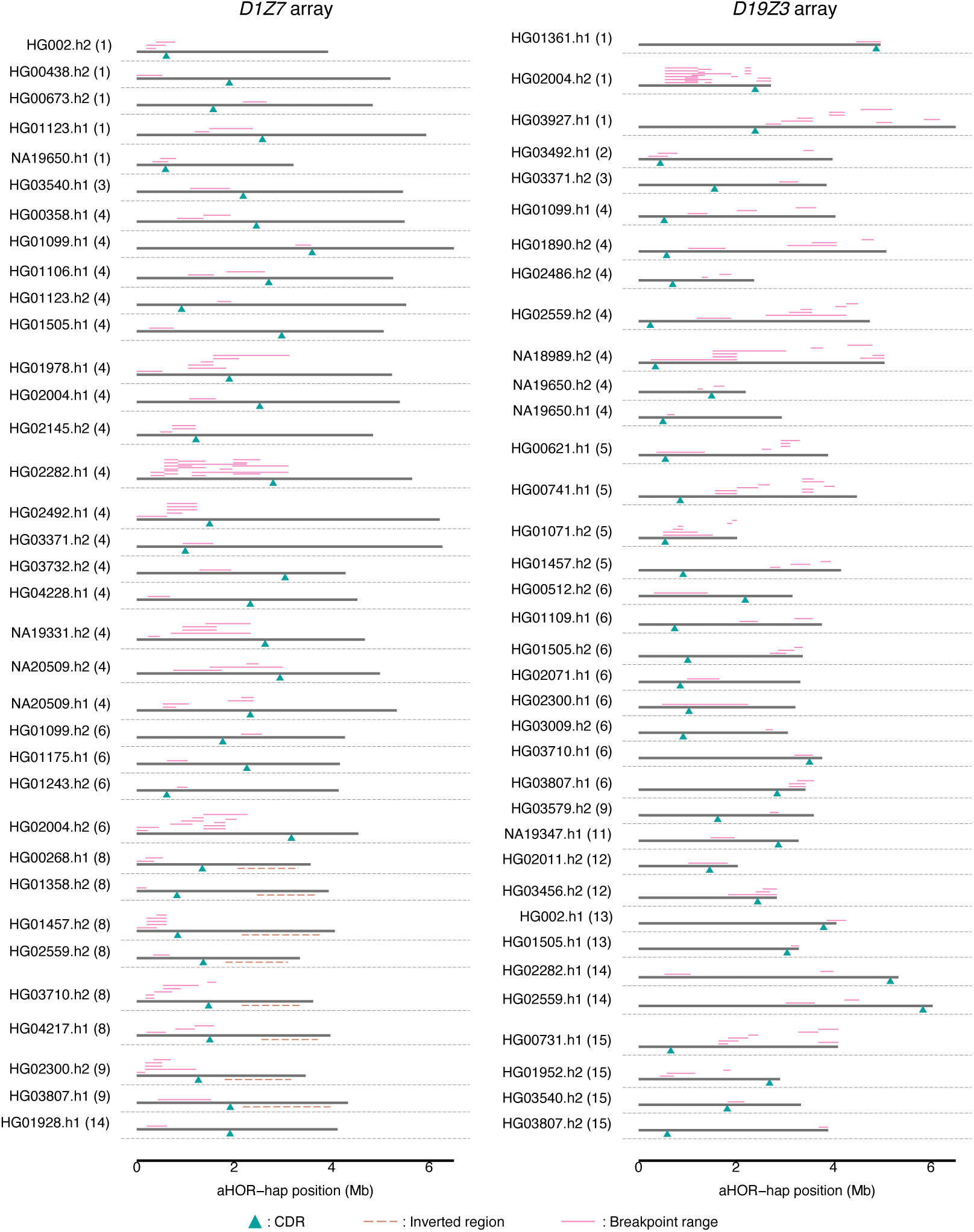
Breakpoint mapping in 142 oligodendroglioma samples. Breakpoints are grouped by proxy aHOR-haps, with the corresponding aHOR-HGs shown in parentheses. Pink horizontal segments indicate putative breakpoint regions mapped onto the corresponding proxy aHOR-hap sequence. Cyan triangles denote centromere dip region (CDR) positions, and brown dashed lines indicate known inversion boundaries.

**Extended Data Figure 10:**
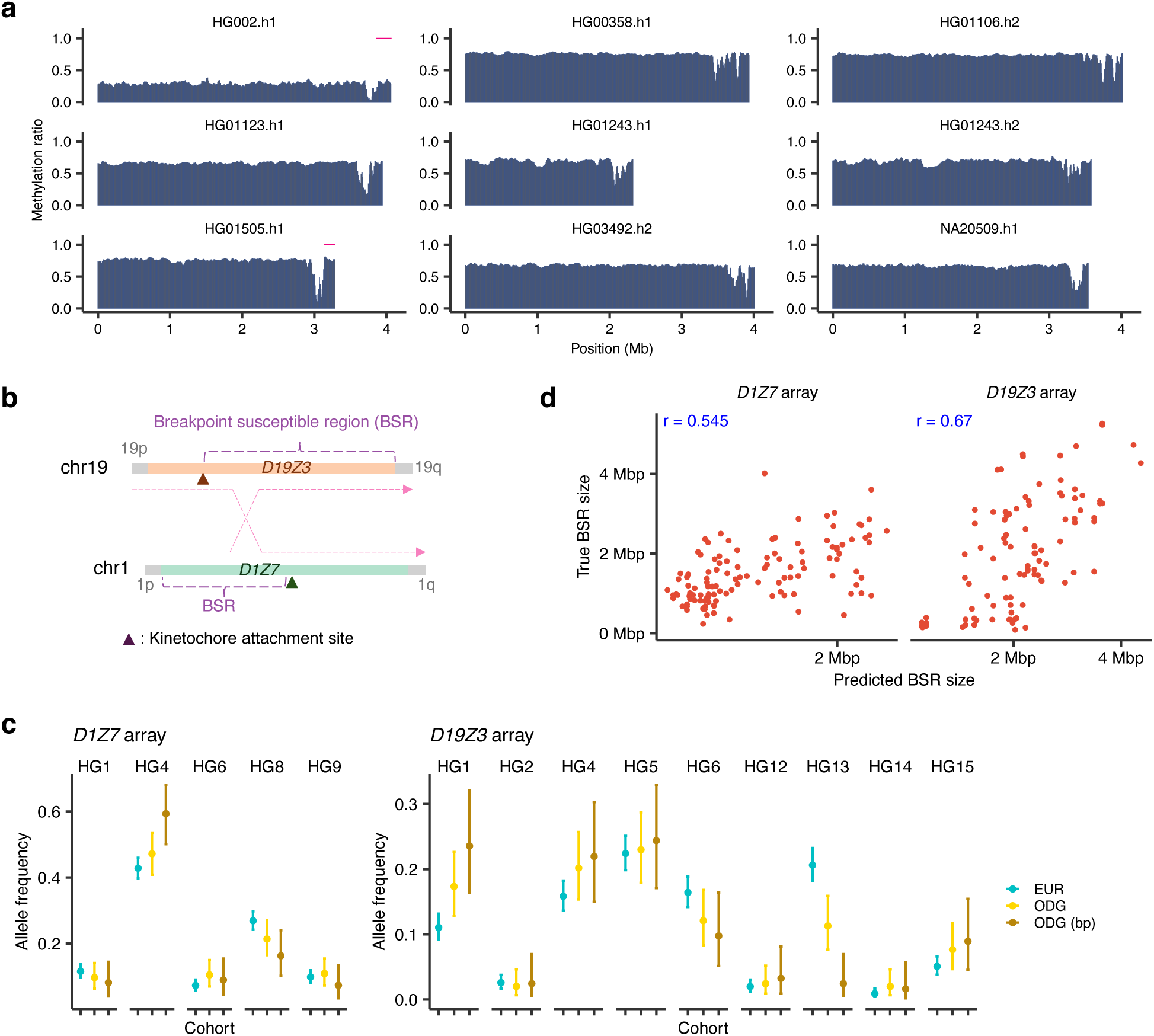
Allele frequencies and allele-selection strength of aHOR-HGs. (a) Methylation plot of aHOR-haps in C19-aHOR-HG 13. Each aHOR-hap contains a centromere dip region (CDR), characterized by a sharp decrease in methylation ratio, representing the kinetochore attachment site. In each aHOR-hap, ranges of breakpoints associated with the aHOR-hap are represented by horizontal pink lines. (b) Schematic of the breakpoint-susceptible region (BSR) for oligodendrogliomas on chromosomes 1 and 19. (c) Allele frequency distributions of aHOR-HGs, focusing on *D1Z7* and *D19Z3* HOR arrays. Comparisons were performed among control populations (European [EUR; teal]), tumor cohorts without allele restrictions (ODG [yellow]), and tumor cohorts restricted to alleles with identified rearrangement breakpoints (ODG (bp) [brown]). Error bars indicate standard deviations. (d) Scatter plots illustrating correlations between the breakpoint-susceptible region (BSR) sizes of each reference aHOR-hap (true) and those of their closest predicted counterparts identified by minimal cosine distance (predicted). Results are shown separately for *D1Z7* (left) and *D19Z3* (right) HOR arrays. Each point represents an individual aHOR-hap.

## Notes

### Competing Interest Statement

The authors have declared no competing interest.

### Summary of Updates

To correct the formatting issues in Figure 3, Figure 5, and the legend of Figure 5"

