## Supplementary Figure for "Rare k-mers reveal centromere haplogroups underlying human diversity and cancer translocations"

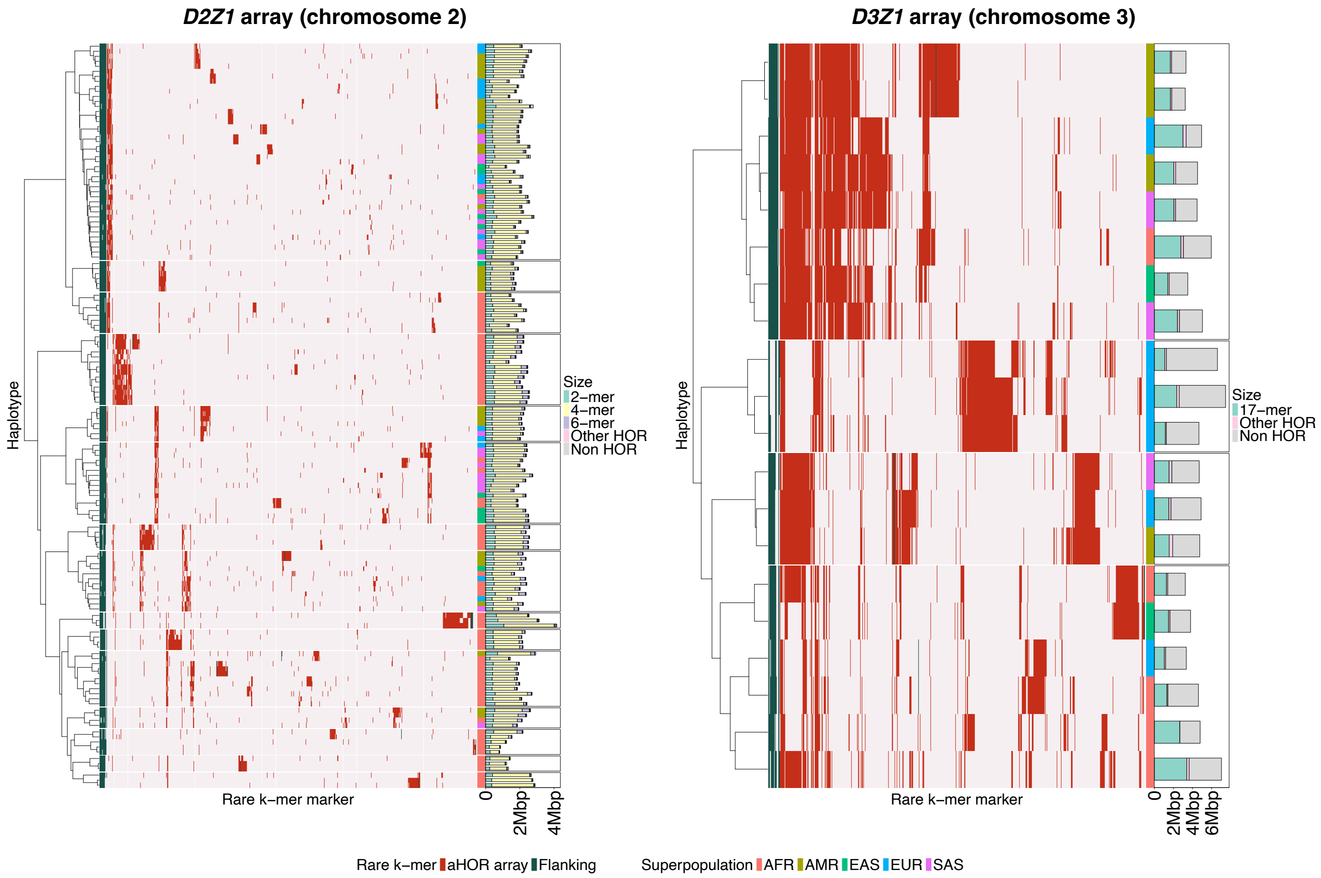


**Supplementary Figure 1: Heatmaps showing rare k-mer presence/absence profiles across aHOR-haps of additional chromosomes.** aHOR-haps were grouped into haplogroups using hierarchical clustering. Each row represents an aHOR-hap, with the rightmost column indicating the superpopulation (AFR: African; AMR: American; EAS: East Asian; EUR: European; SAS: South Asian). Accompanying bar plots display the size of each aHOR-hap, color-coded by distinct HOR patterns.

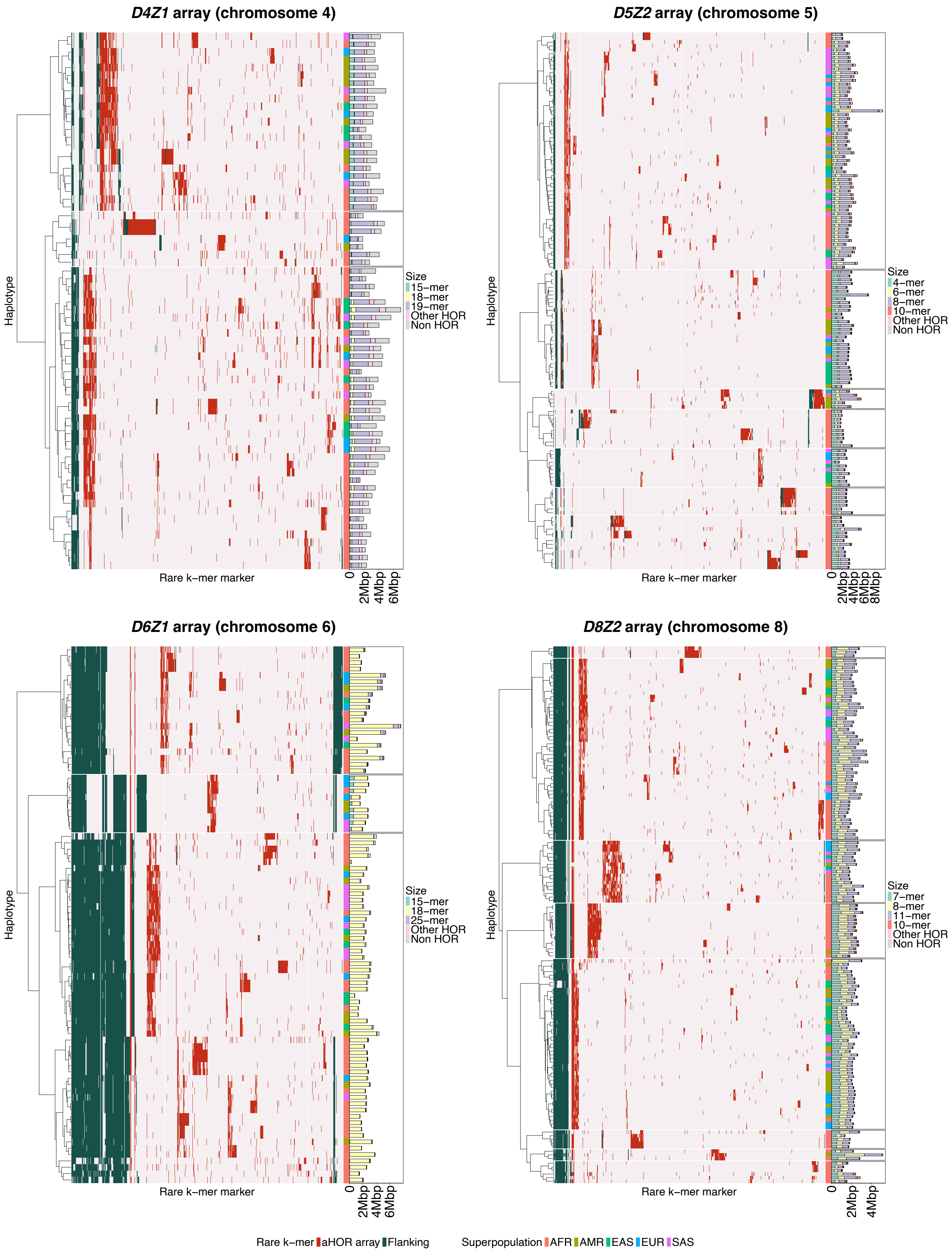


**Supplementary Figure 1 (continued)**

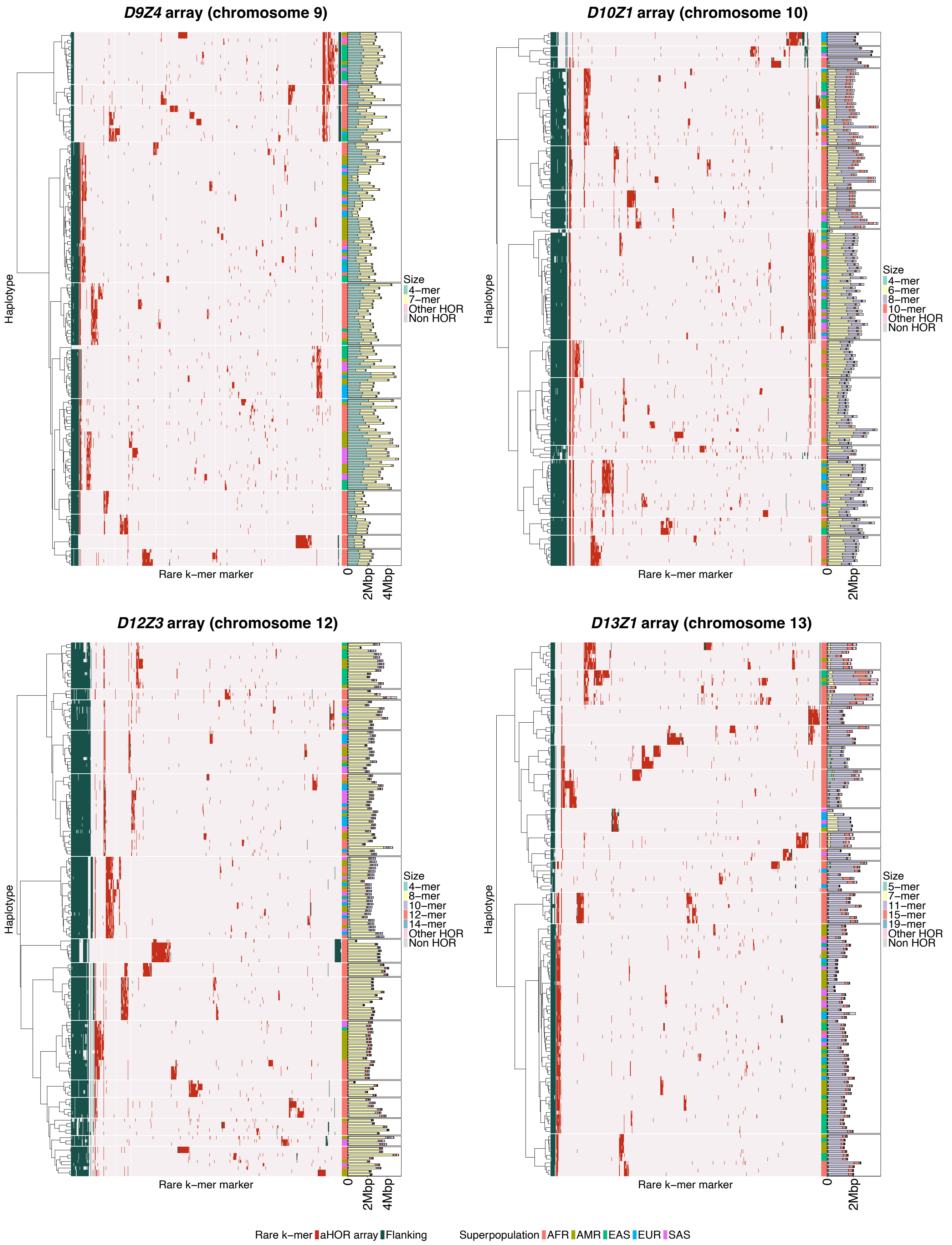


### Supplementary Figure 1 (continued)

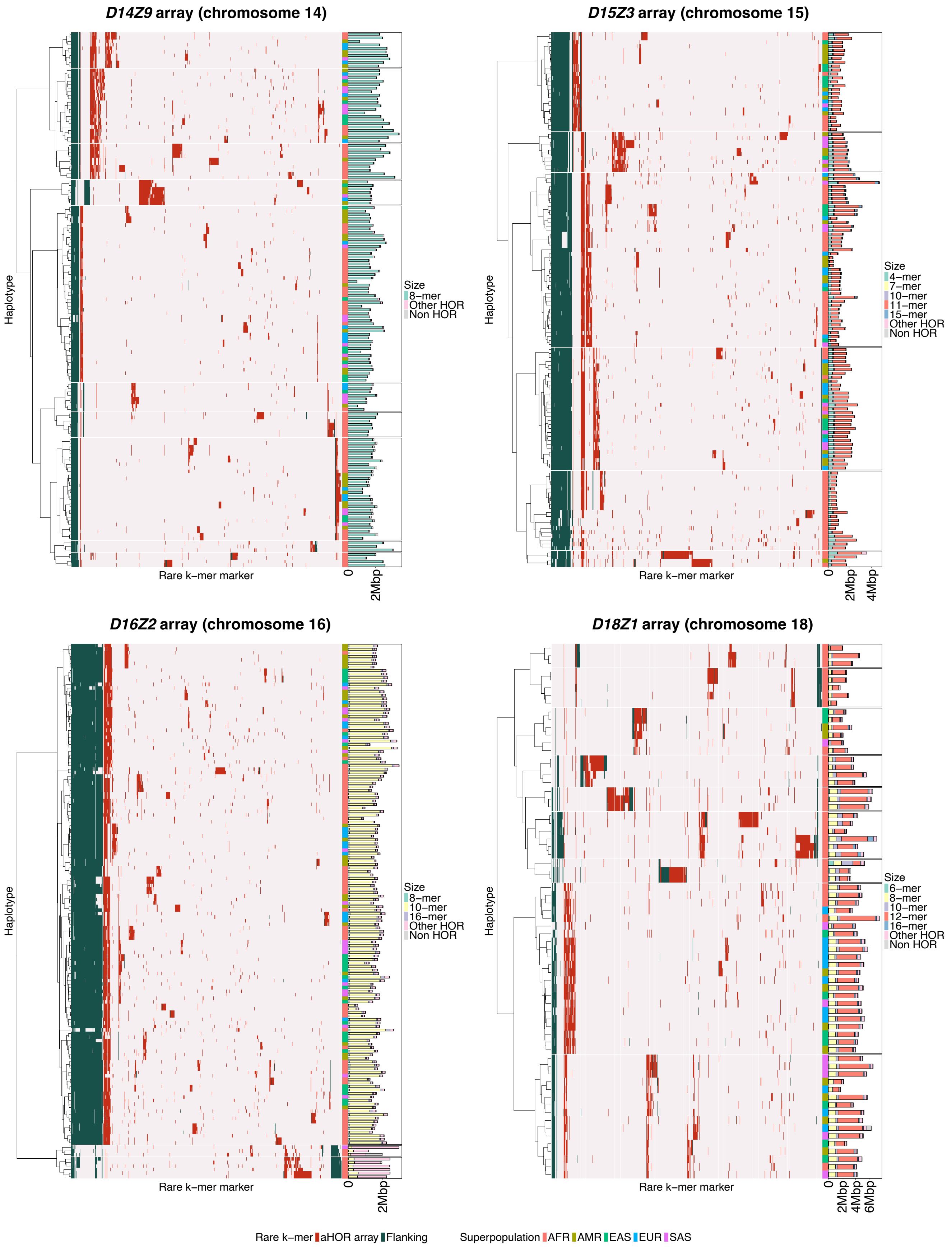


**Supplementary Figure 1 (continued)**

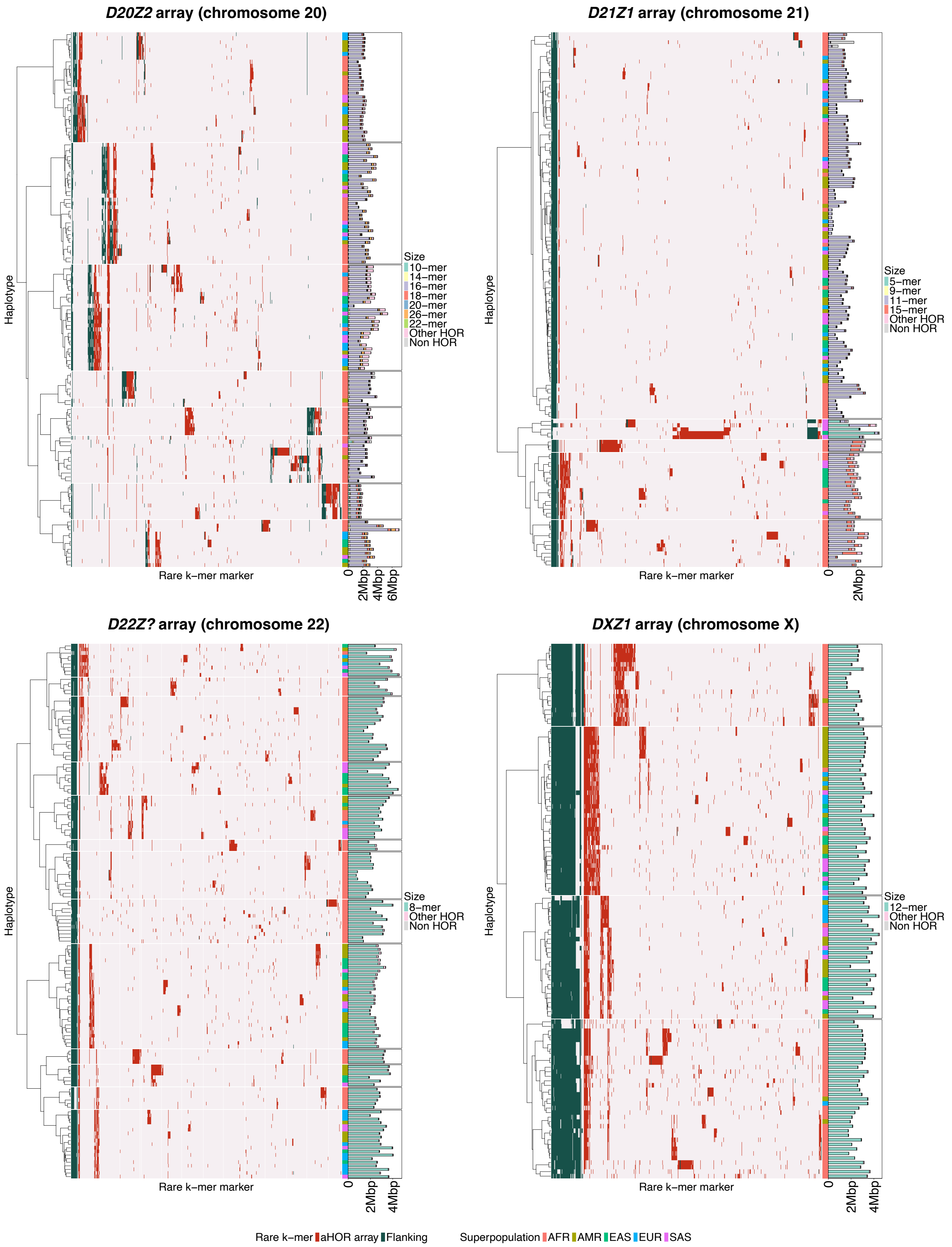


**Supplementary Figure 1 (continued)**

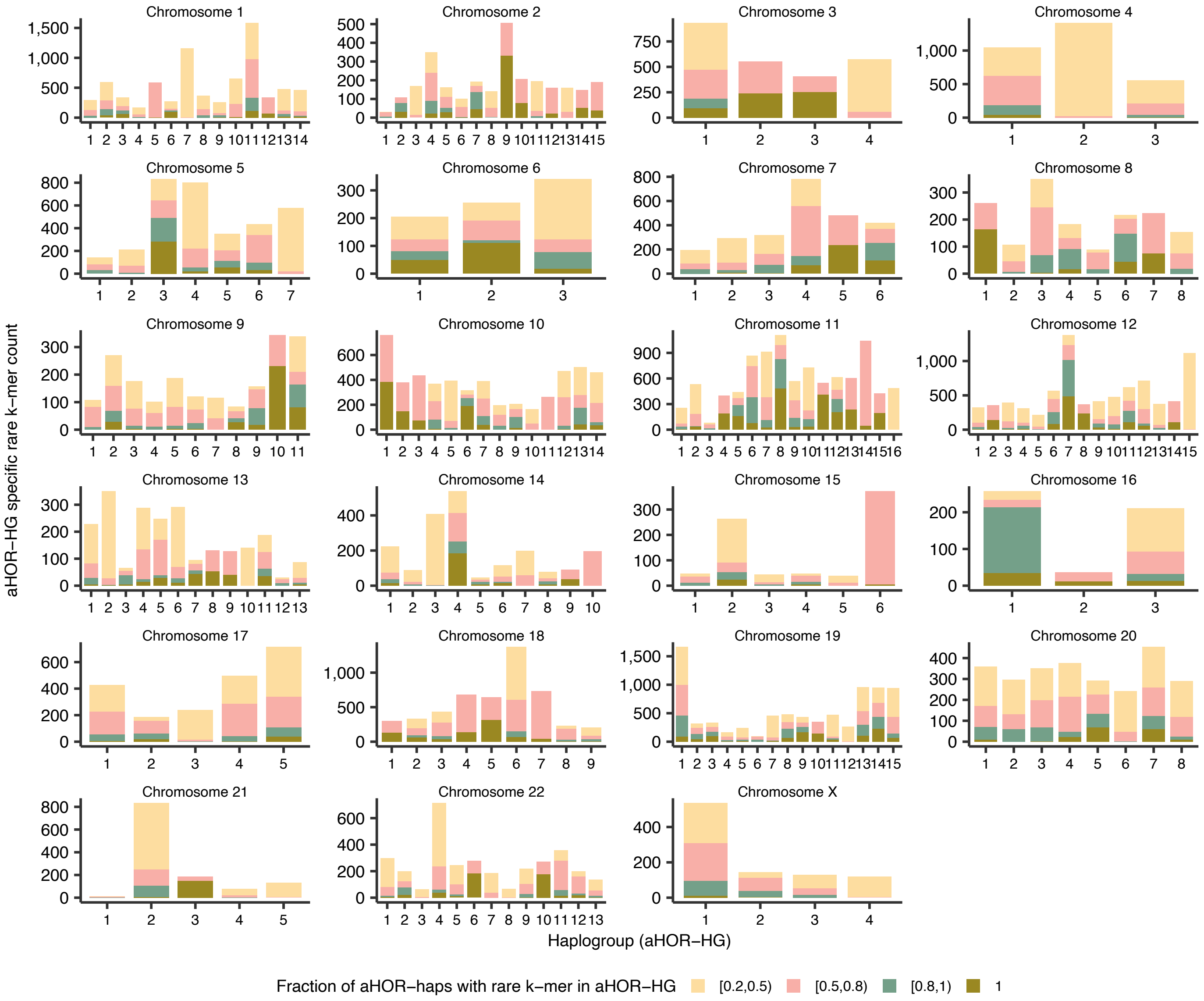


**Supplementary Figure 2: aHOR-HG-specific rare k-mer counts per haplogroup.** Bar plots show the number of rare k-mers specifically observed in each aHOR-HG on each chromosome. Only rare k-mers detected in at least two aHOR-haps were included, and further restricted to those present in ≥0.2 of aHOR-haps within each aHOR-HG. Bars are stratified according to the fraction of aHOR-haps within each aHOR-HG that carry each rare k-mer: [0.2, 0.5) (yellow), [0.5, 0.8) (pink), [0.8, 1) (green), and 1 (brown).

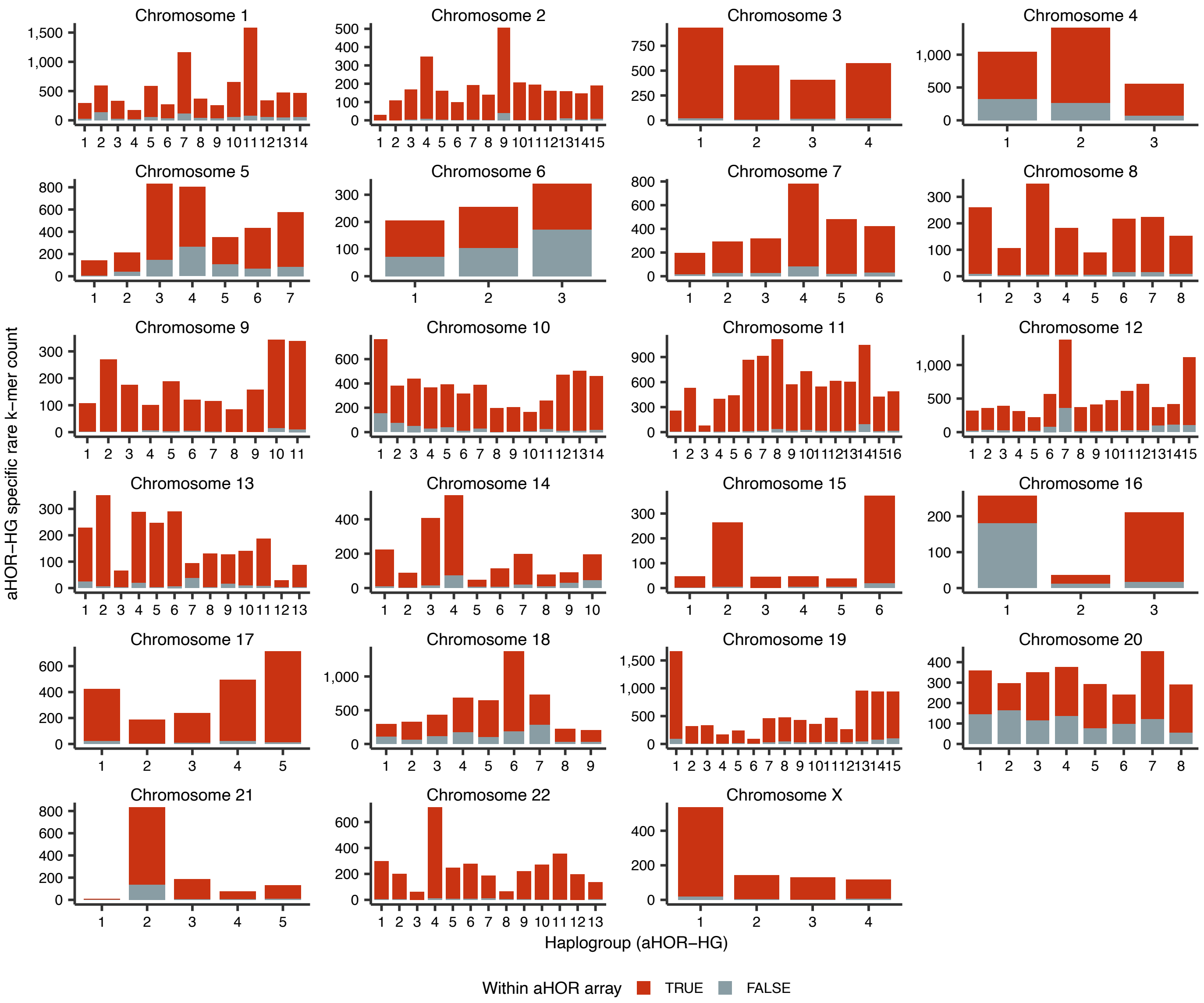


**Supplementary Figure 3: aHOR-HG-specific rare k-mer counts per haplogroup.** Bar plots show the number of rare k-mers specifically observed in each aHOR-HG on each chromosome. Only rare k-mers detected in at least two aHOR-haps were included, and further restricted to those present in ≥0.2 of aHOR-haps within each aHOR-HG. Bars are stratified according to the location of rare k-mers: Within aHOR array (red) or not (grey).

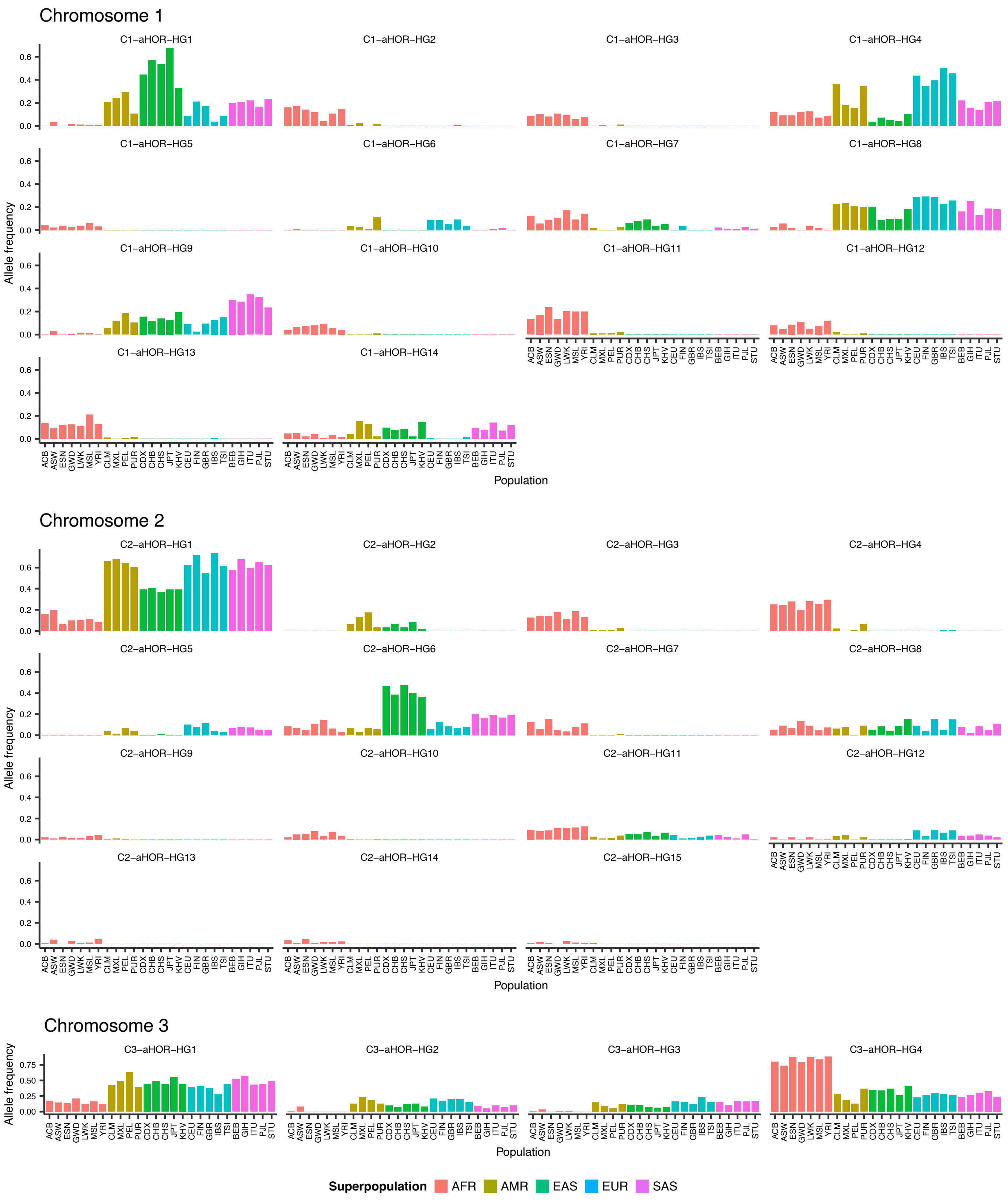


**Supplementary Figure 4: Allele frequency distribution of aHOR-HGs across chromosomes in the 1000 Genomes Project populations.** Each panel represents one chromosome, with individual subplots illustrating distinct aHOR-HGs. Populations are grouped by superpopulation (AFR: African, AMR: American, EAS: East Asian, EUR: European, SAS: South Asian), indicated by distinct colors.

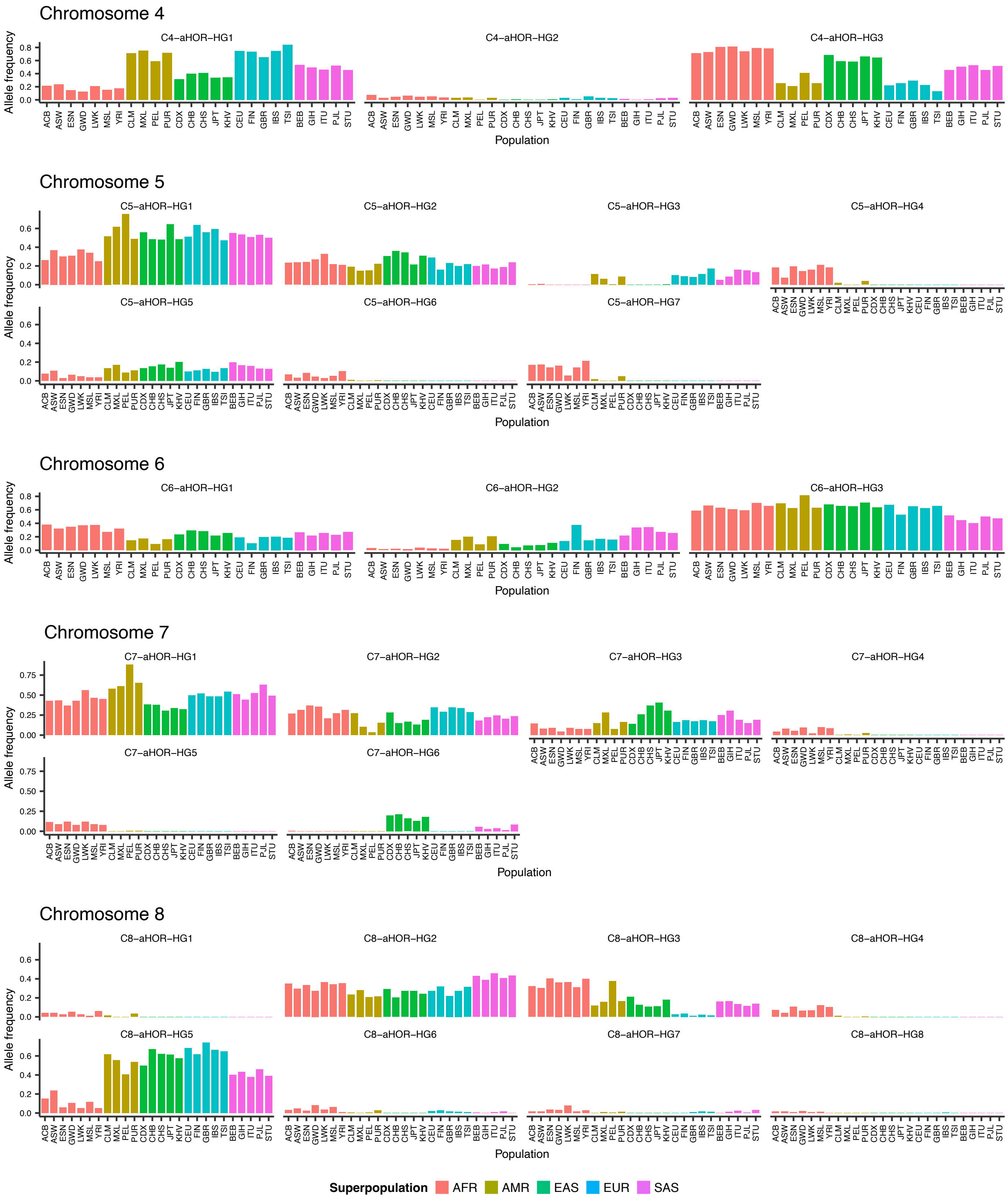


**Supplementary Figure. 4 (continued)**

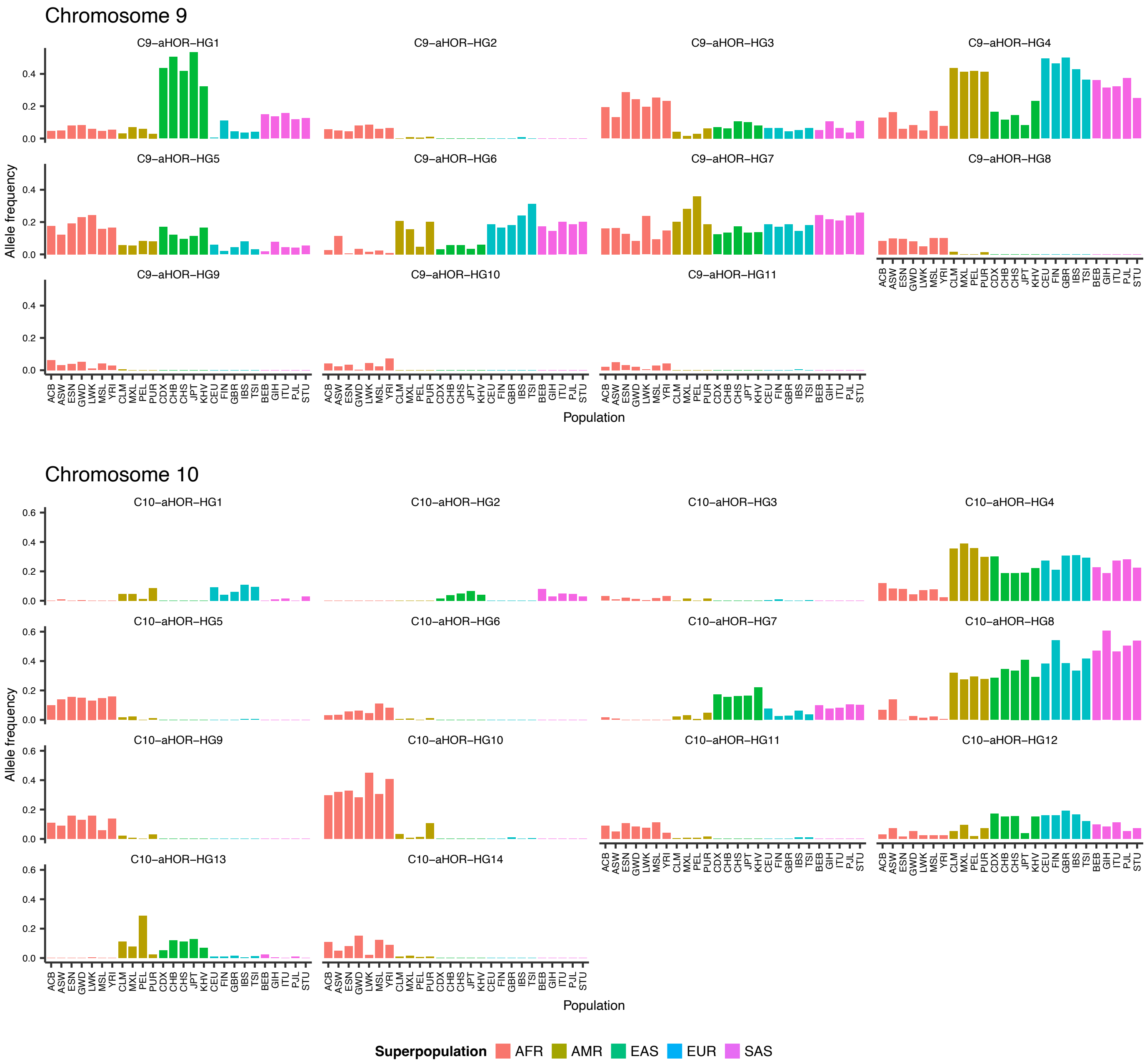


**Supplementary Figure. 4 (continued)**

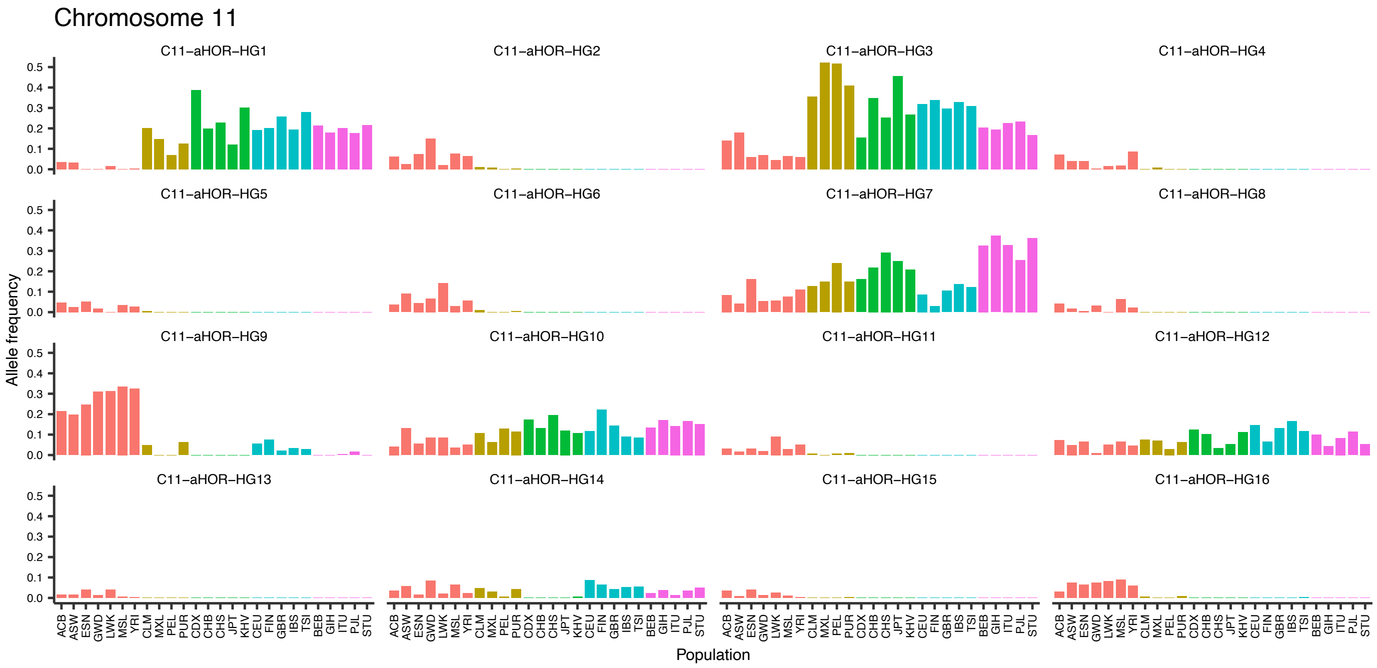


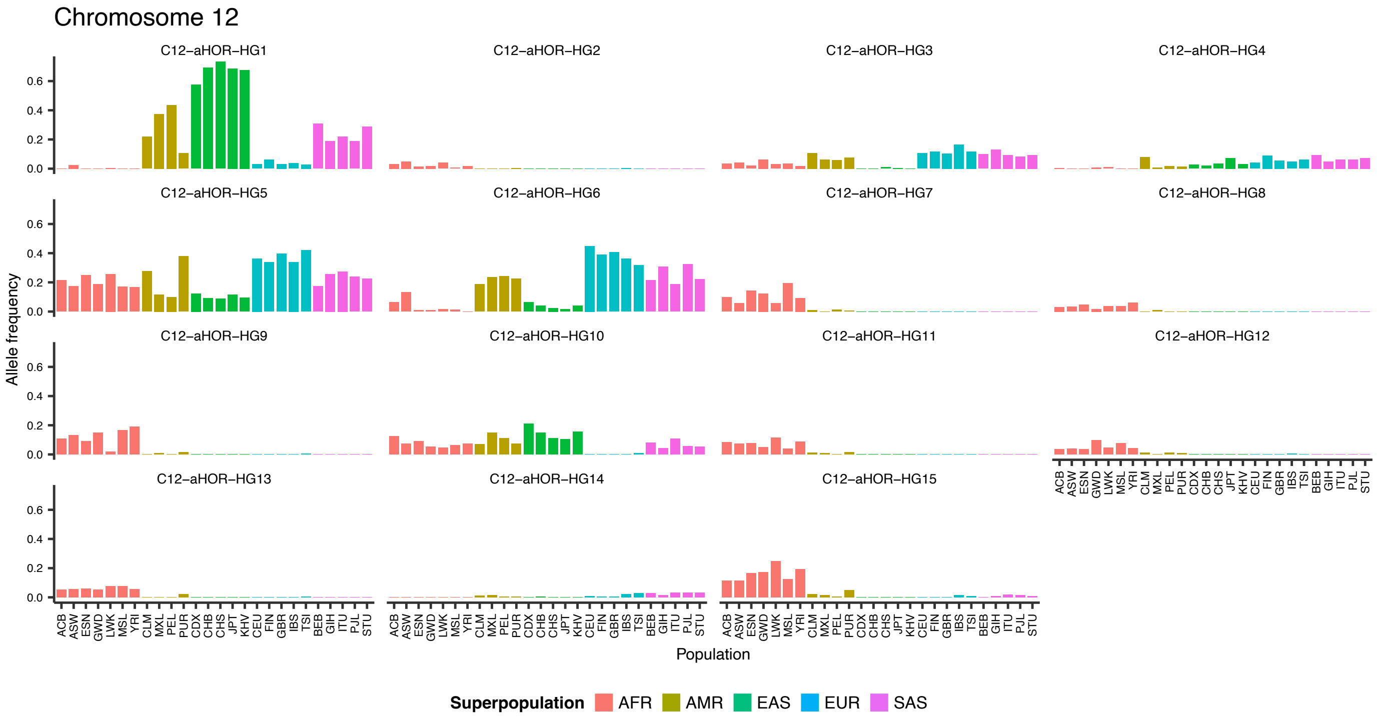


**Supplementary Figure. 4 (continued)**

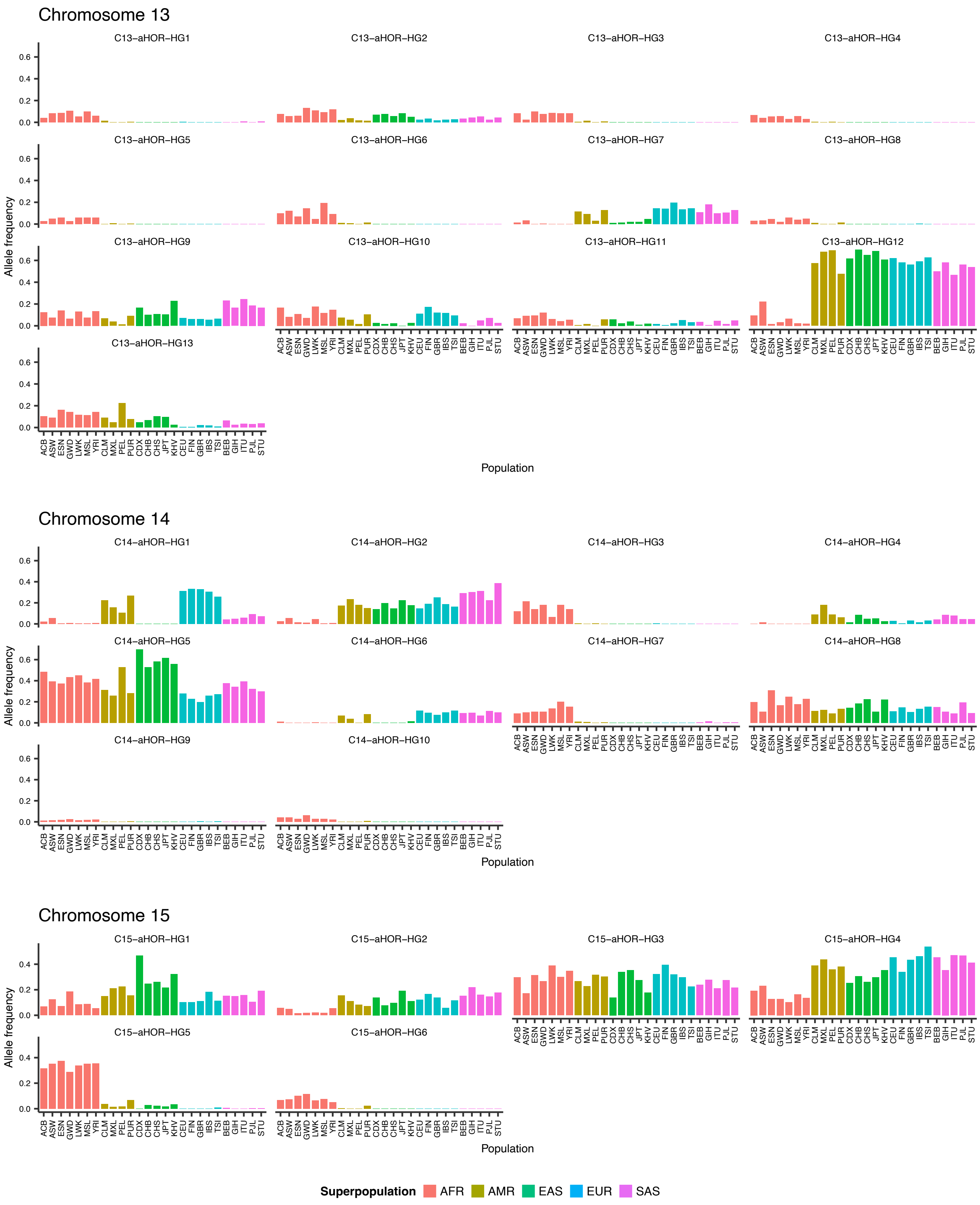


**Supplementary Figure. 4 (continued)**

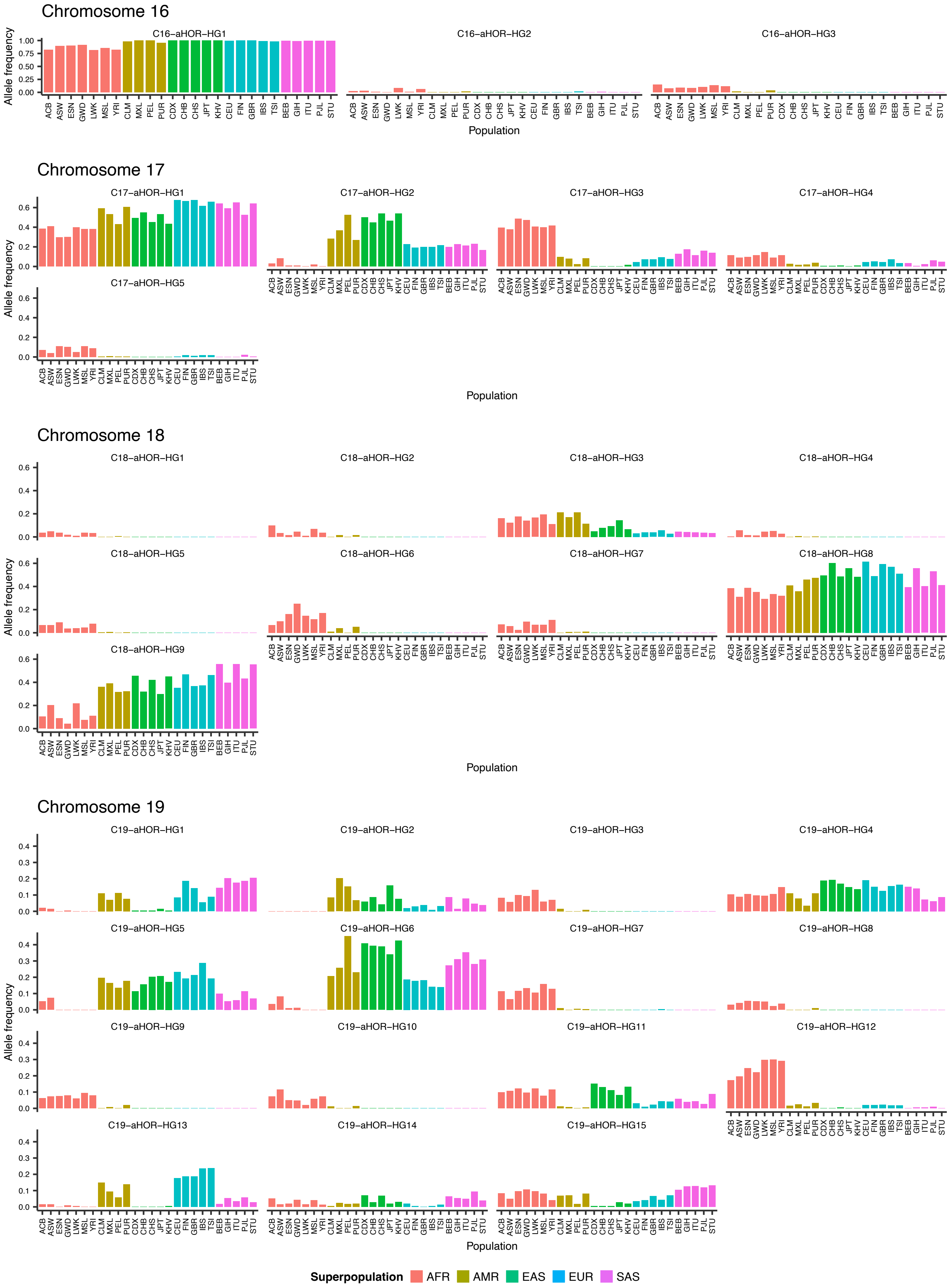


**Supplementary Figure. 4 (continued)**

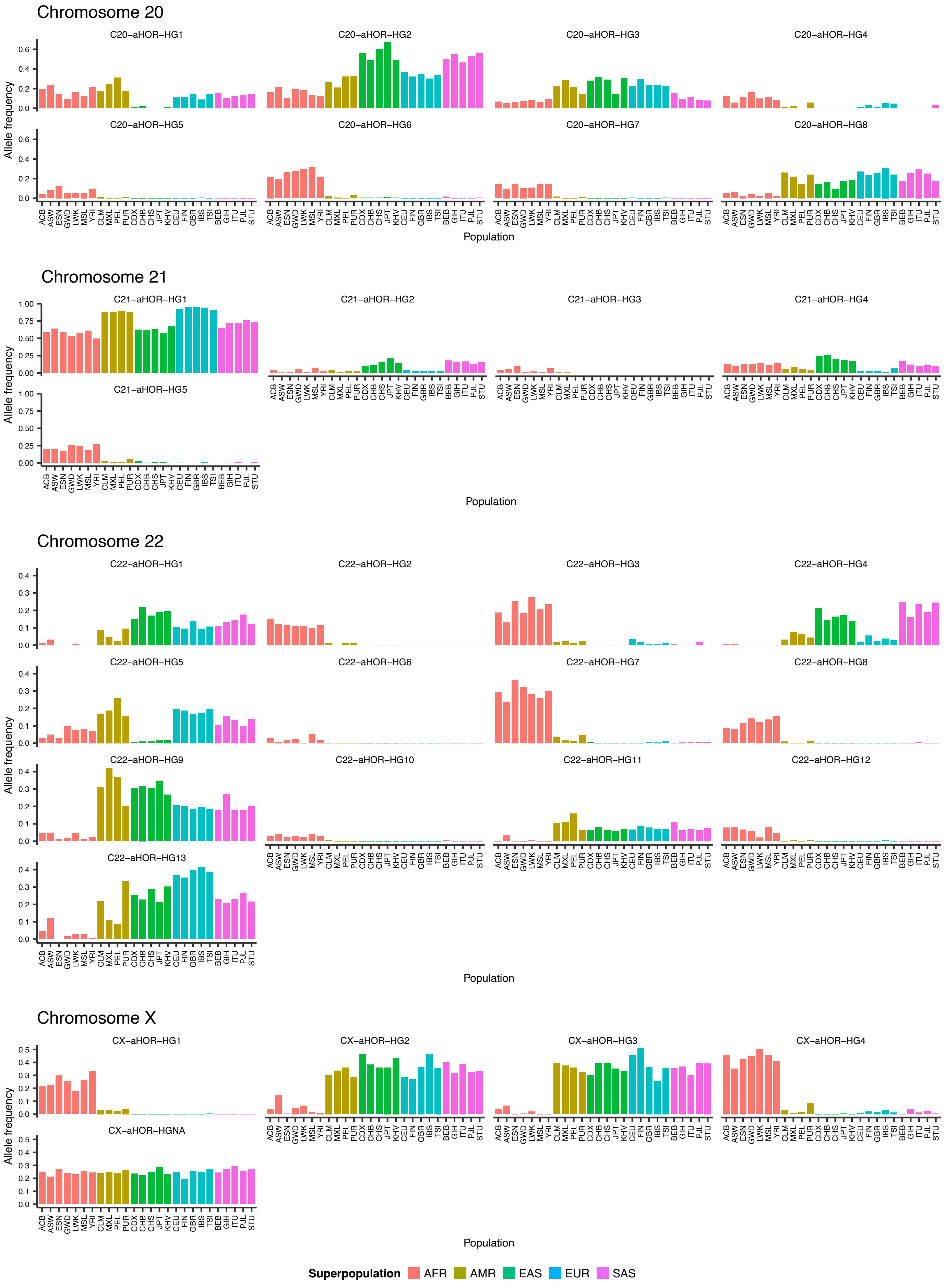


**Supplementary Figure. 4 (continued)**

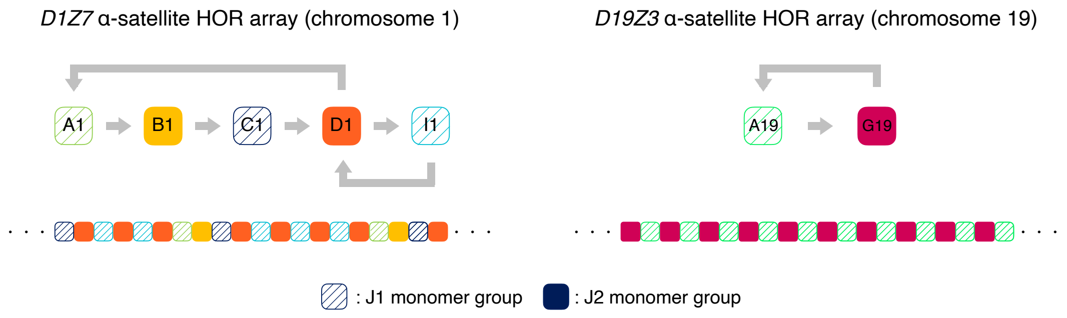


**Supplementary Figure. 5: Structures of aHOR arrays on chromosomes 1 and 19 (known as *D1Z7* and *D19Z3* arrays, respectively).** Each array is composed of tandemly arranged ~171 bp monomers, which are organized into chromosome-specific HOR structures spanning several megabases. *D1Z7* and *D19Z3* arrays belong to suprachromosomal family 1 (SF1), characterized by a repeating pattern of monomers from the J1 and J2 groups in an alternating configuration.

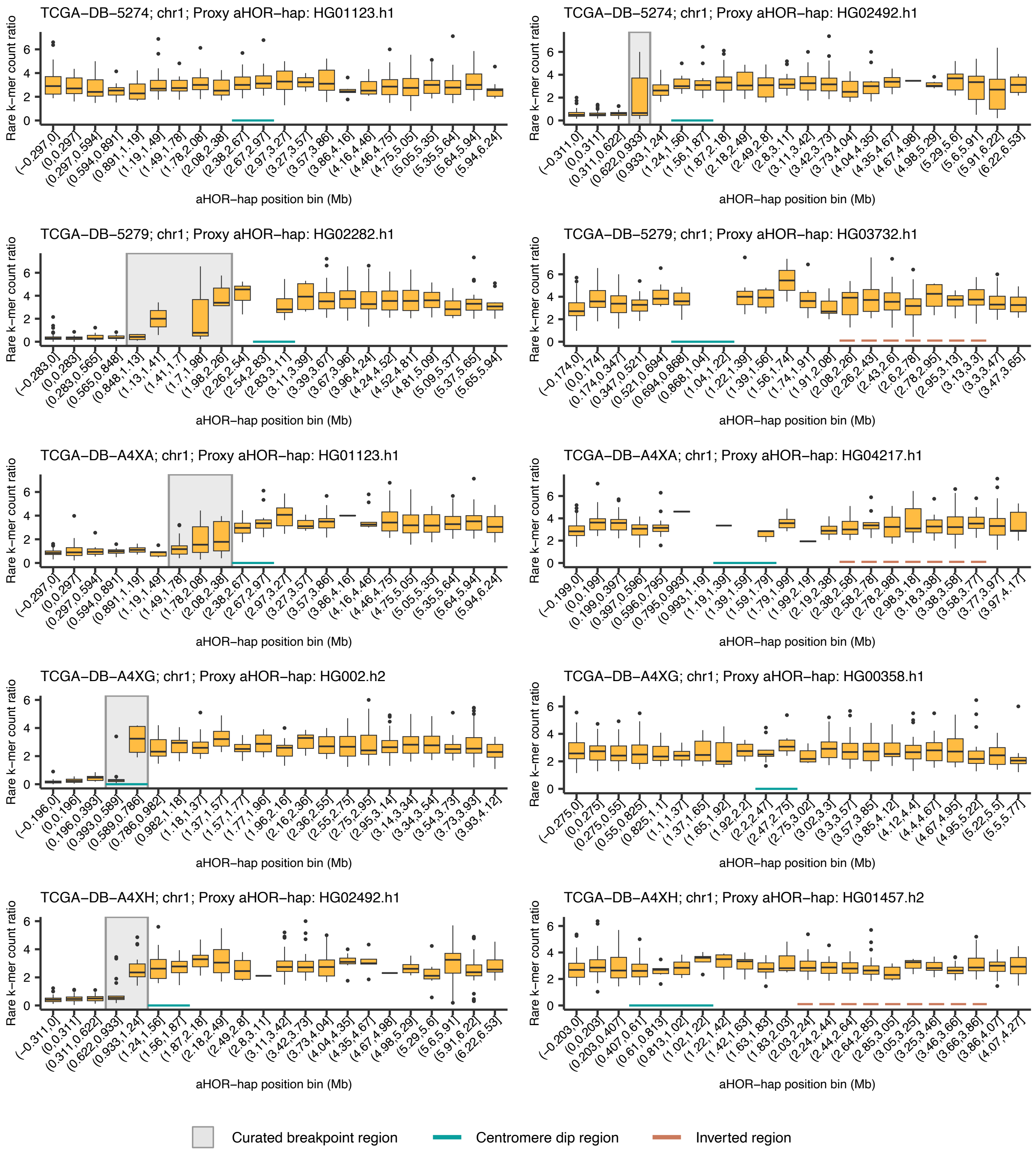


**Supplementary Figure. 6: Copy number profiles at the *D1Z7* array regions computed from 103 TCGA low-grade glioma tumor and matched normal WGS data.** Boxplots show the rare k-mer count ratio between tumor and normal samples for each positional bin along the proxy aHOR-haps (in megabases [Mb]). Shaded gray areas highlight inferred breakpoint locations of rearrangements identified by copy number alterations. Cyan horizontal bars beneath the boxplots denote centromere dip regions (CDRs), indicating kinetochore attachment sites. Brown lines indicate inverted regions.

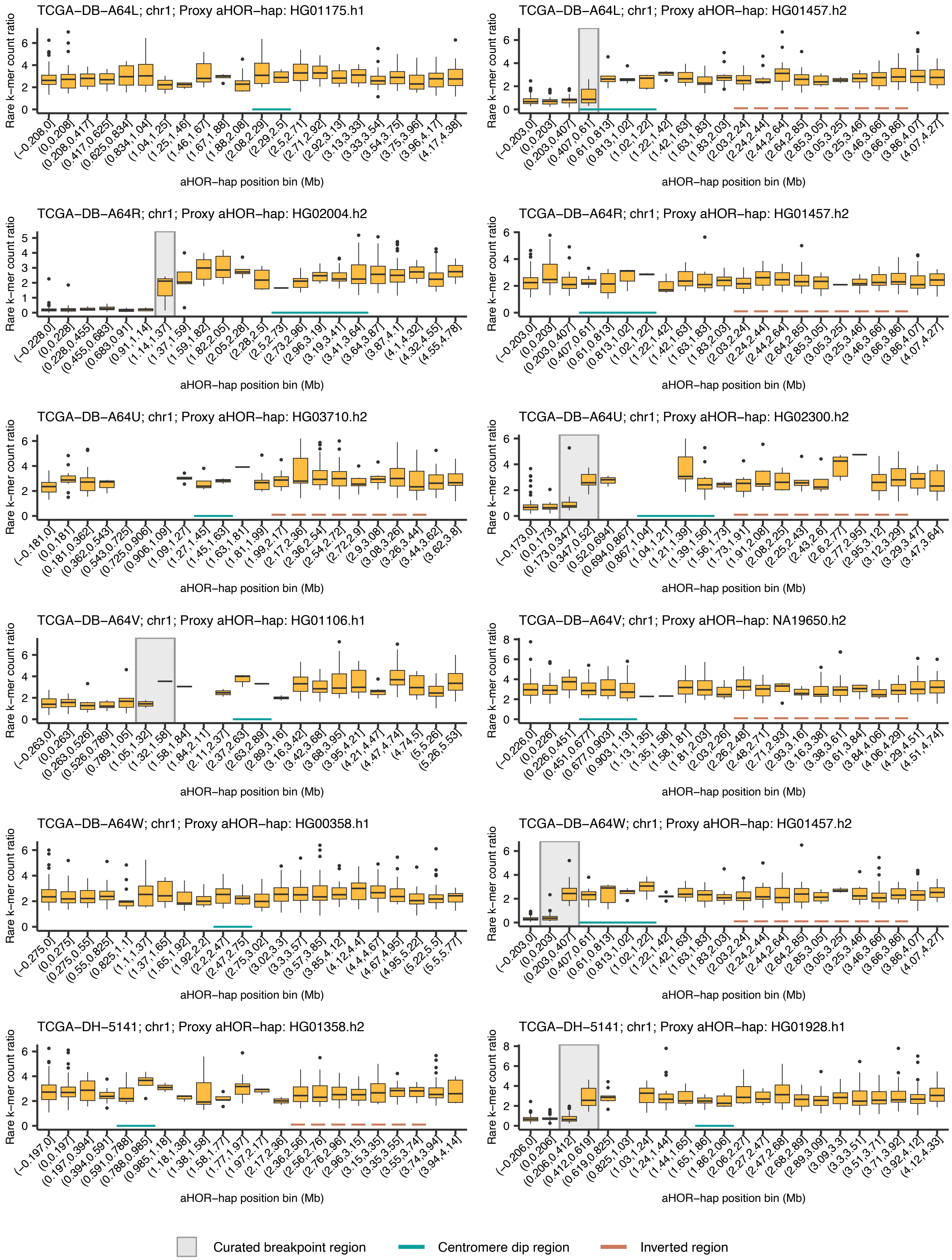


**Supplementary Figure. 6 (continued)**

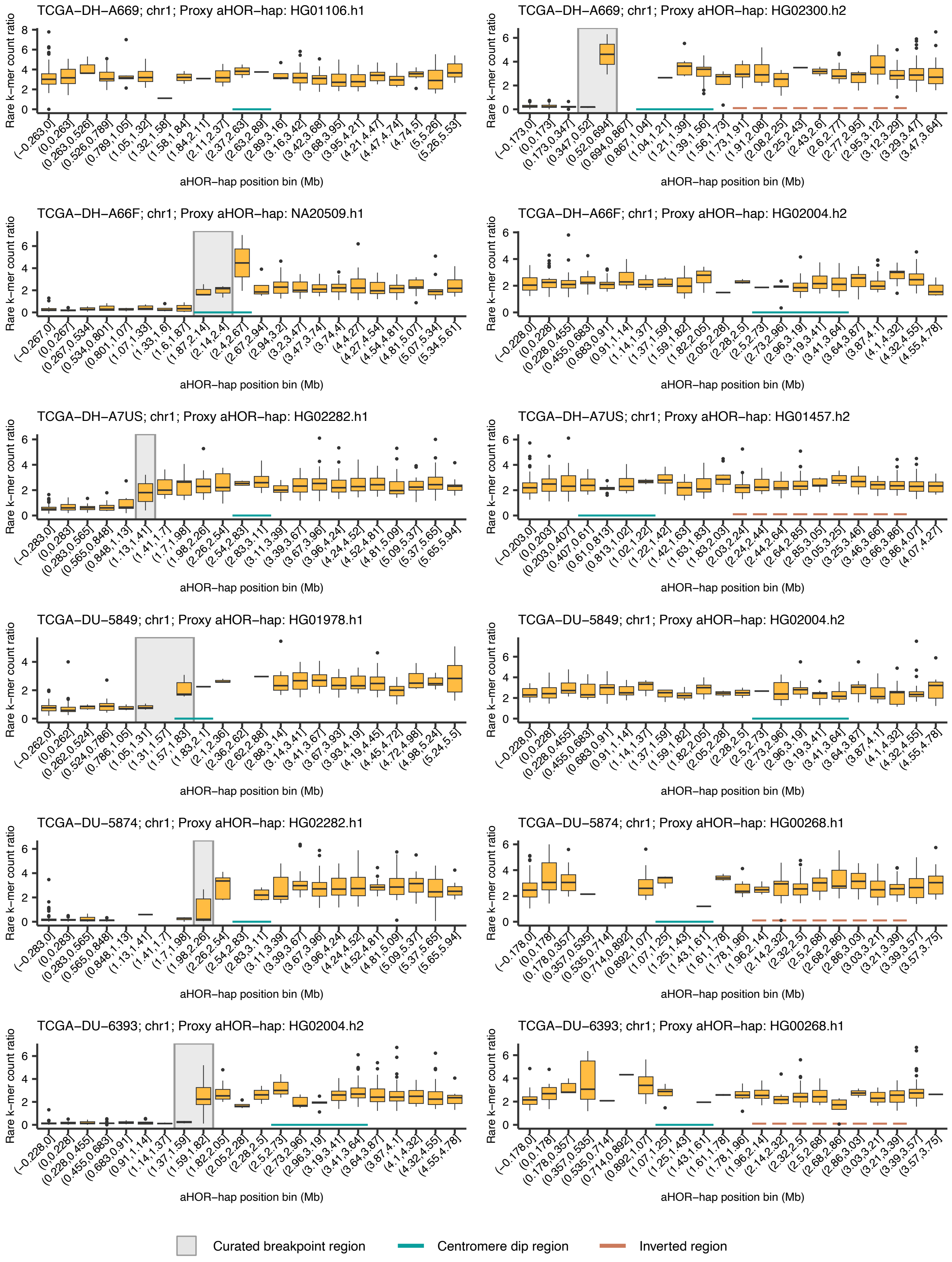


**Supplementary Figure. 6 (continued)**

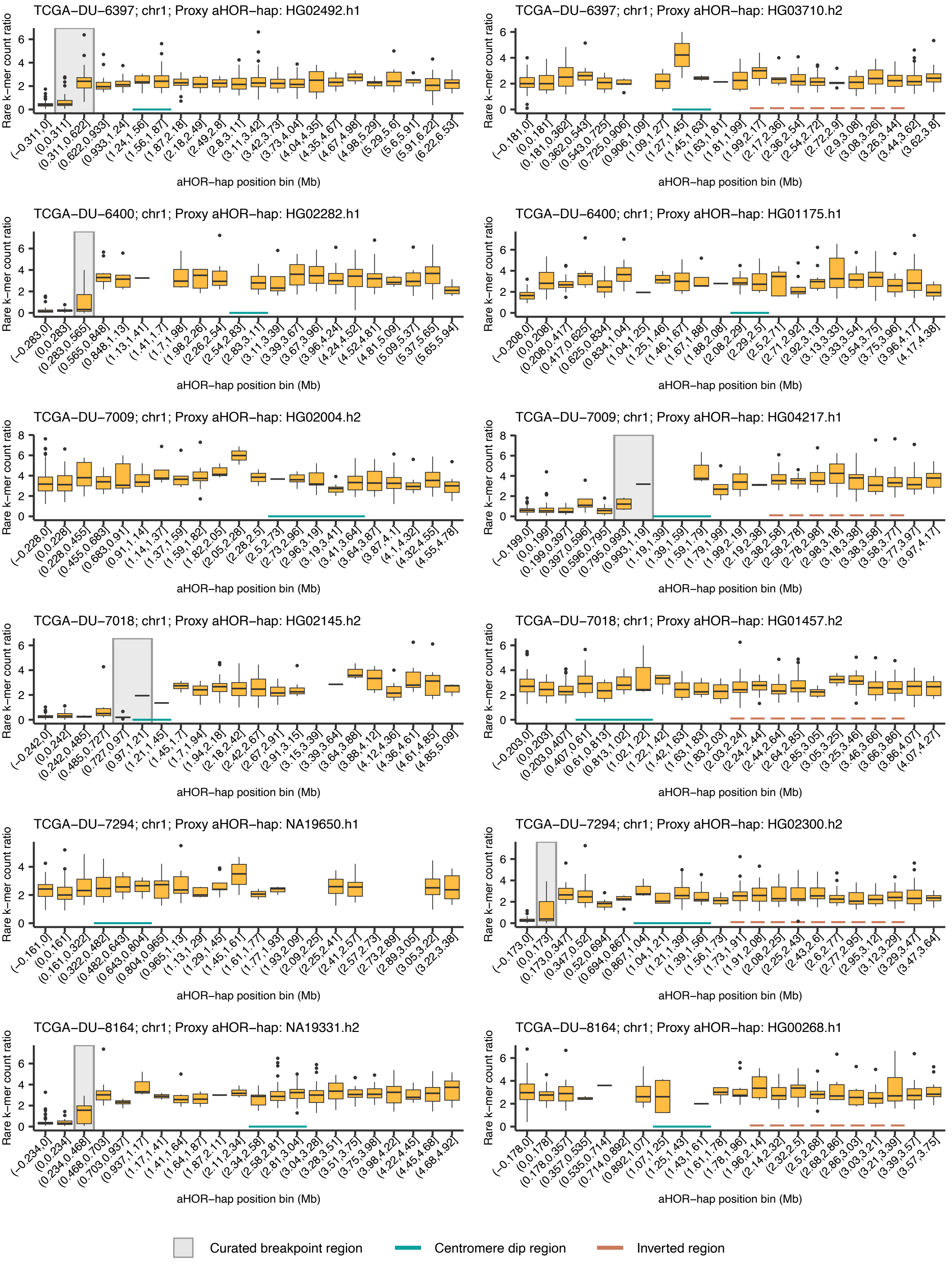


**Supplementary Figure. 6 (continued)**

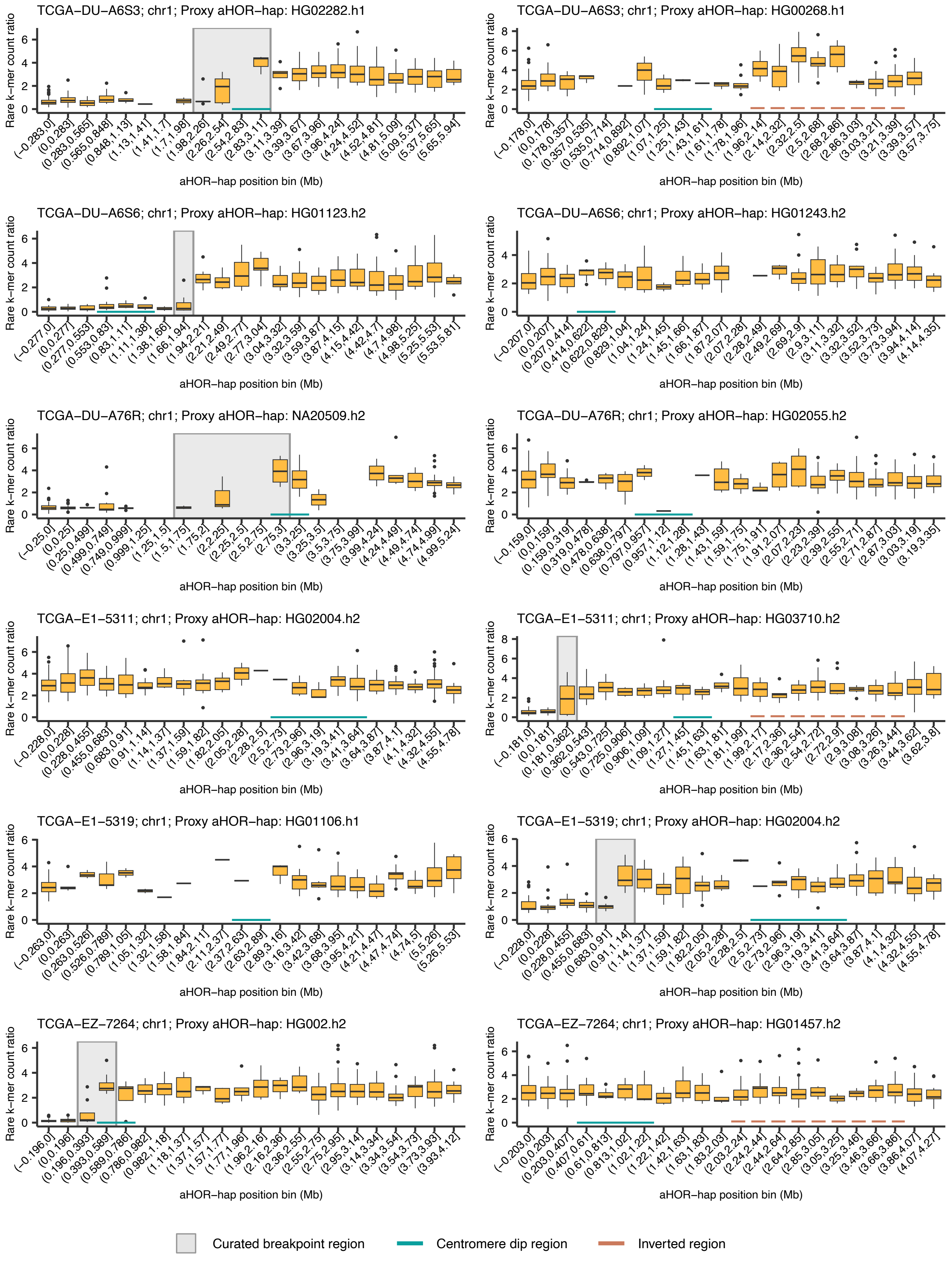


**Supplementary Figure. 6 (continued)**

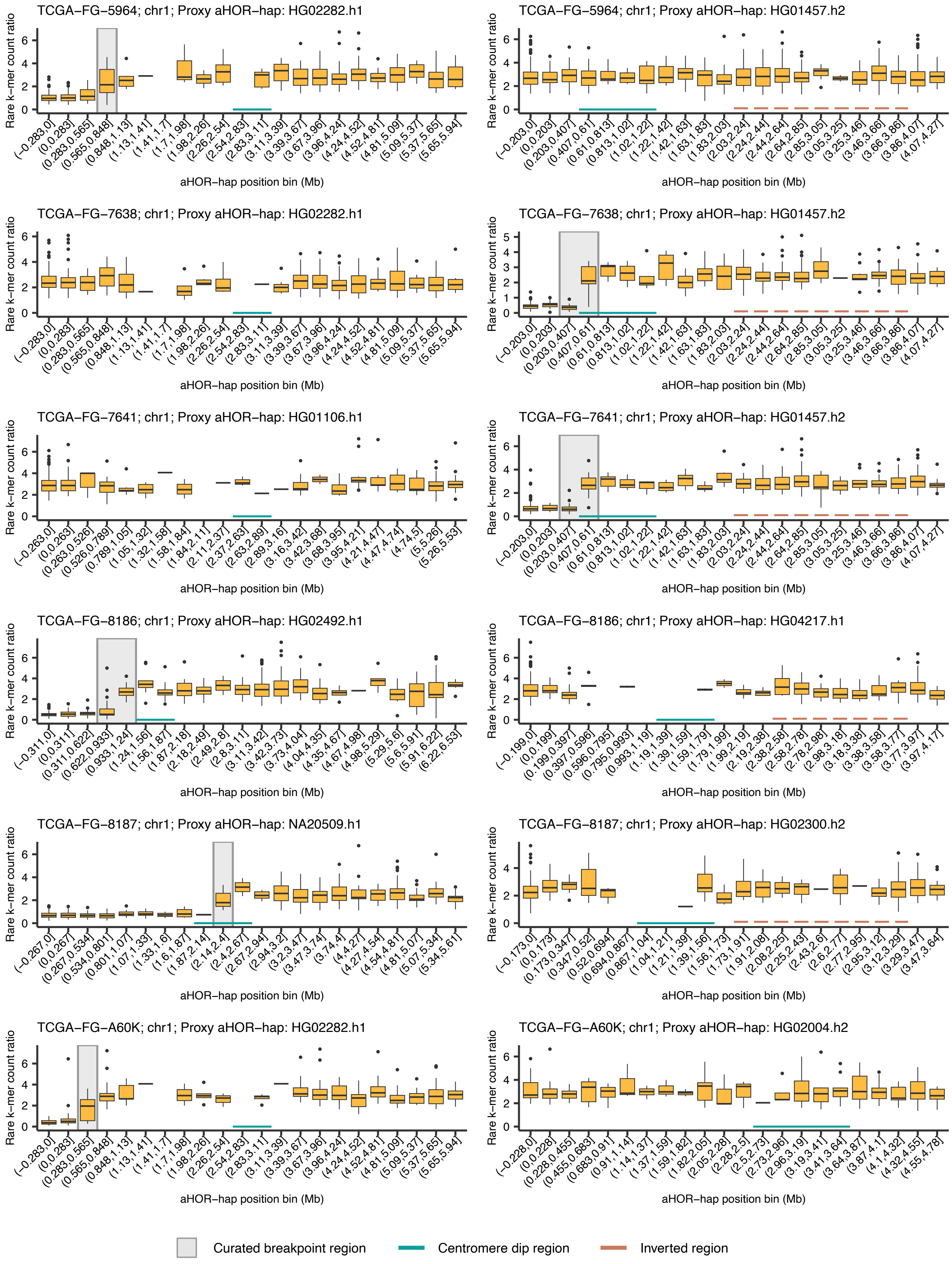


**Supplementary Figure. 6 (continued)**

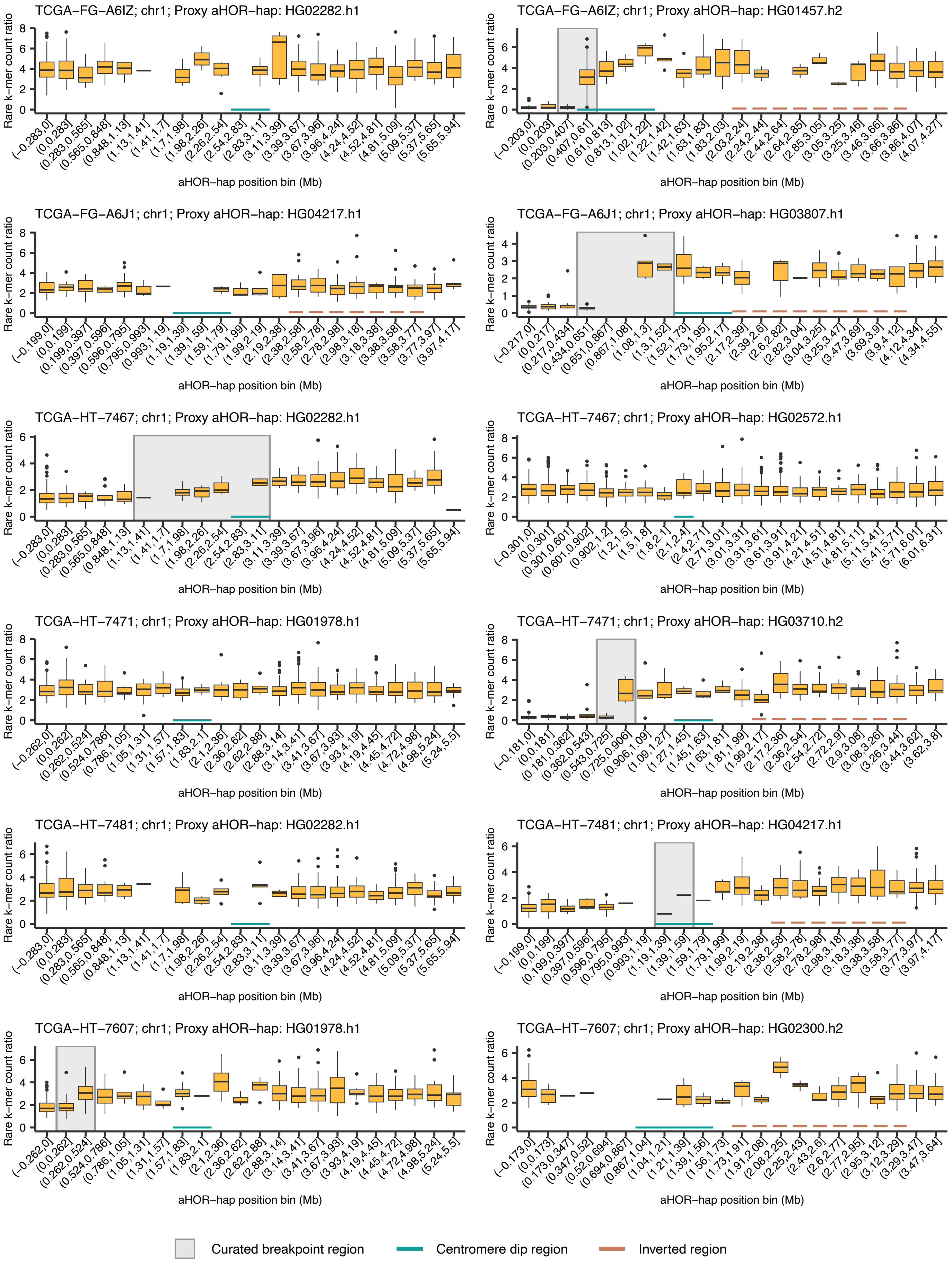


**Supplementary Figure. 6 (continued)**

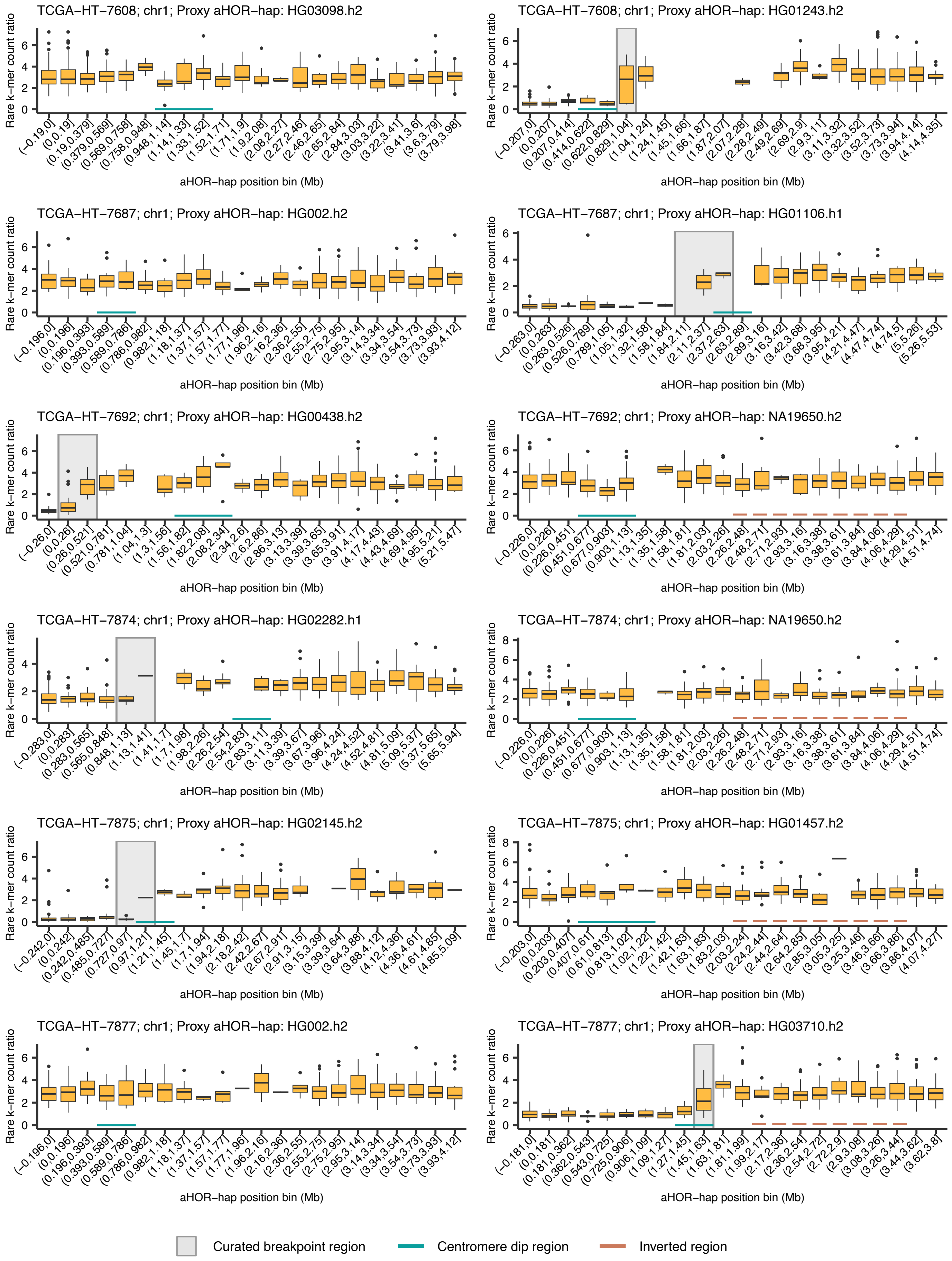


**Supplementary Figure. 6 (continued)**

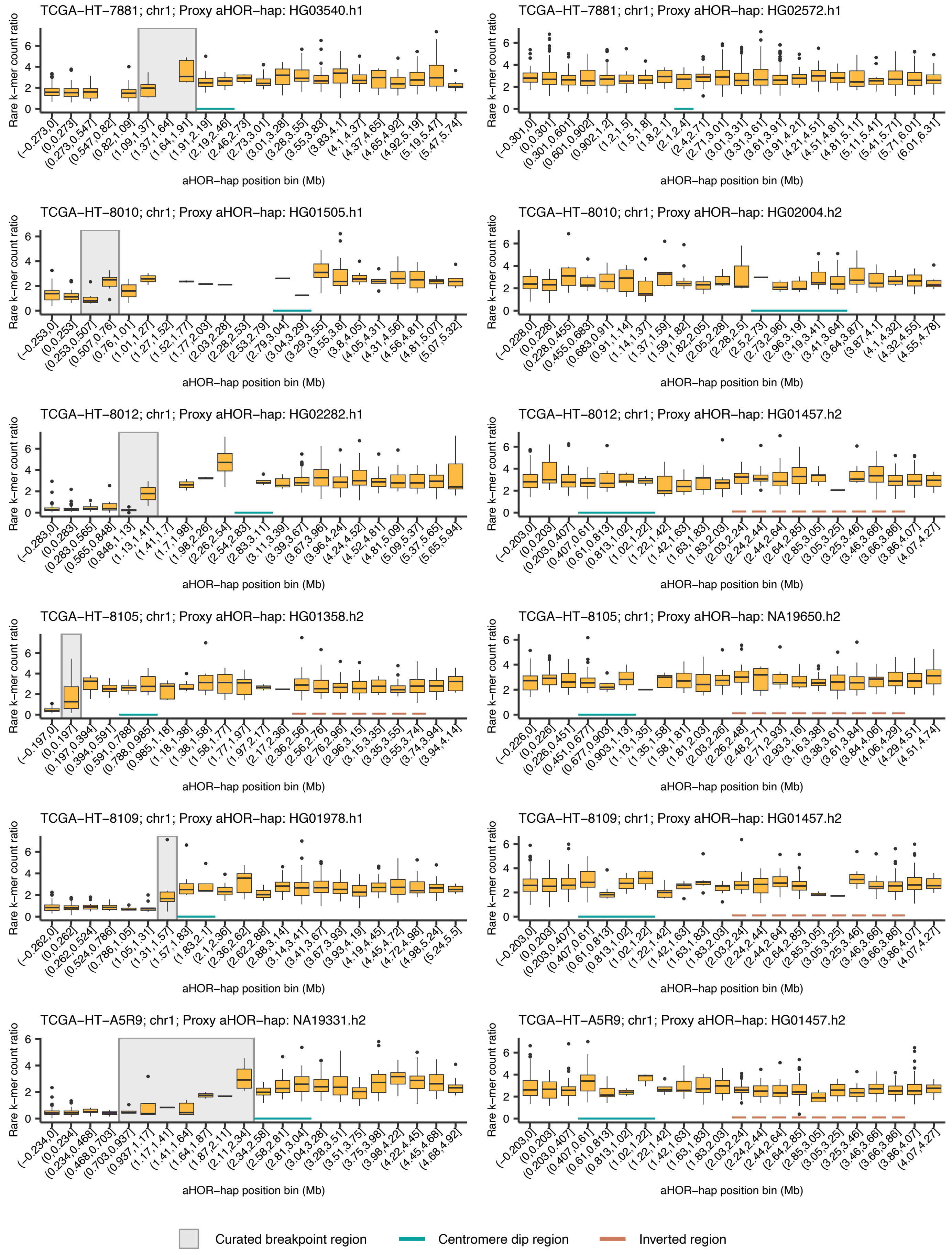


**Supplementary Figure. 6 (continued)**

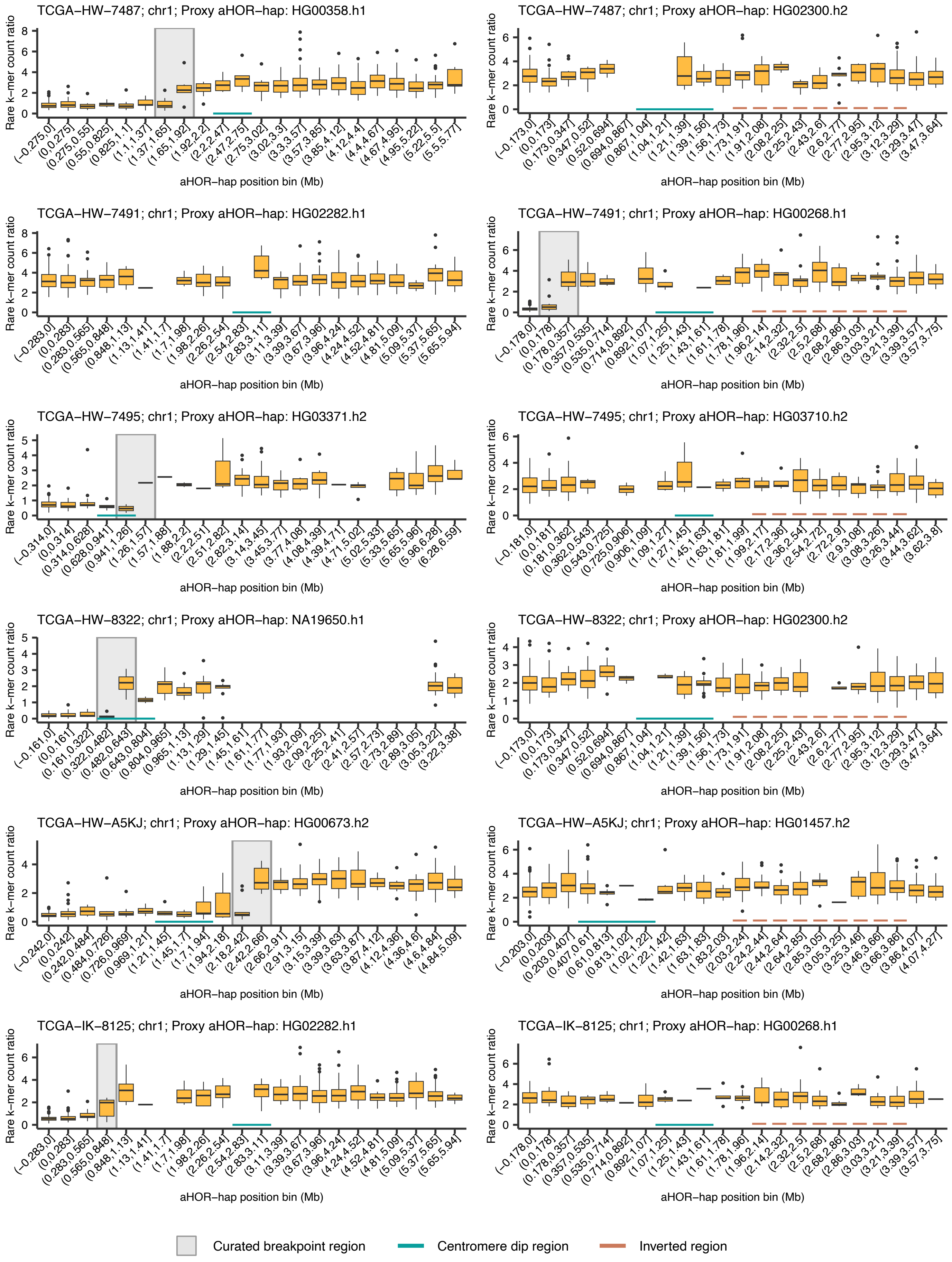


**Supplementary Figure. 6 (continued)**

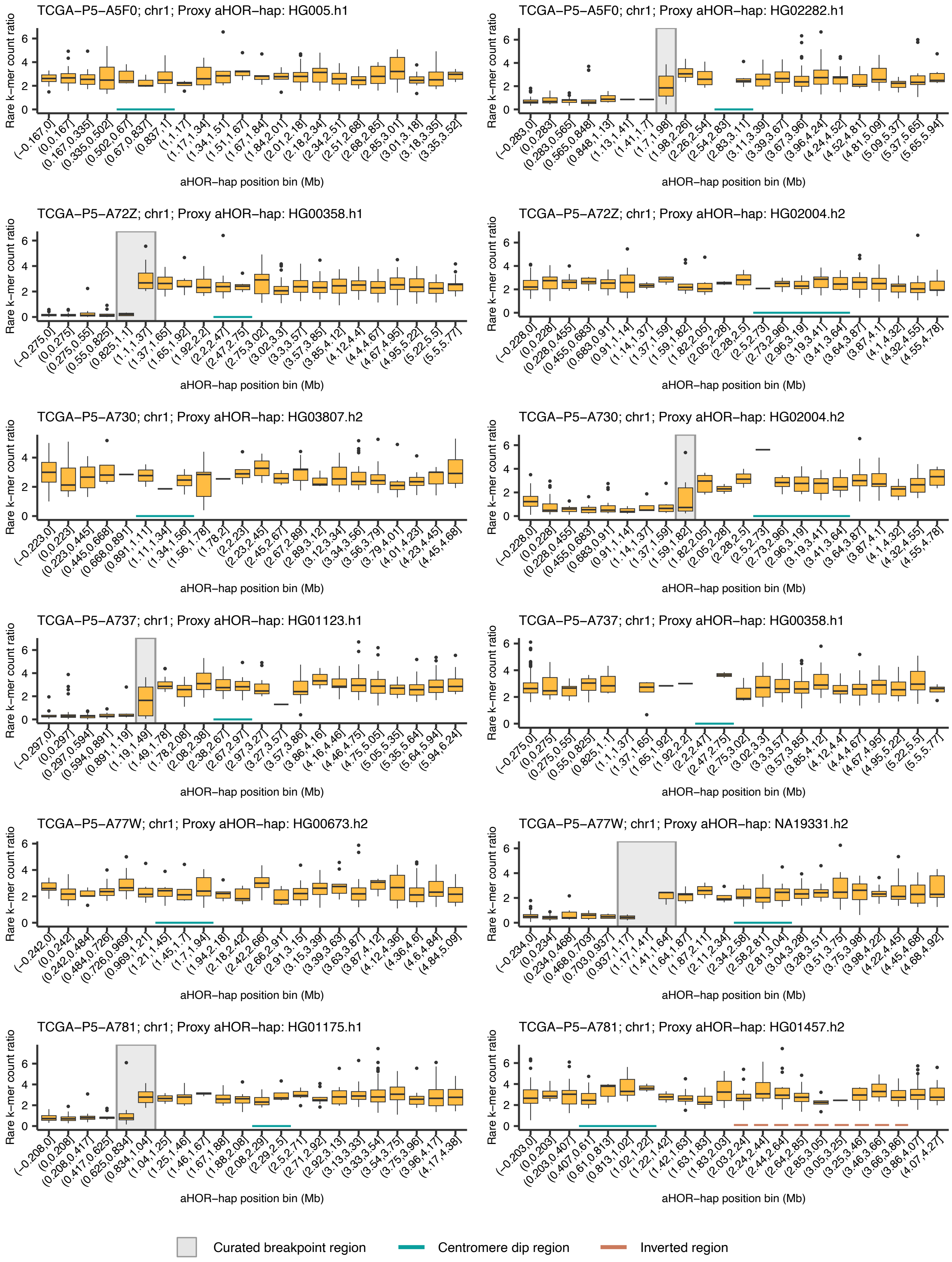


**Supplementary Figure. 6 (continued)**

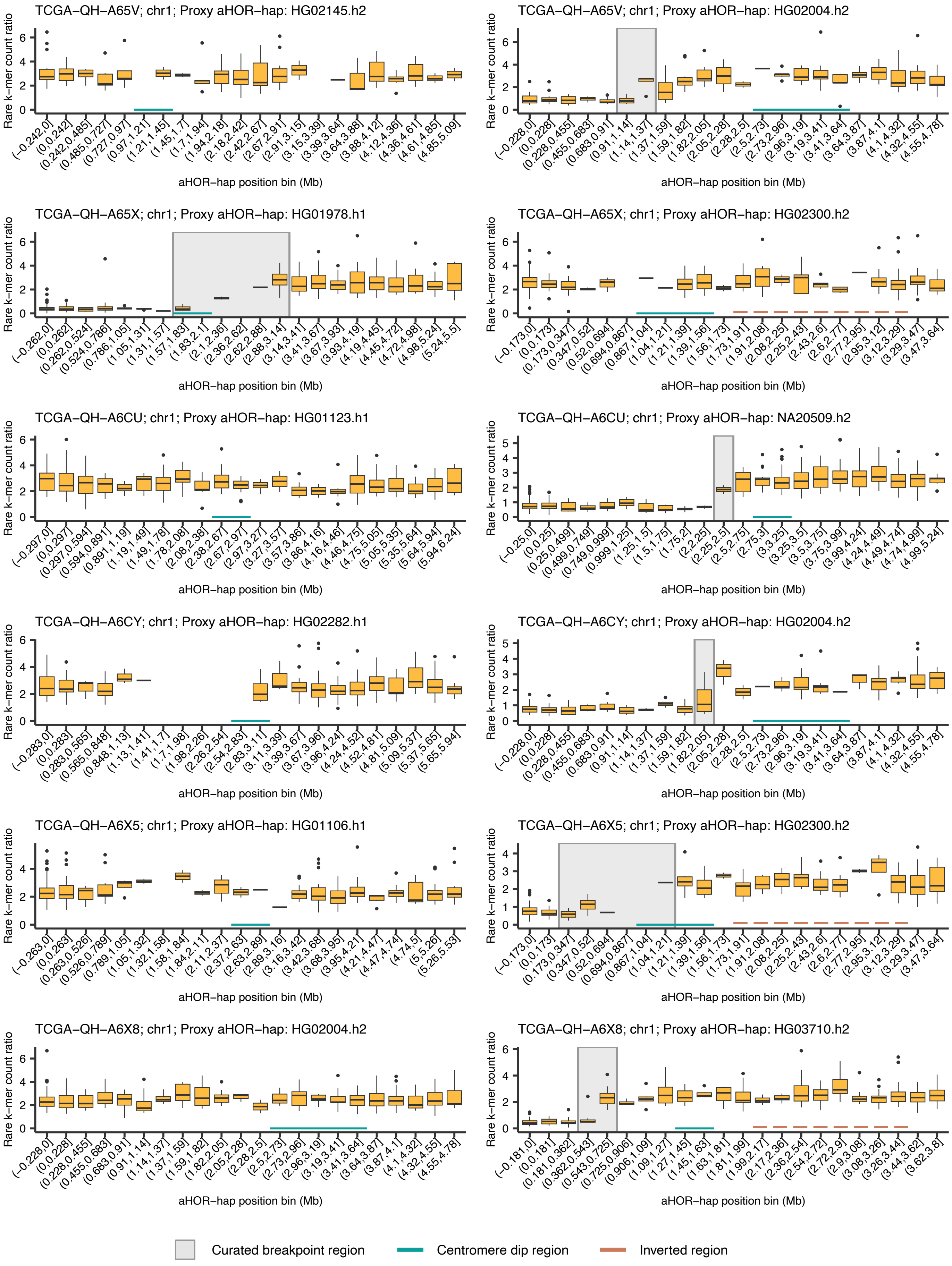


**Supplementary Figure. 6 (continued)**

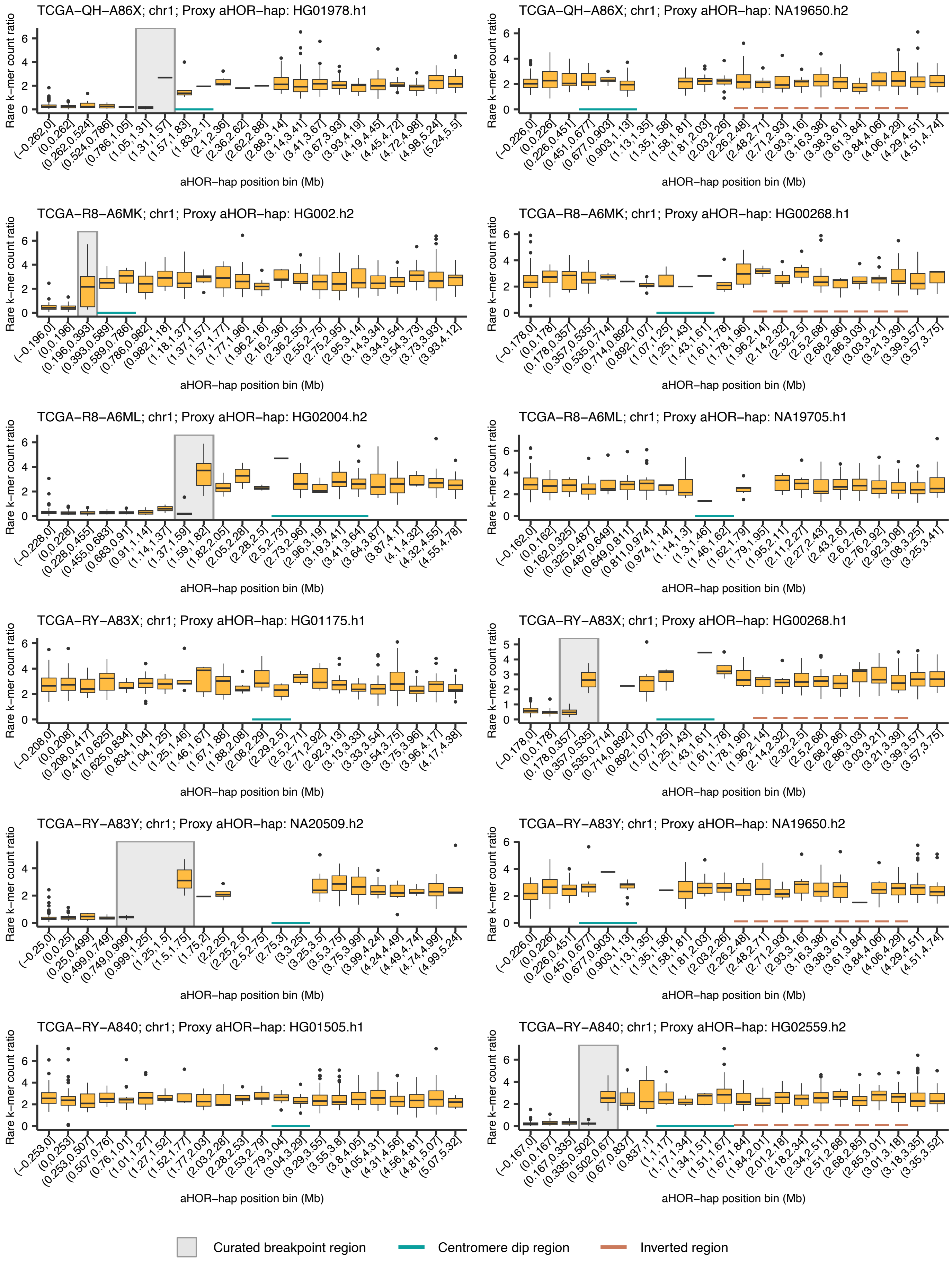


**Supplementary Figure. 6 (continued)**

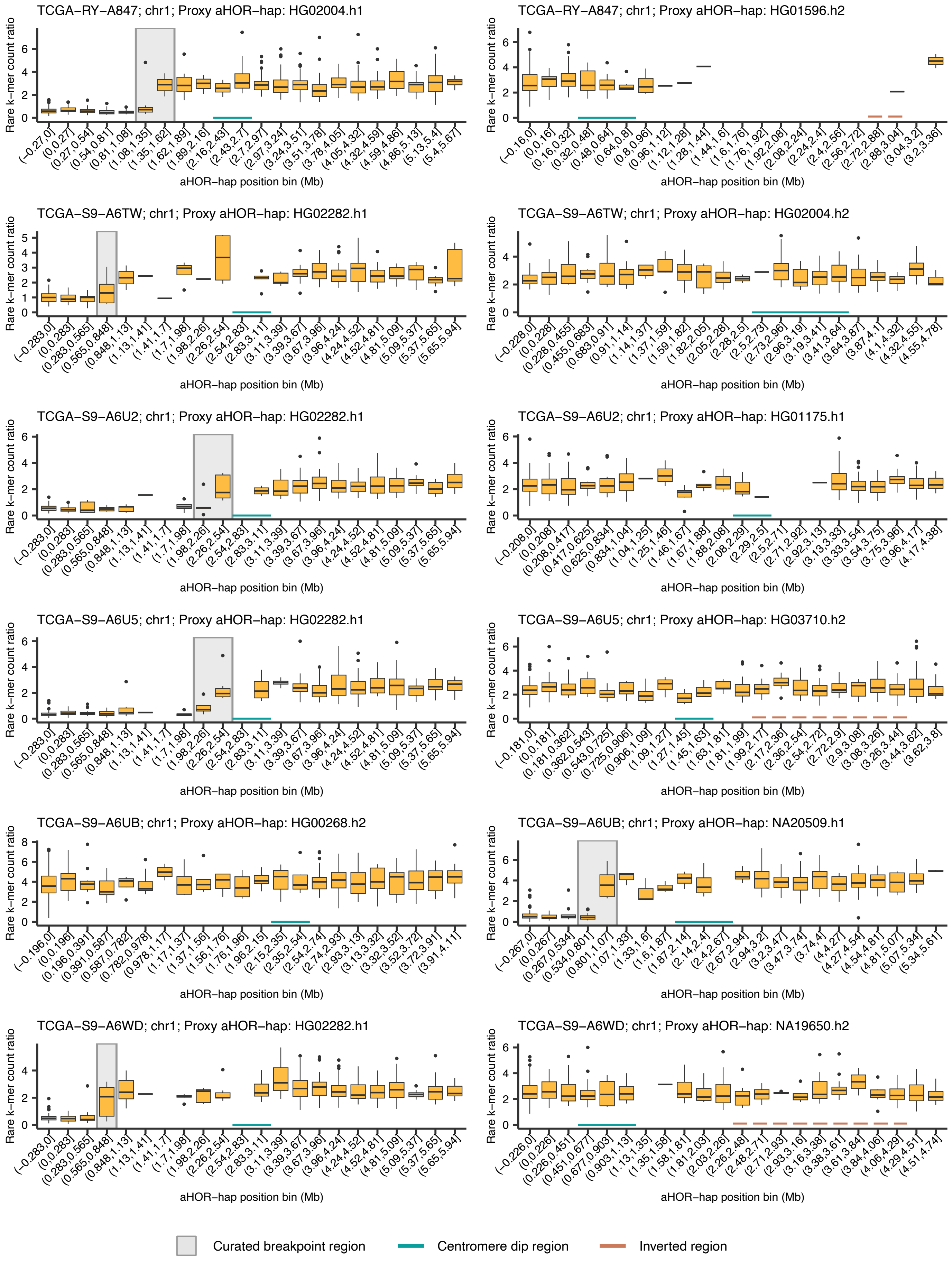


**Supplementary Figure. 6 (continued)**

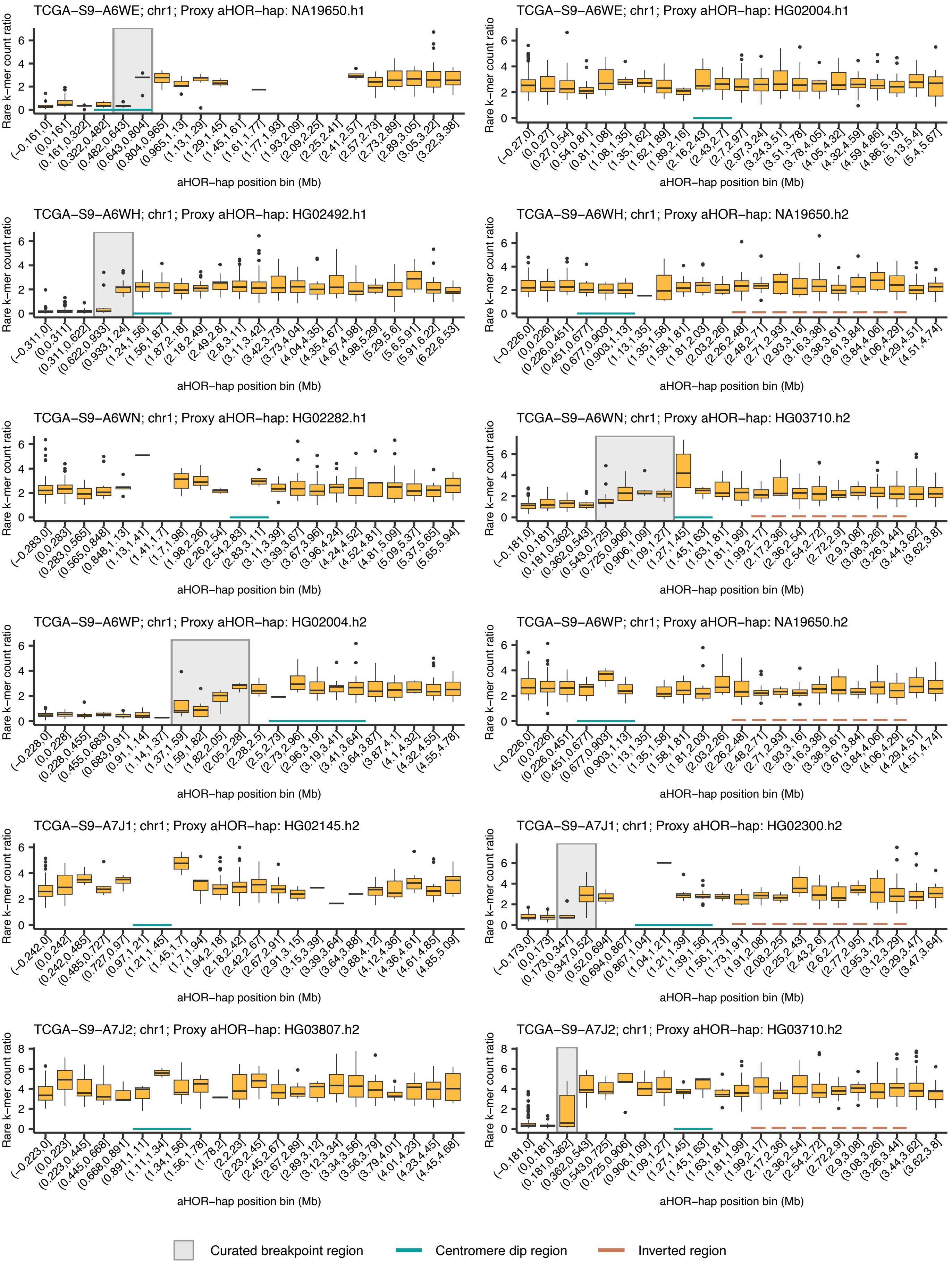


**Supplementary Figure. 6 (continued)**

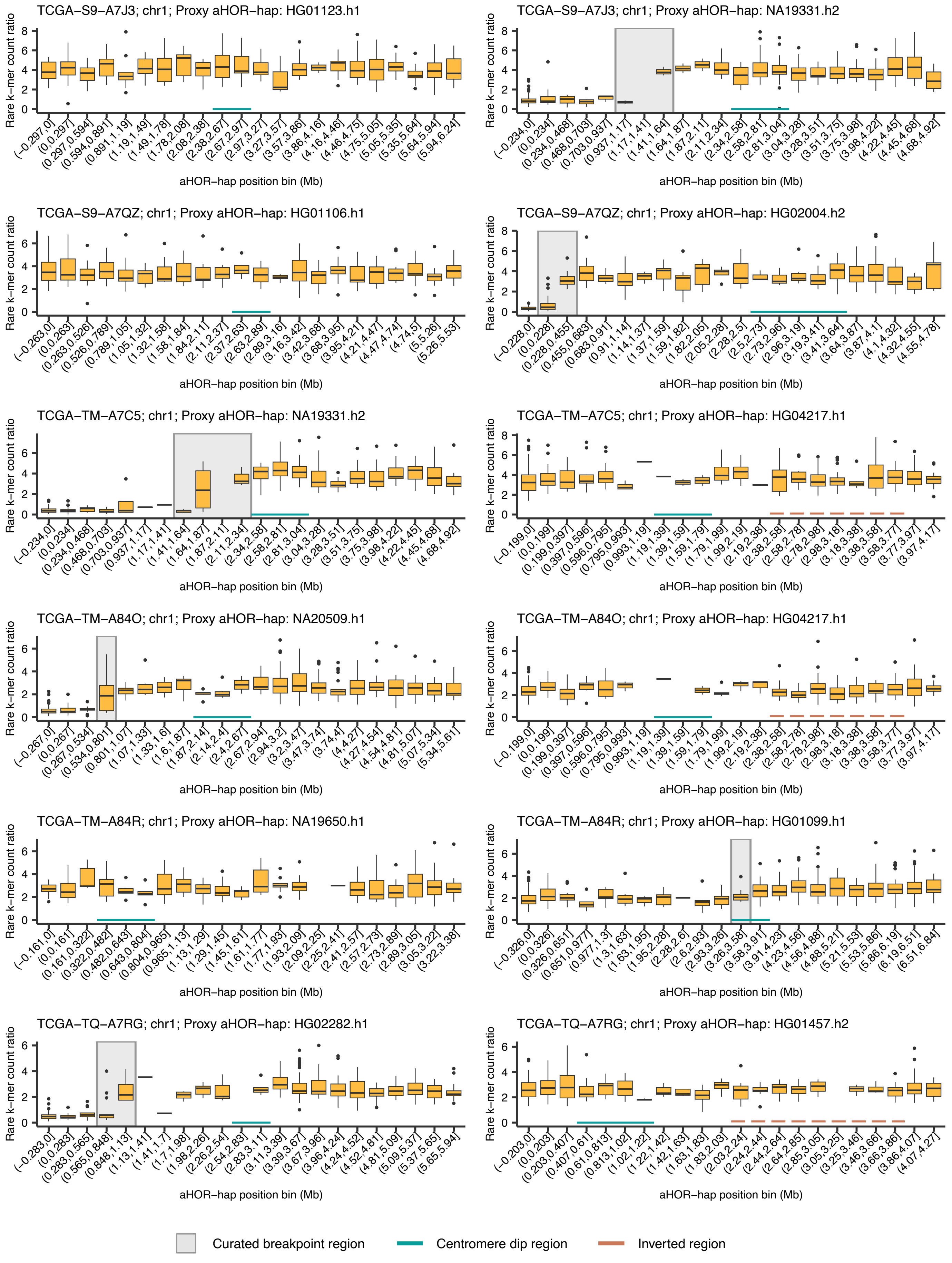


**Supplementary Figure. 6 (continued)**

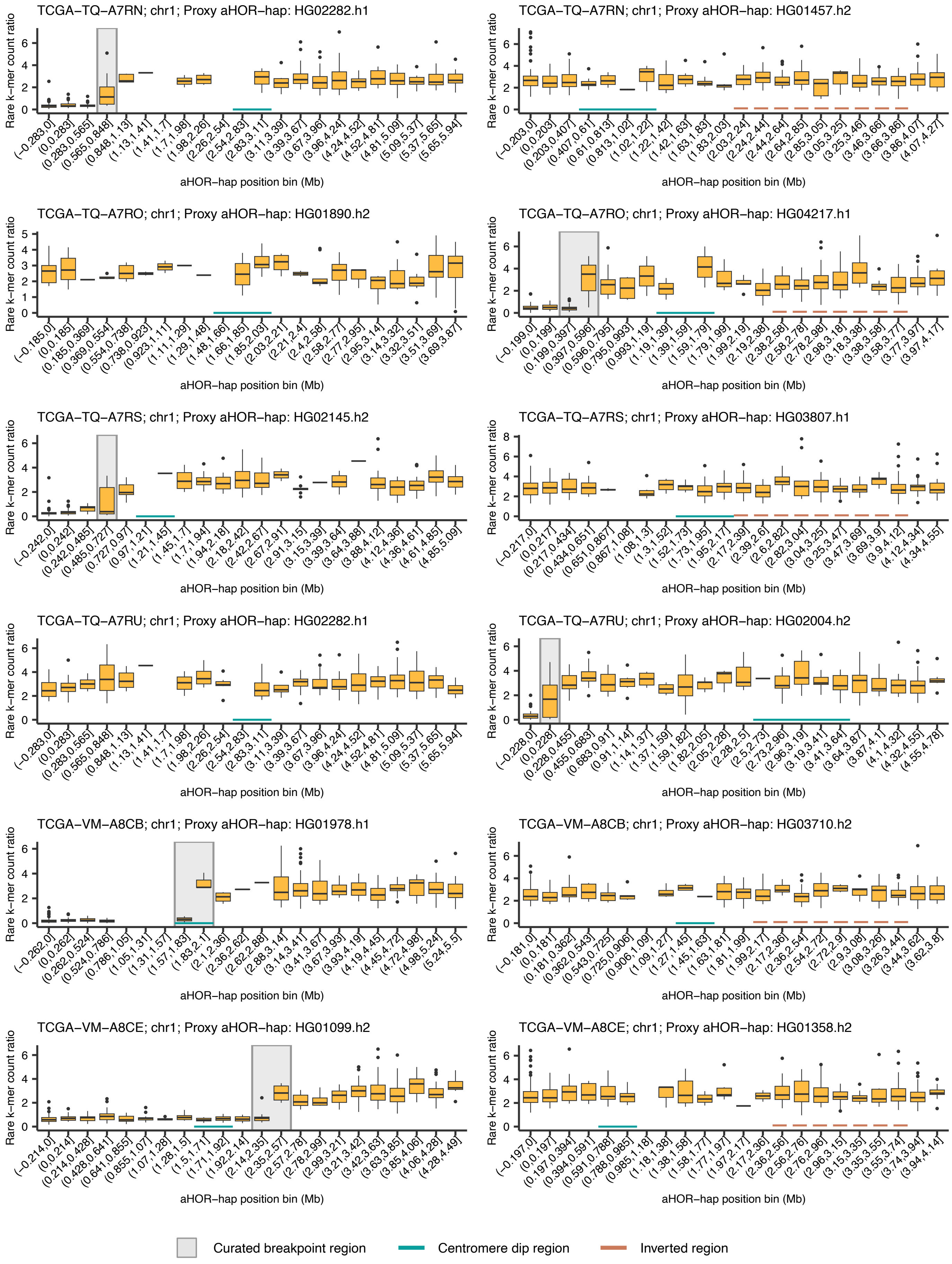


**Supplementary Figure. 6 (continued)**

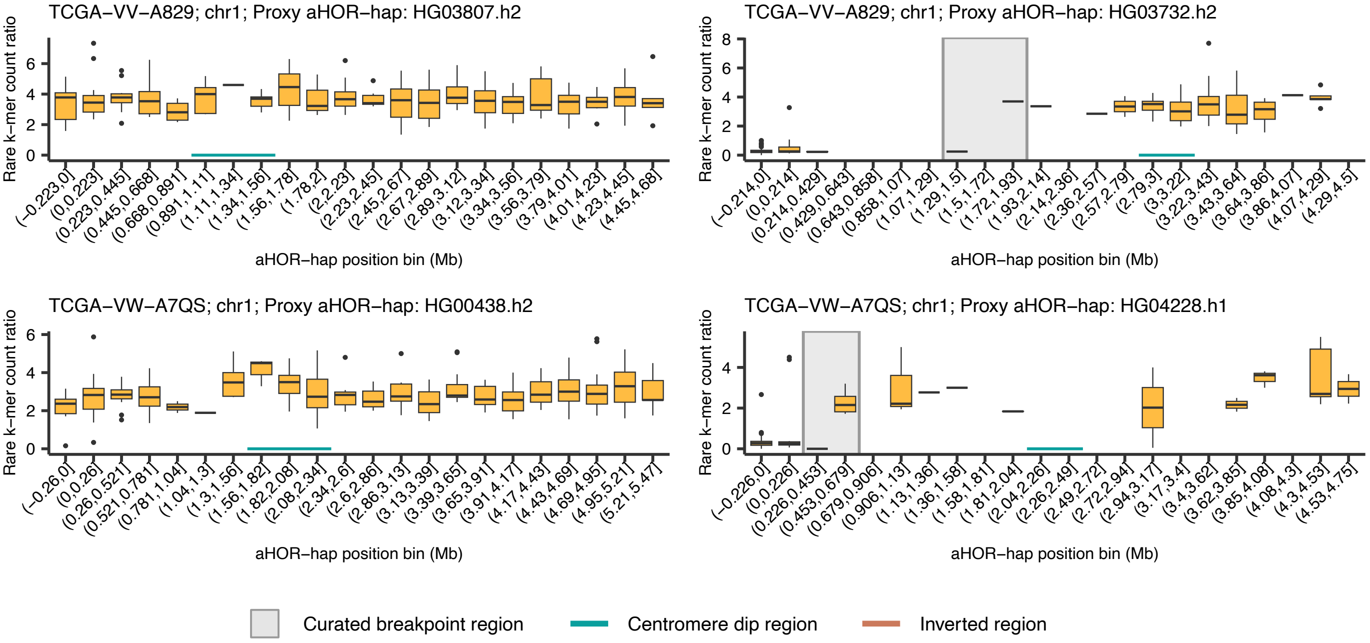


**Supplementary Figure. 6 (continued)**

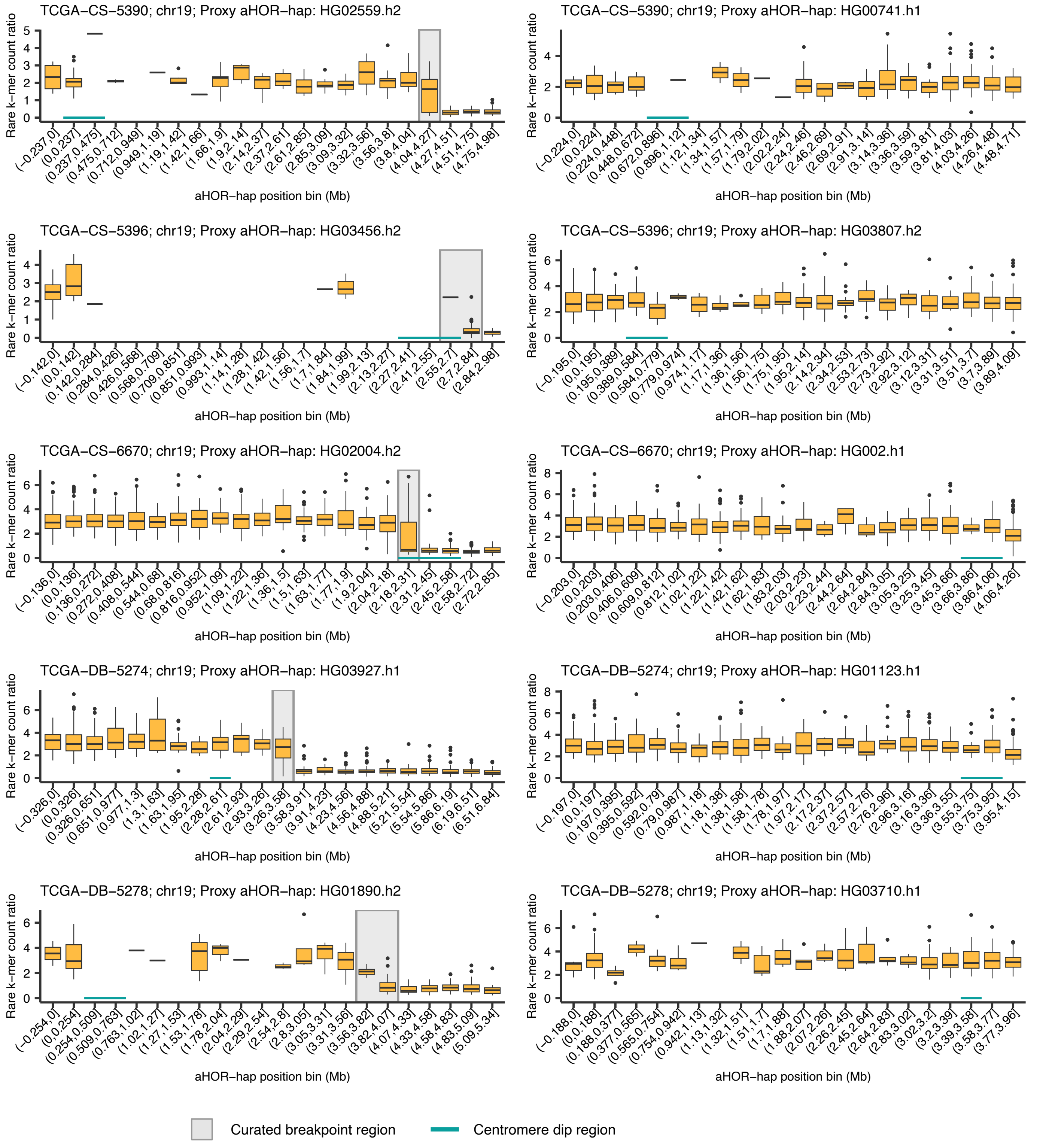


**Supplementary Figure. 7: Copy number profiles at the *D19Z3* array regions computed from 118 TCGA low-grade glioma tumor and matched normal WGS data.** Boxplots show the rare k-mer count ratio between tumor and normal samples for each positional bin along the proxy aHOR-haps (in megabases [Mb]). Shaded gray areas highlight inferred breakpoint locations of rearrangements identified by copy number alterations. Cyan horizontal bars beneath the boxplots denote centromere dip regions (CDRs), indicating kinetochore attachment sites.

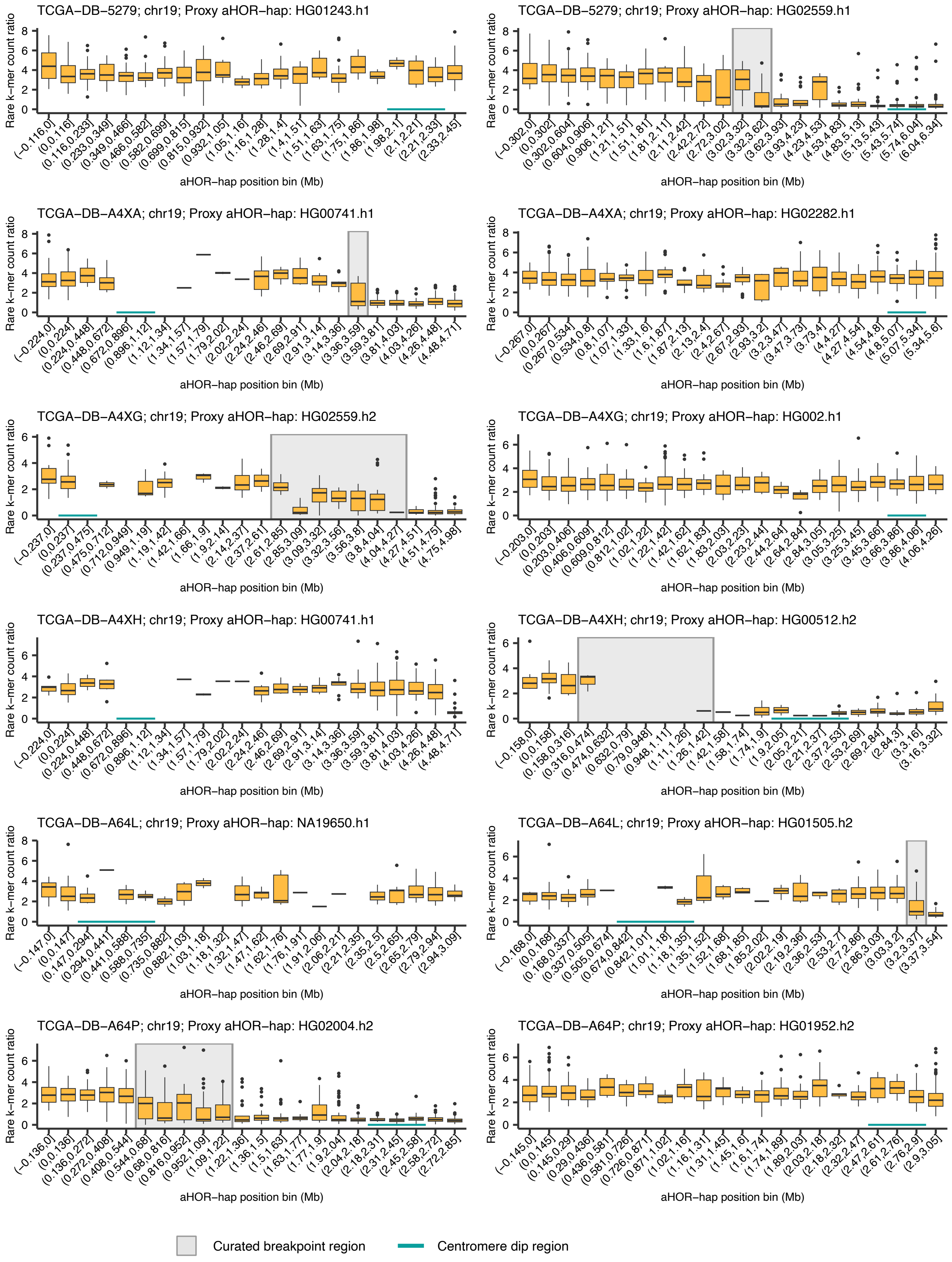


**Supplementary Figure. 7 (continued)**

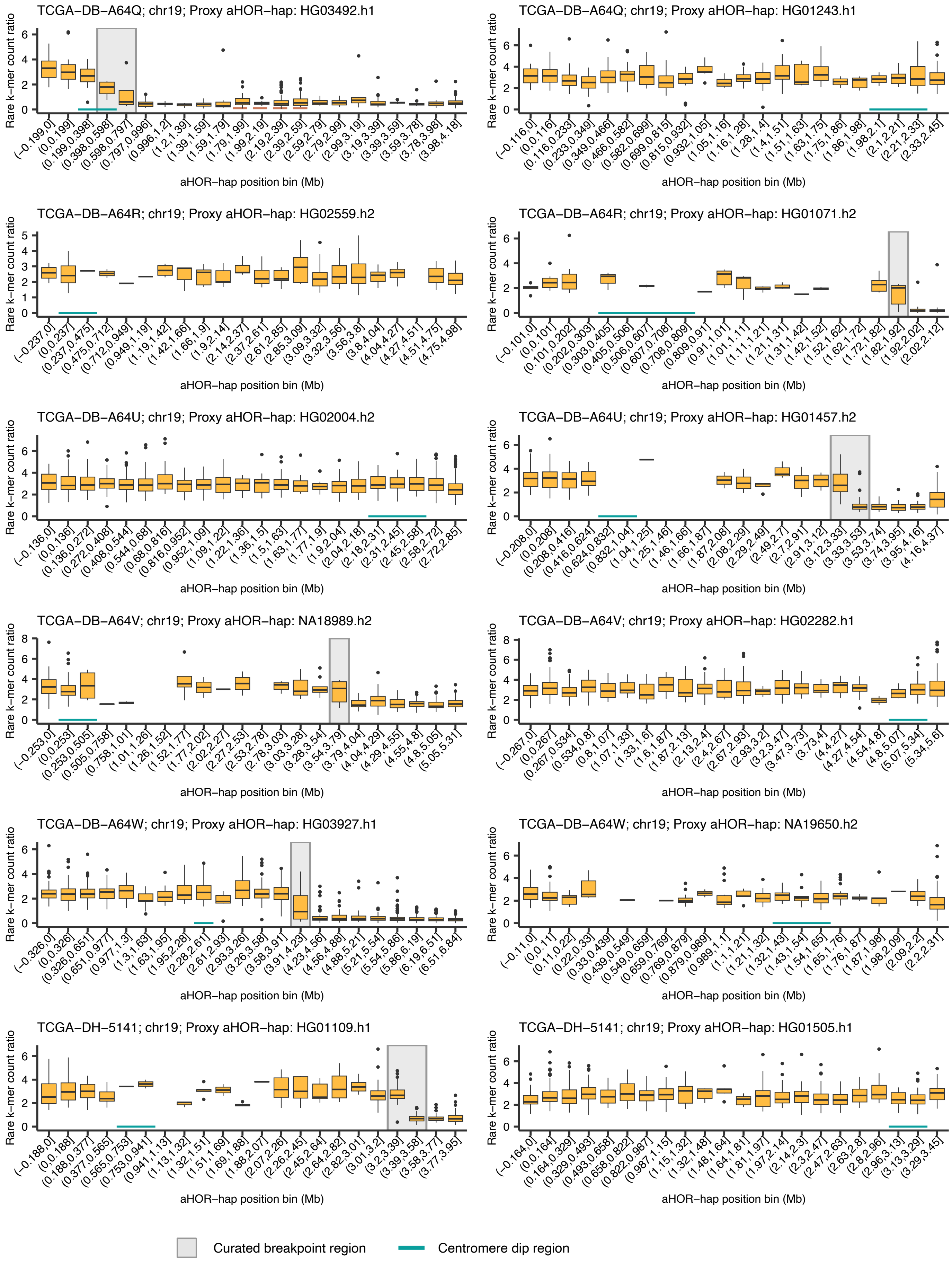


**Supplementary Figure. 7 (continued)**

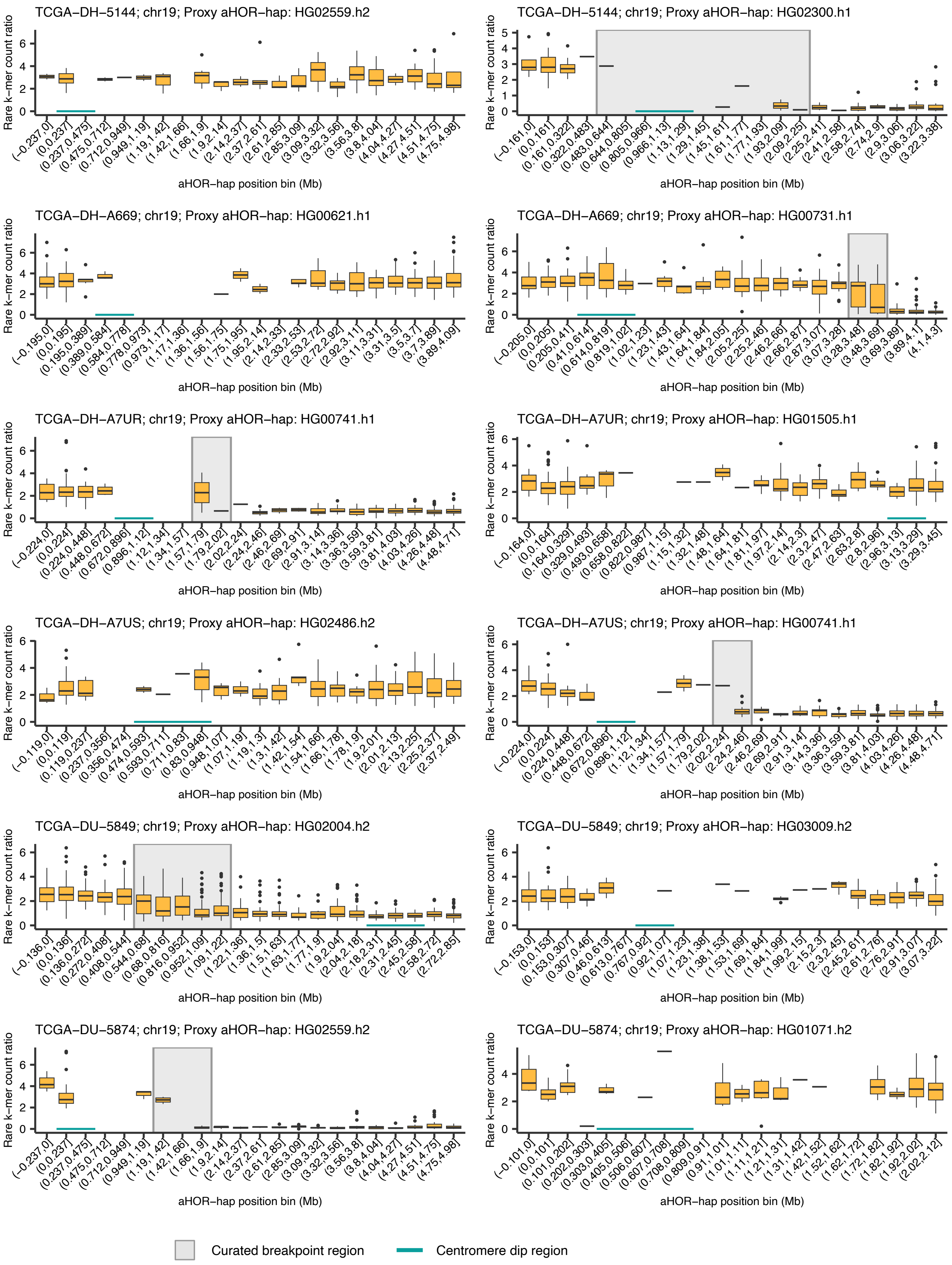


**Supplementary Figure. 7 (continued)**

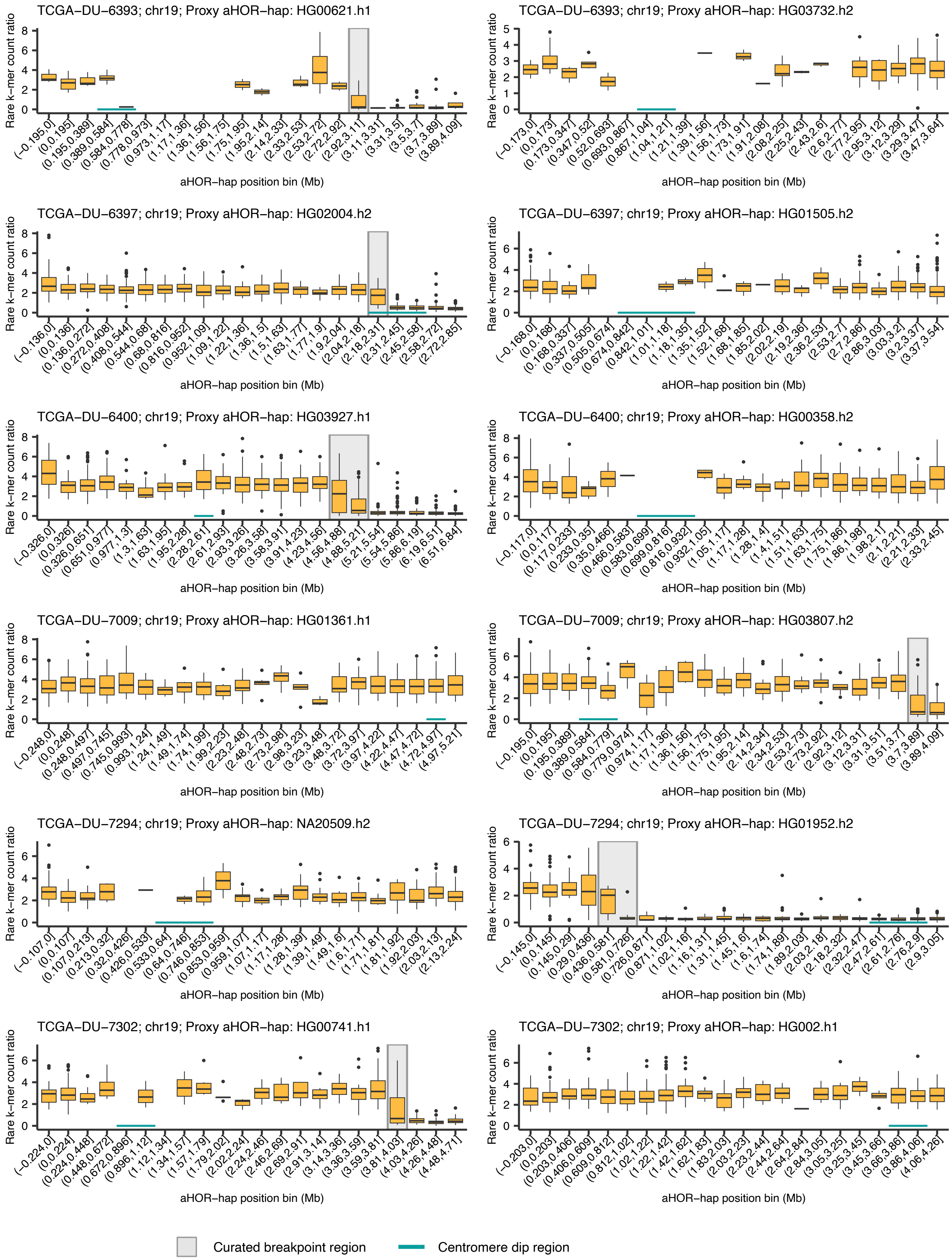


**Supplementary Figure. 7 (continued)**

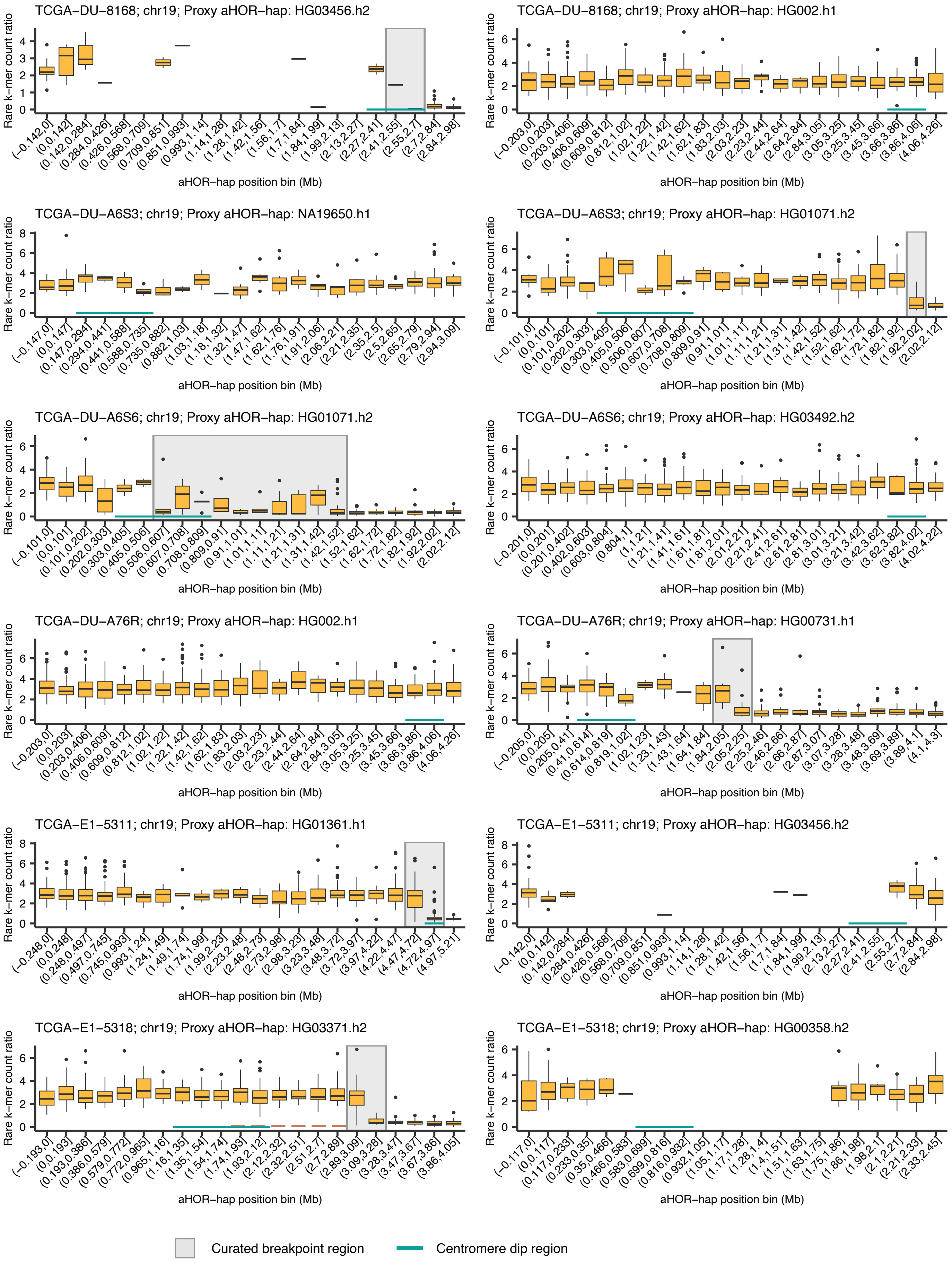


**Supplementary Figure. 7 (continued)**

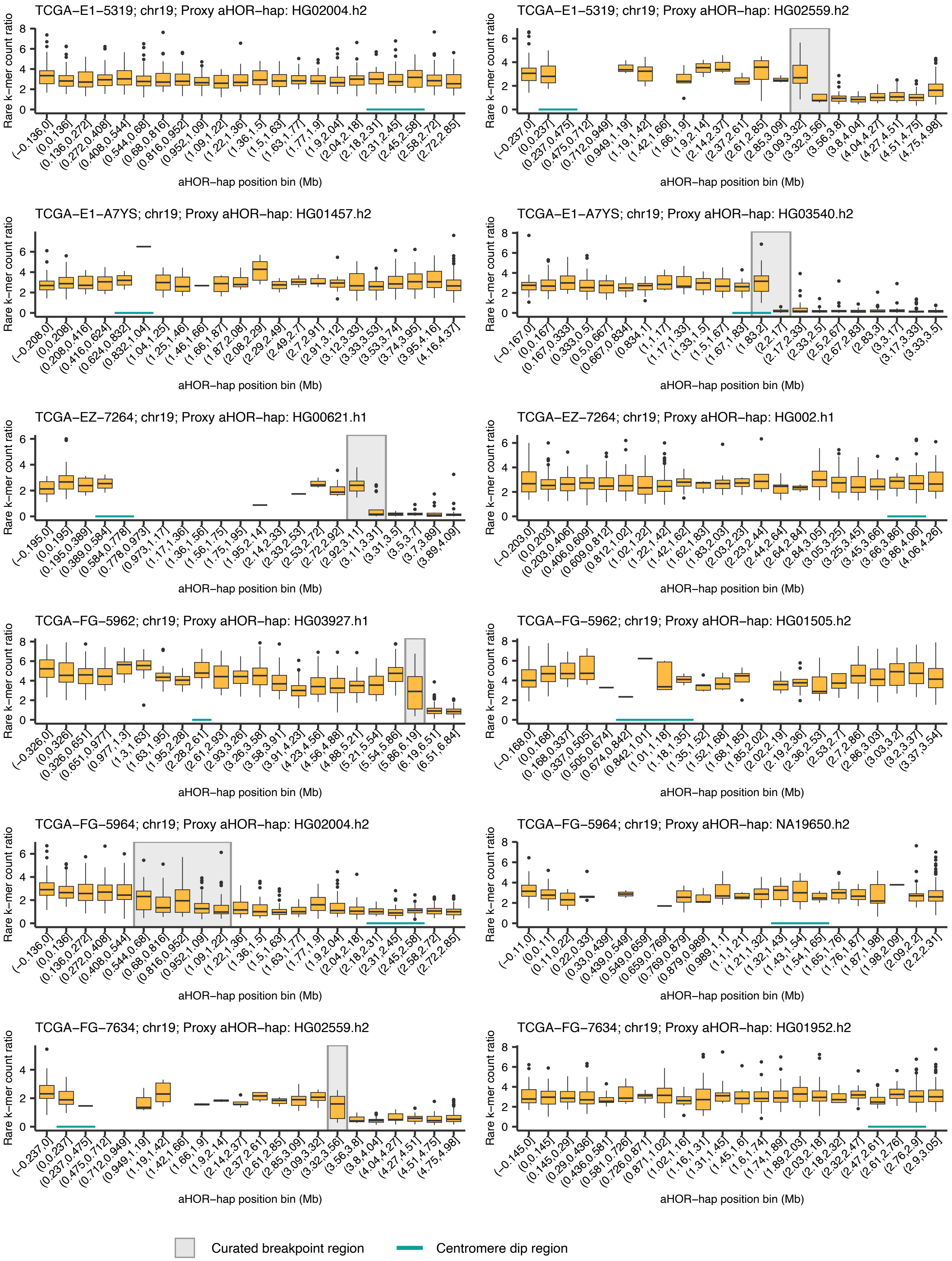


**Supplementary Figure. 7 (continued)**

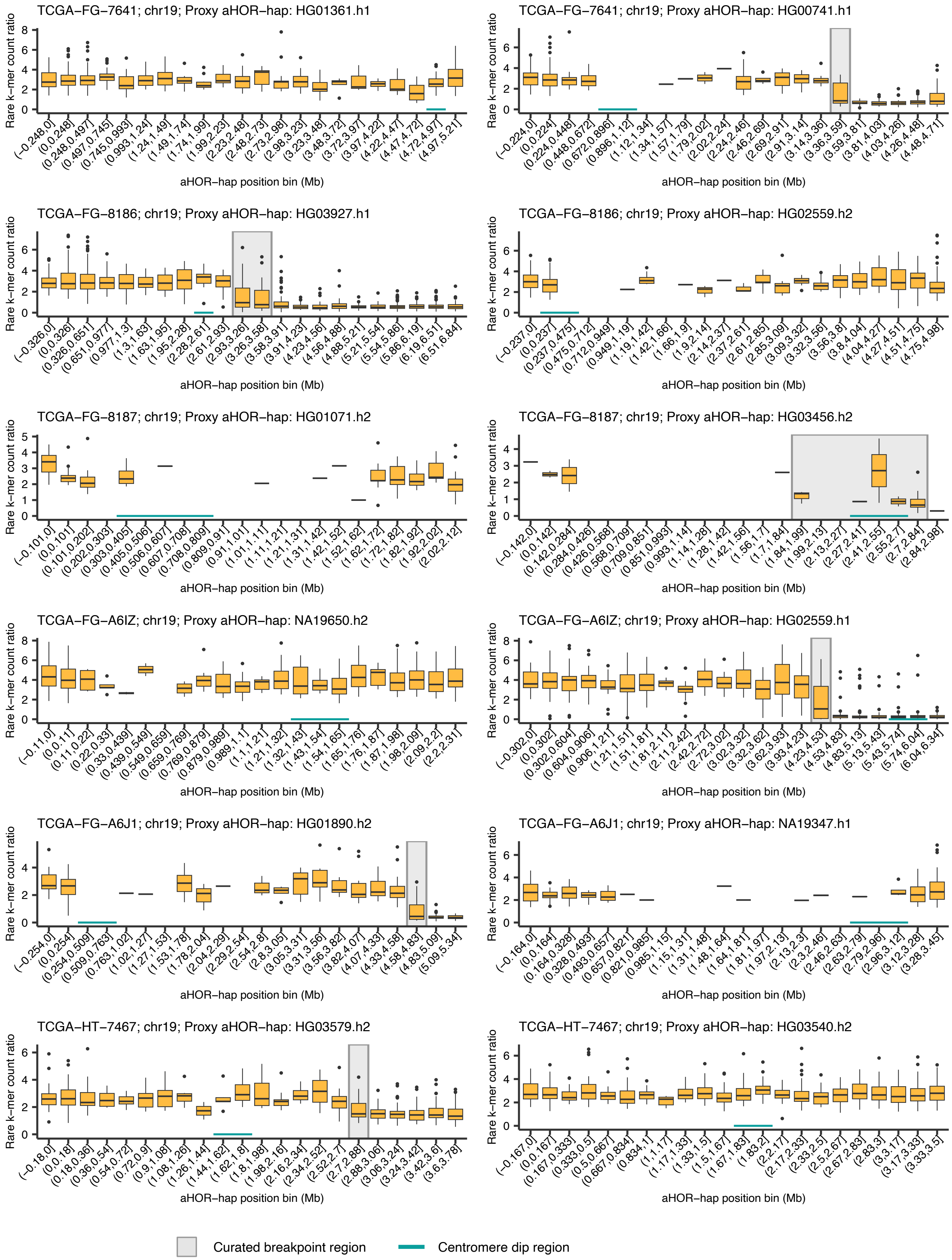


**Supplementary Figure. 7 (continued)**

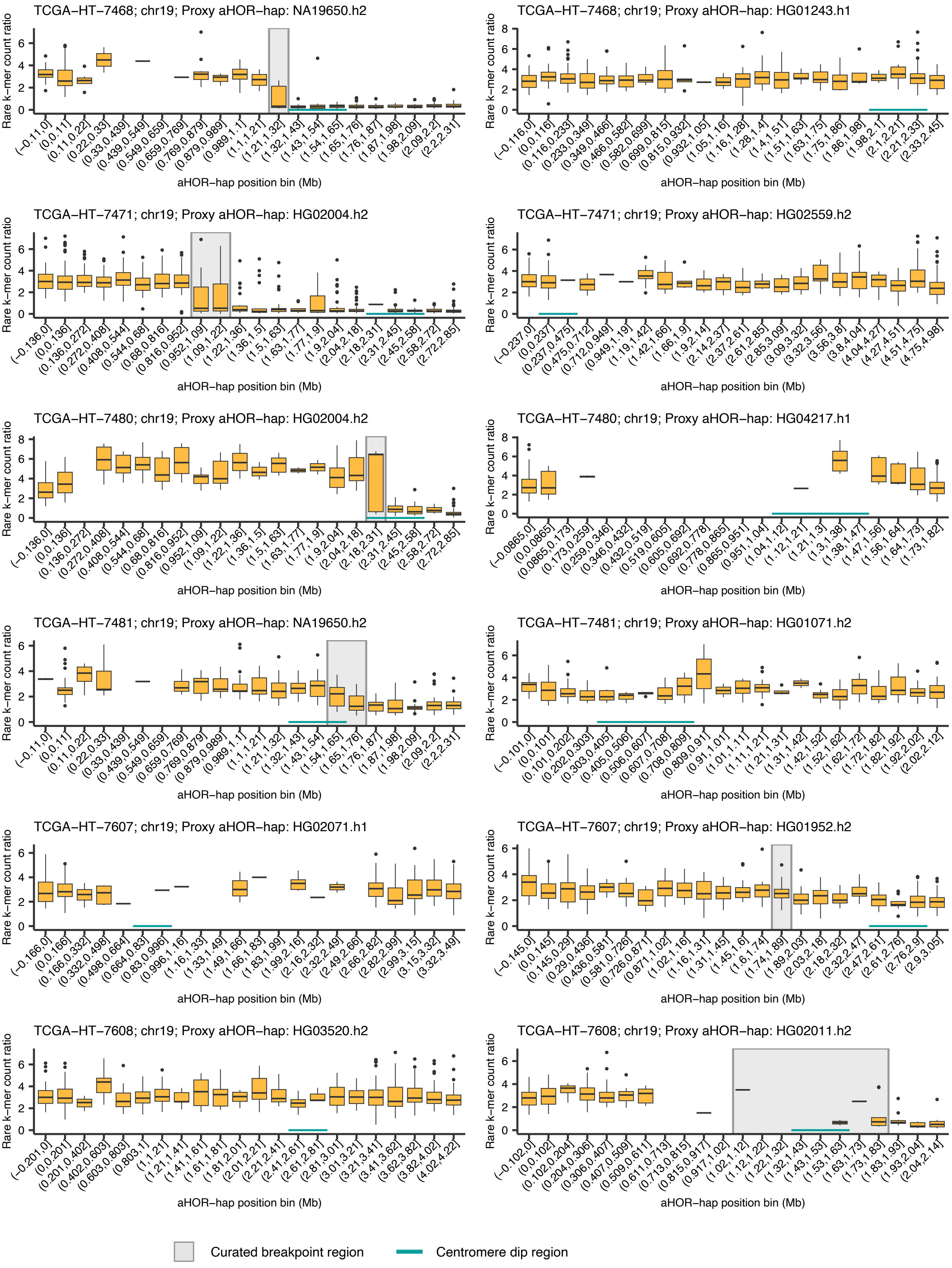


**Supplementary Figure. 7 (continued)**

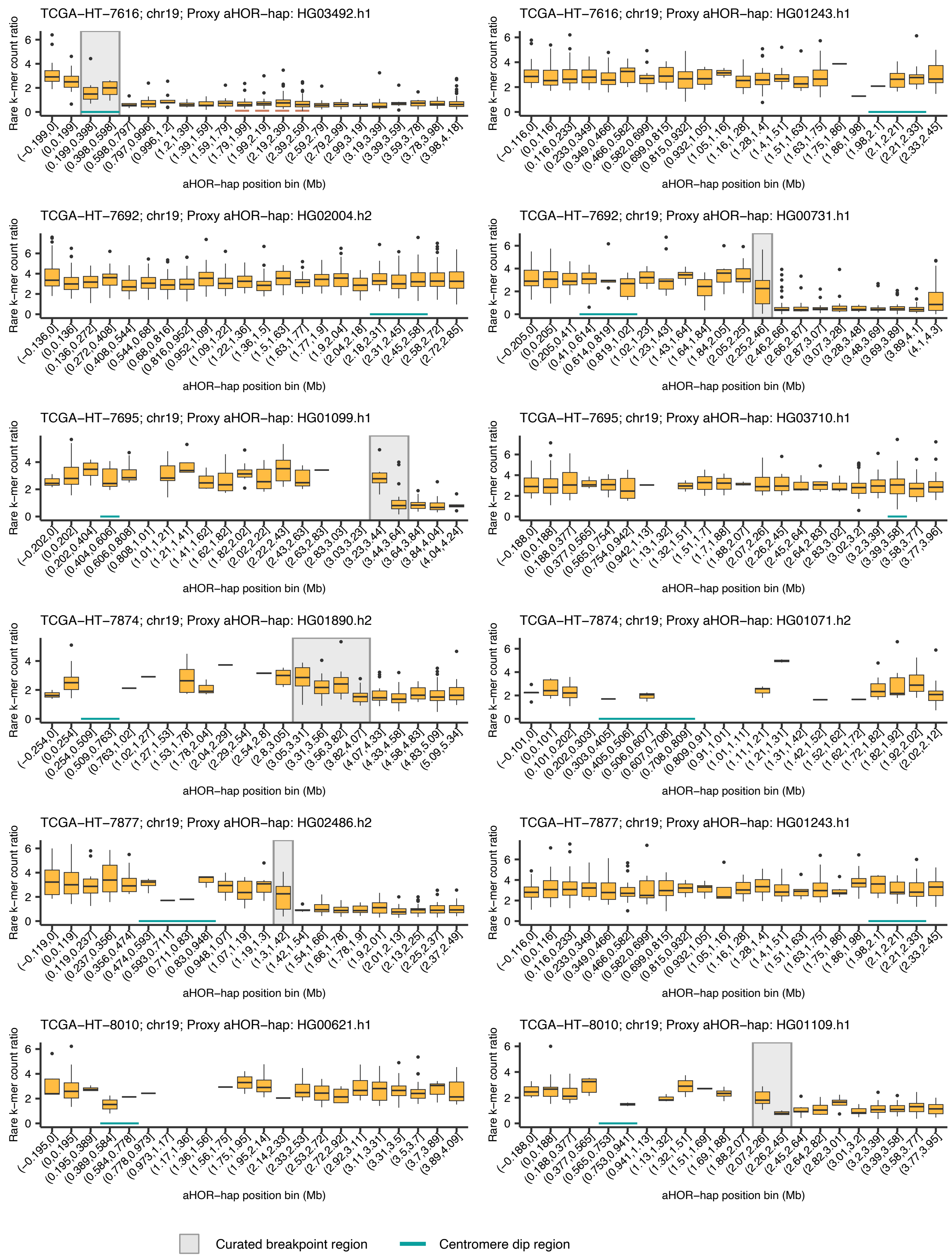


**Supplementary Figure. 7 (continued)**

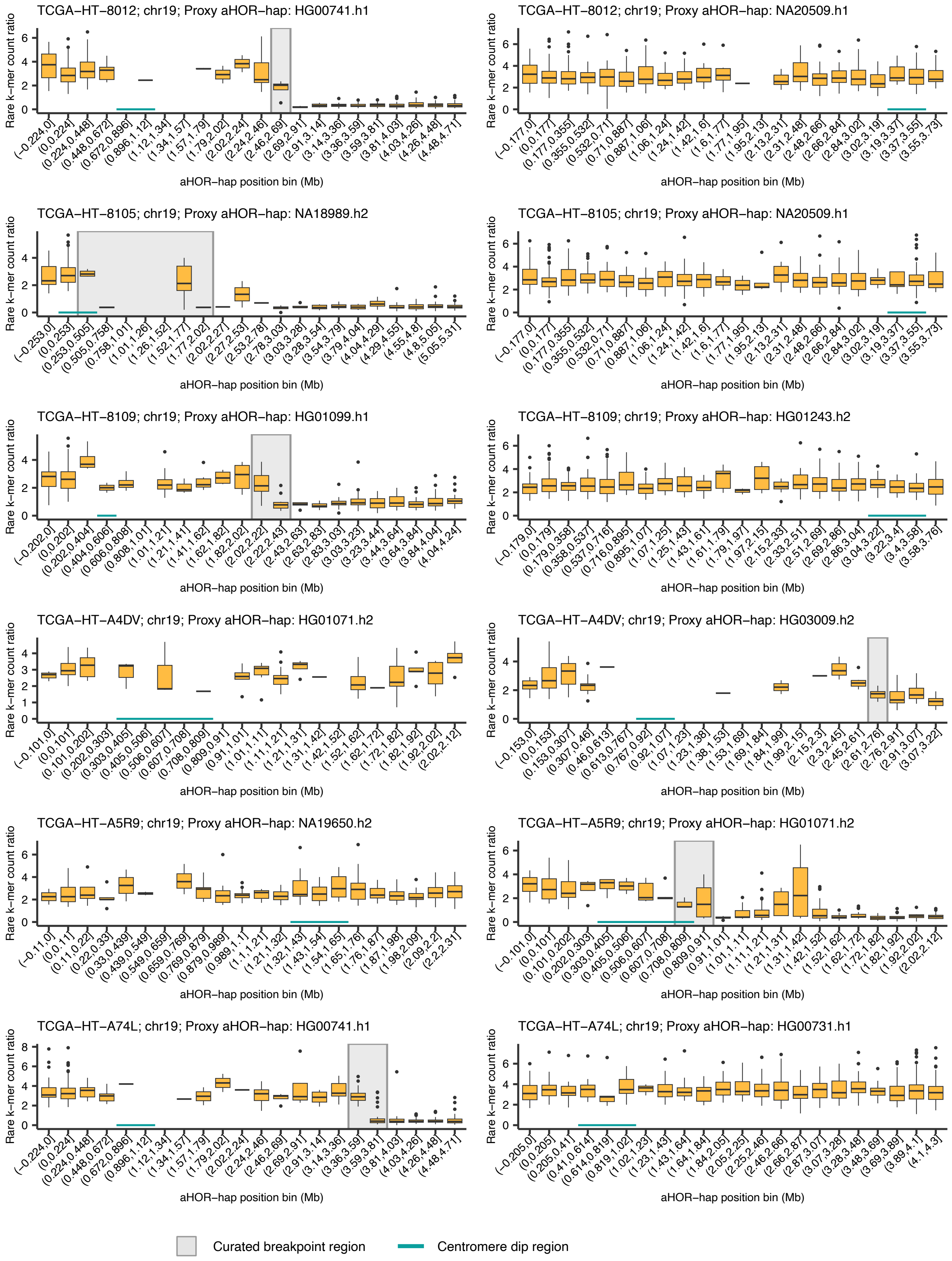


**Supplementary Figure. 7 (continued)**

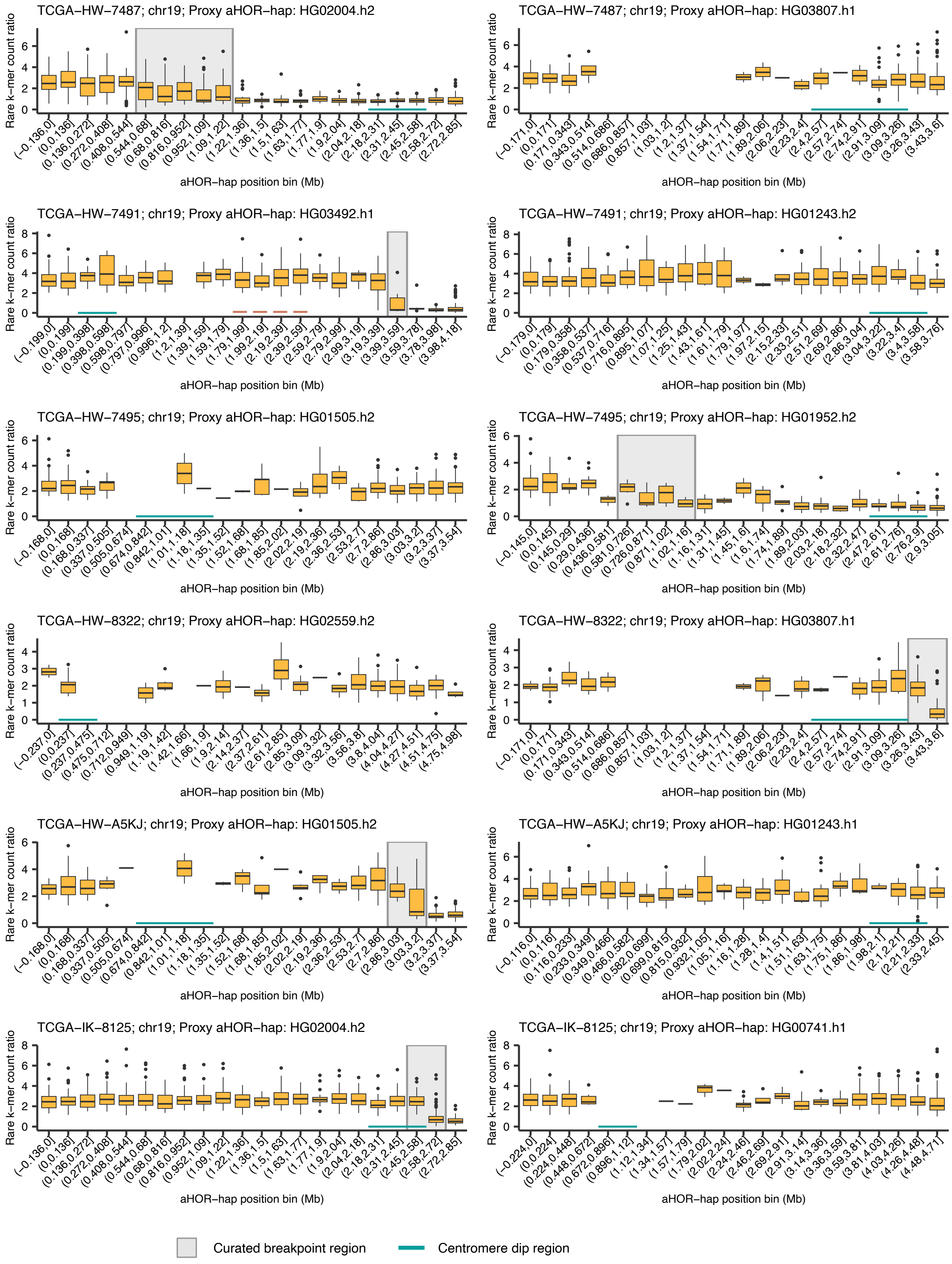


**Supplementary Figure. 7 (continued)**

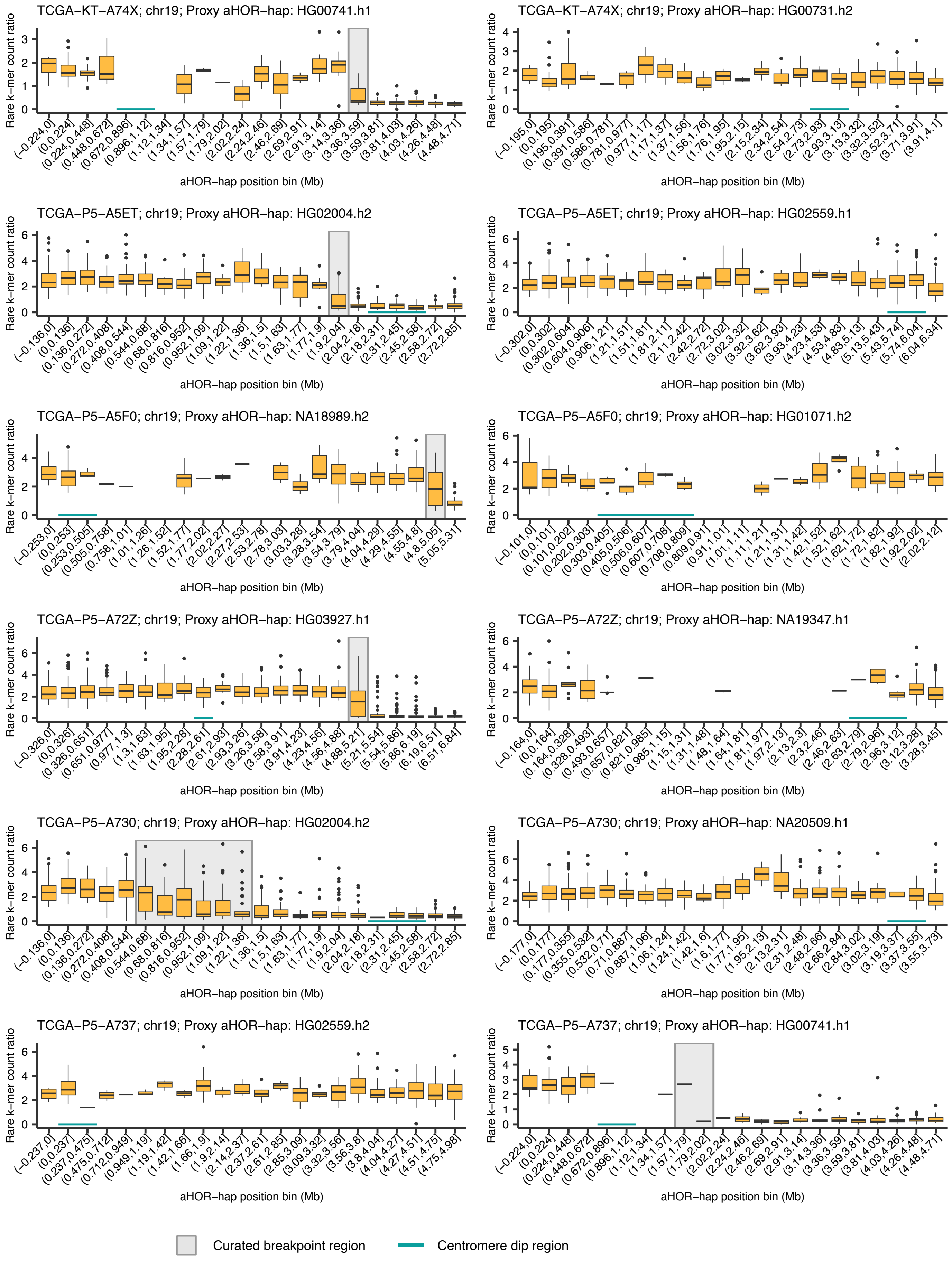


**Supplementary Figure. 7 (continued)**

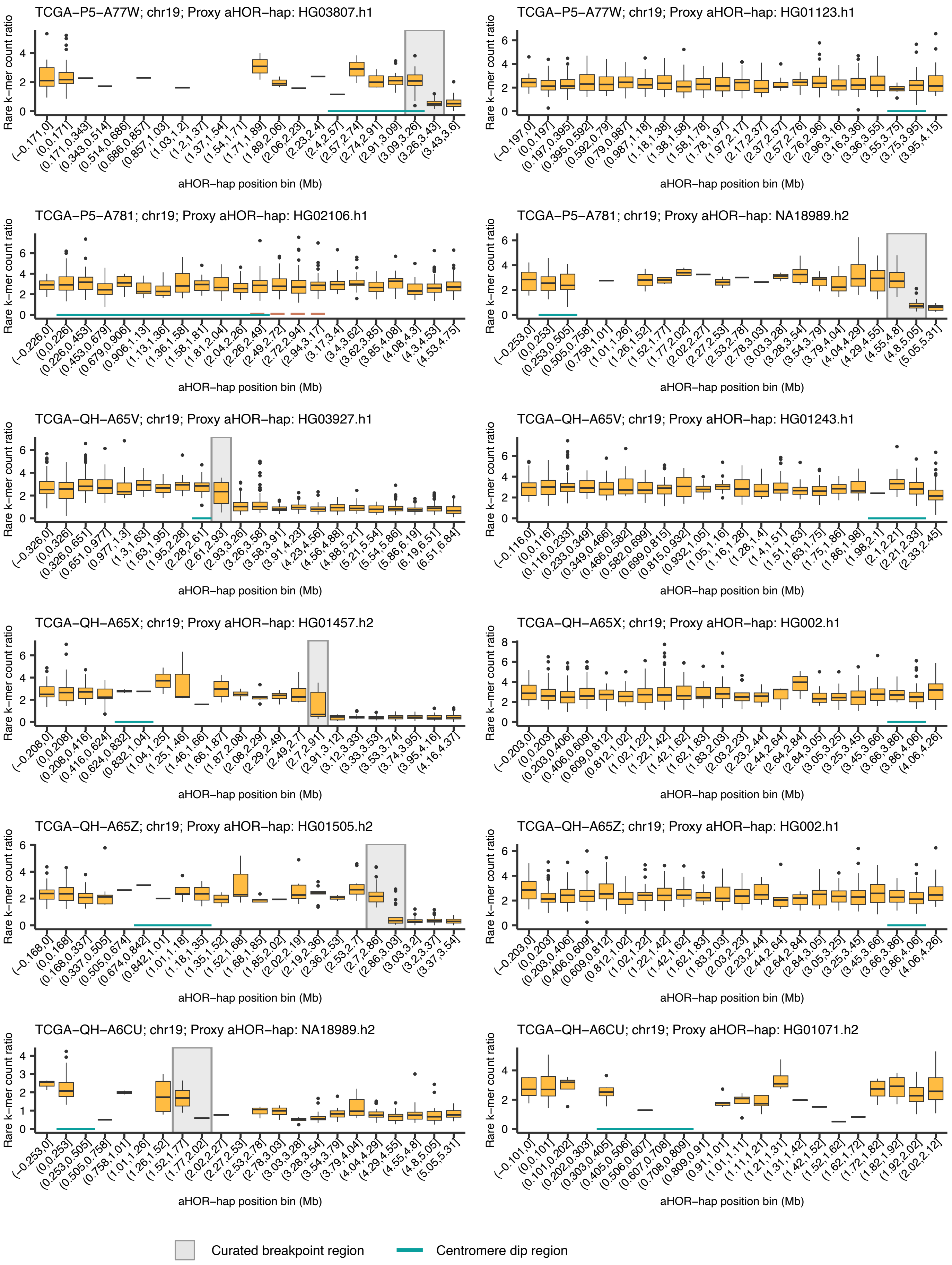


**Supplementary Figure. 7 (continued)**

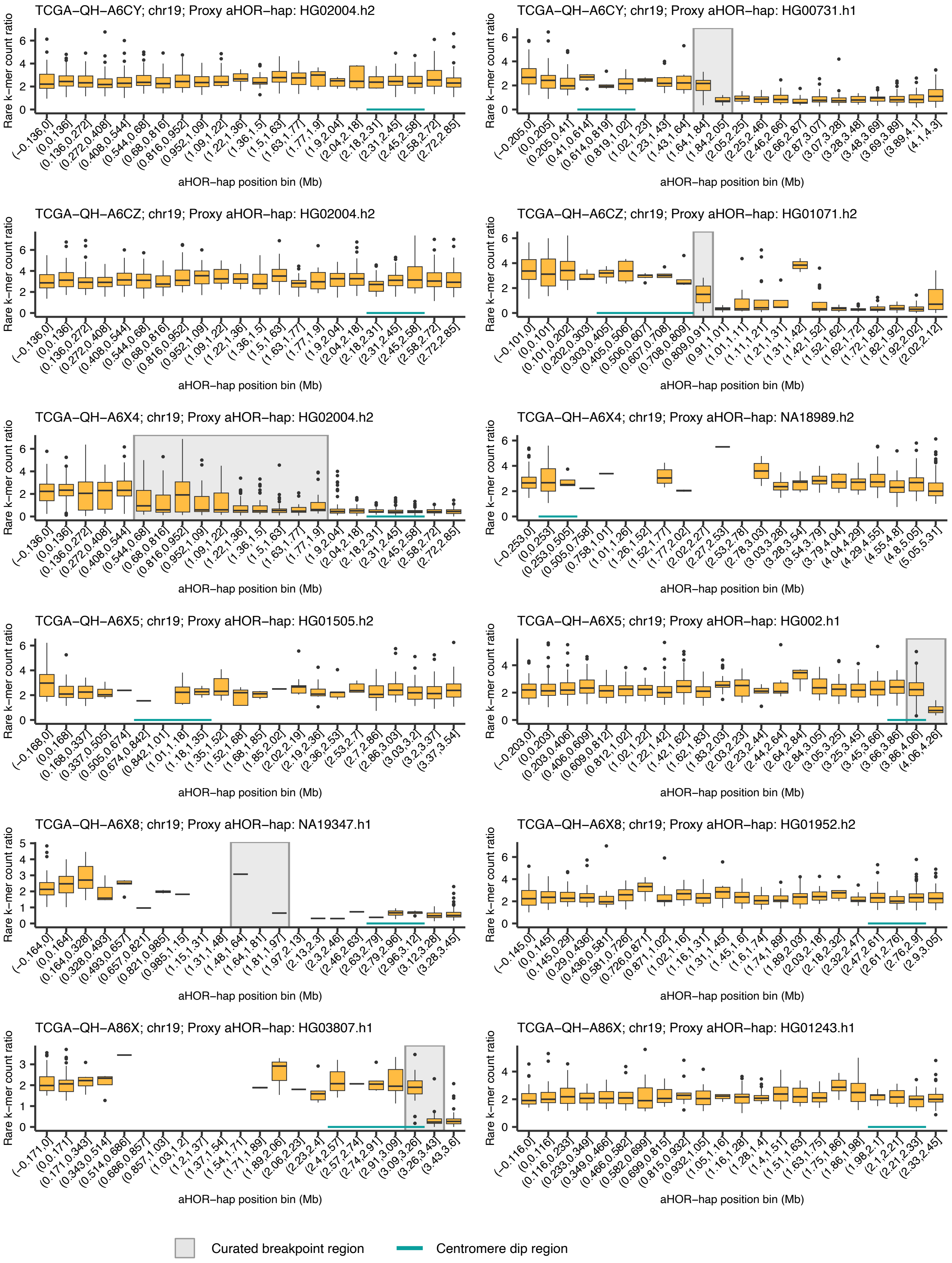


**Supplementary Figure. 7 (continued)**

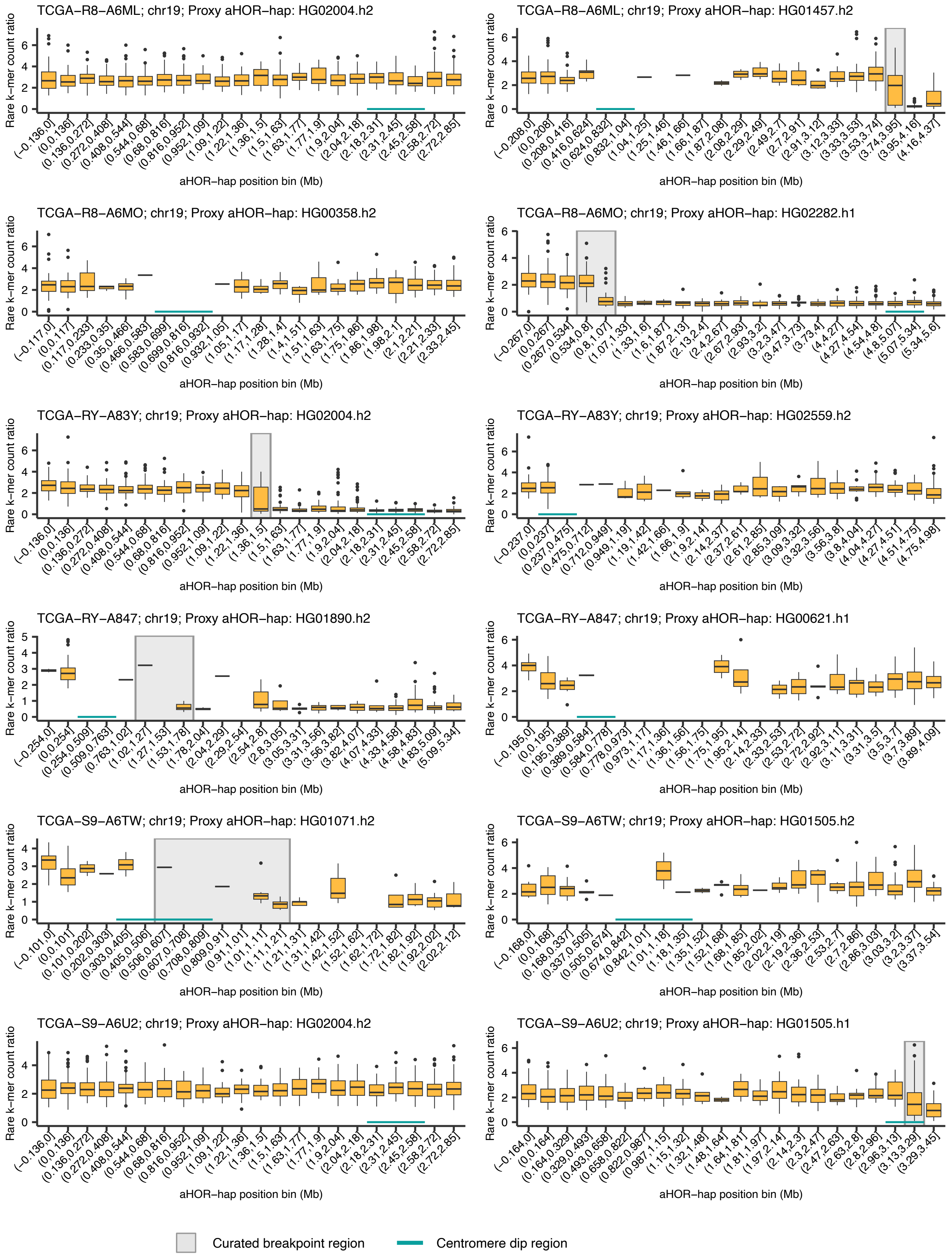


**Supplementary Figure. 7 (continued)**

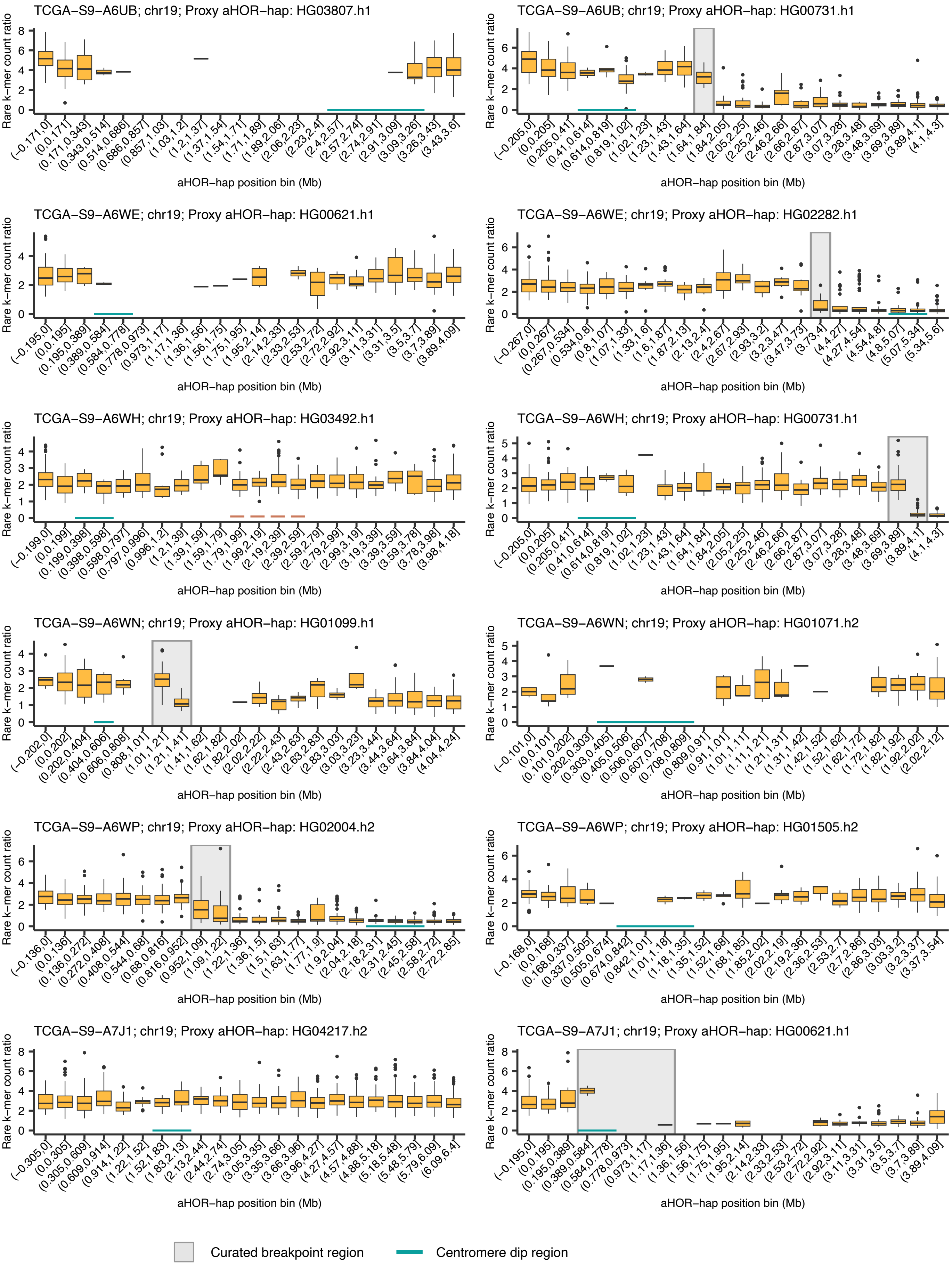


**Supplementary Figure. 7 (continued)**

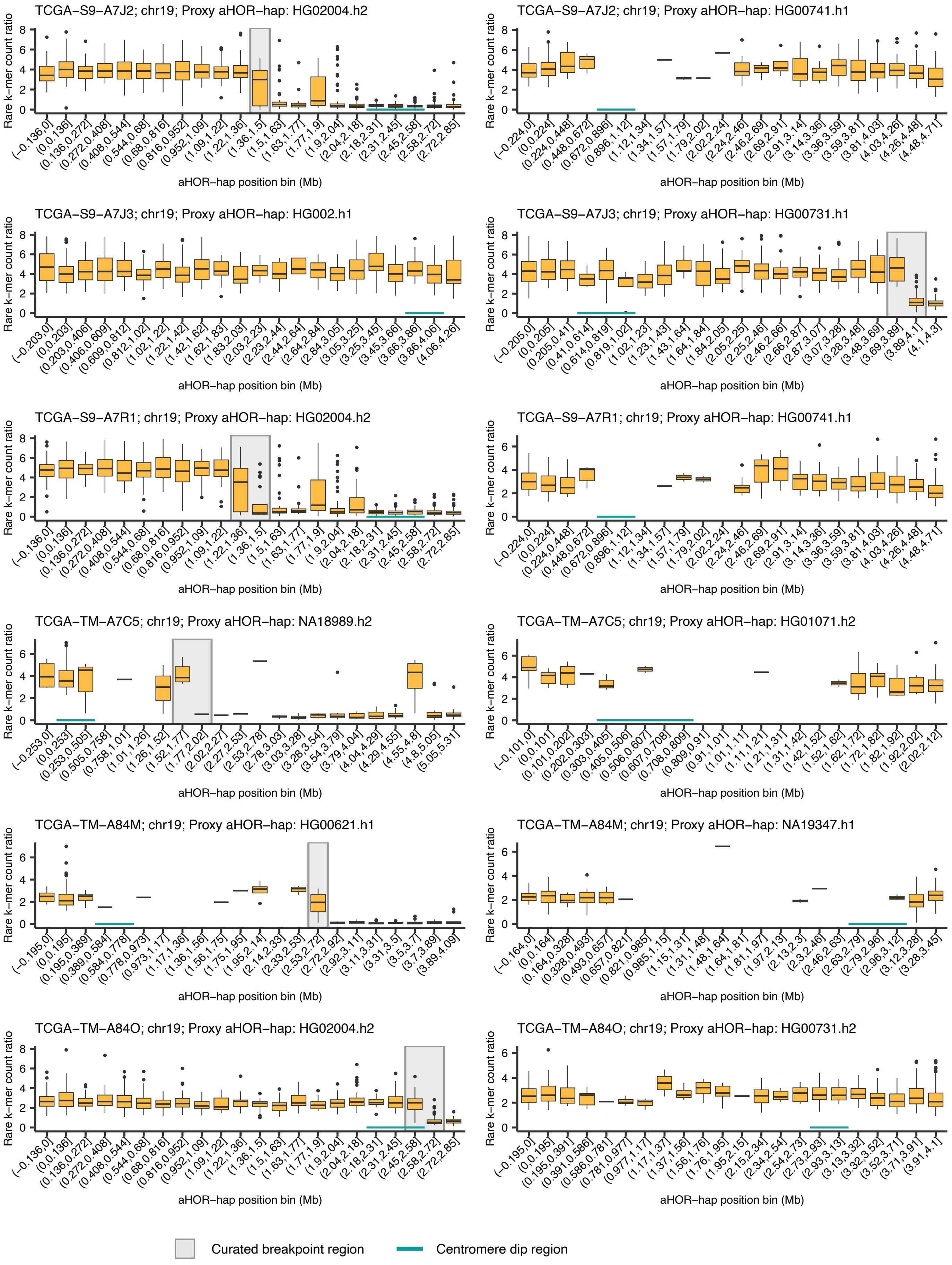


**Supplementary Figure. 7 (continued)**

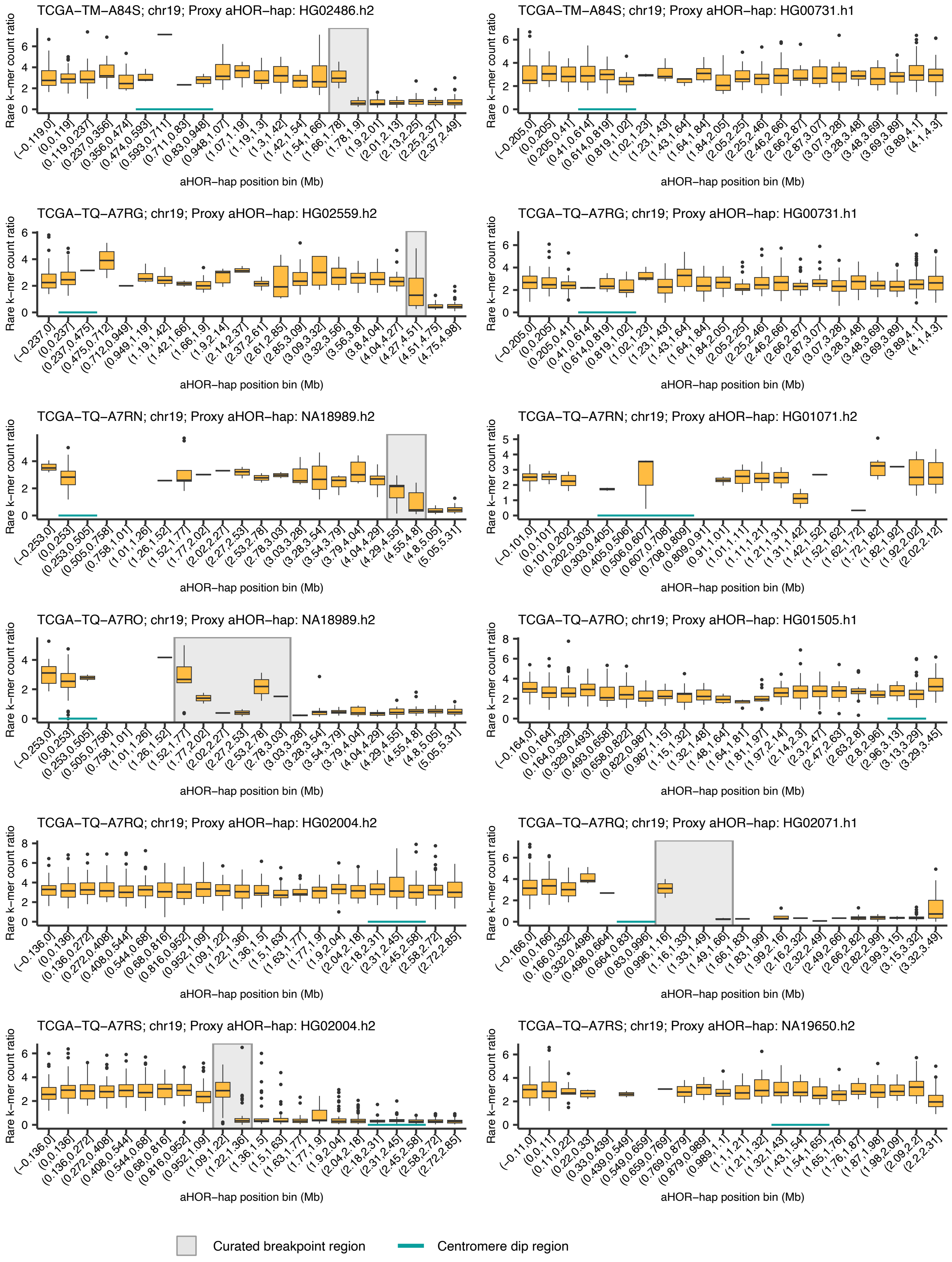


**Supplementary Figure. 7 (continued)**

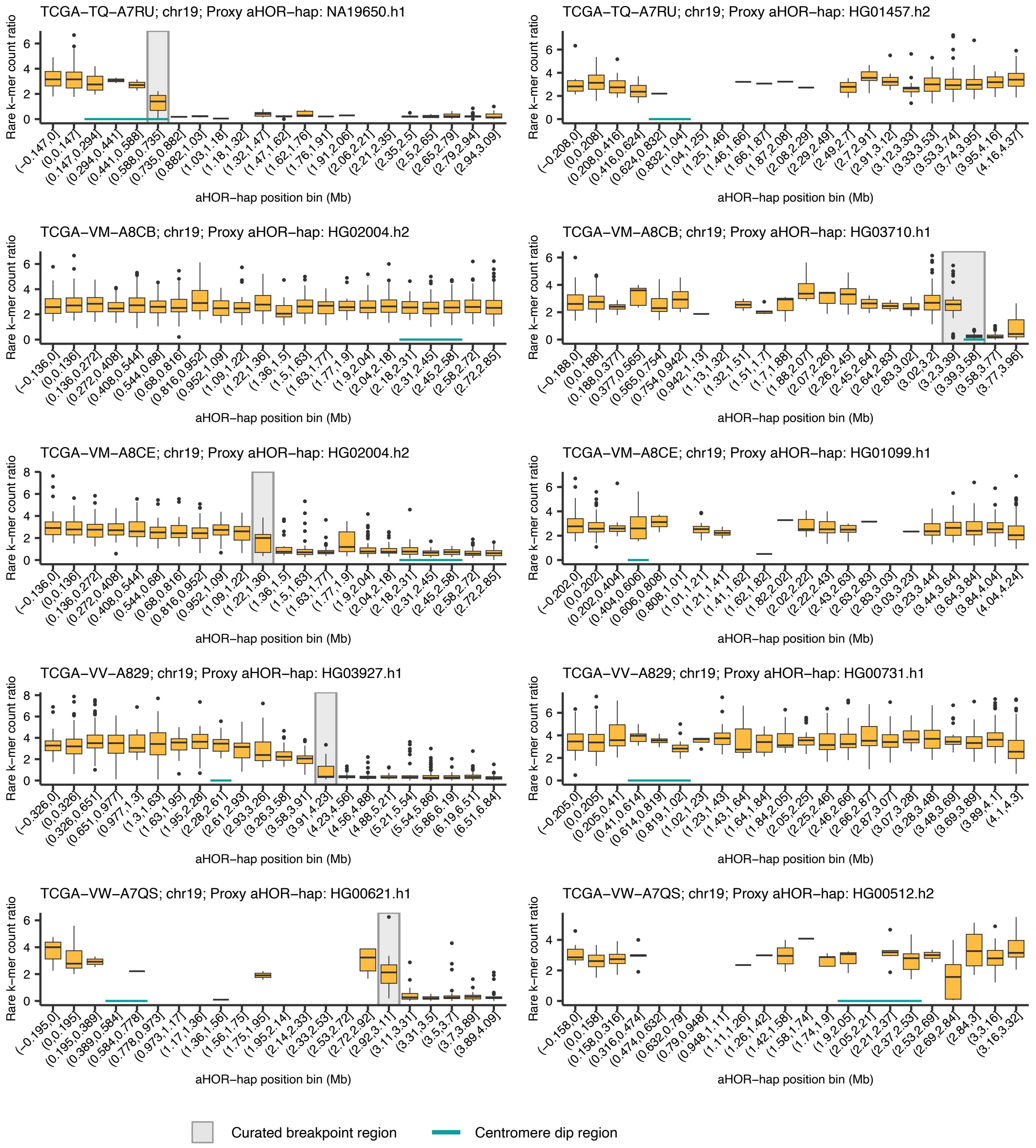


**Supplementary Figure. 7 (continued)**

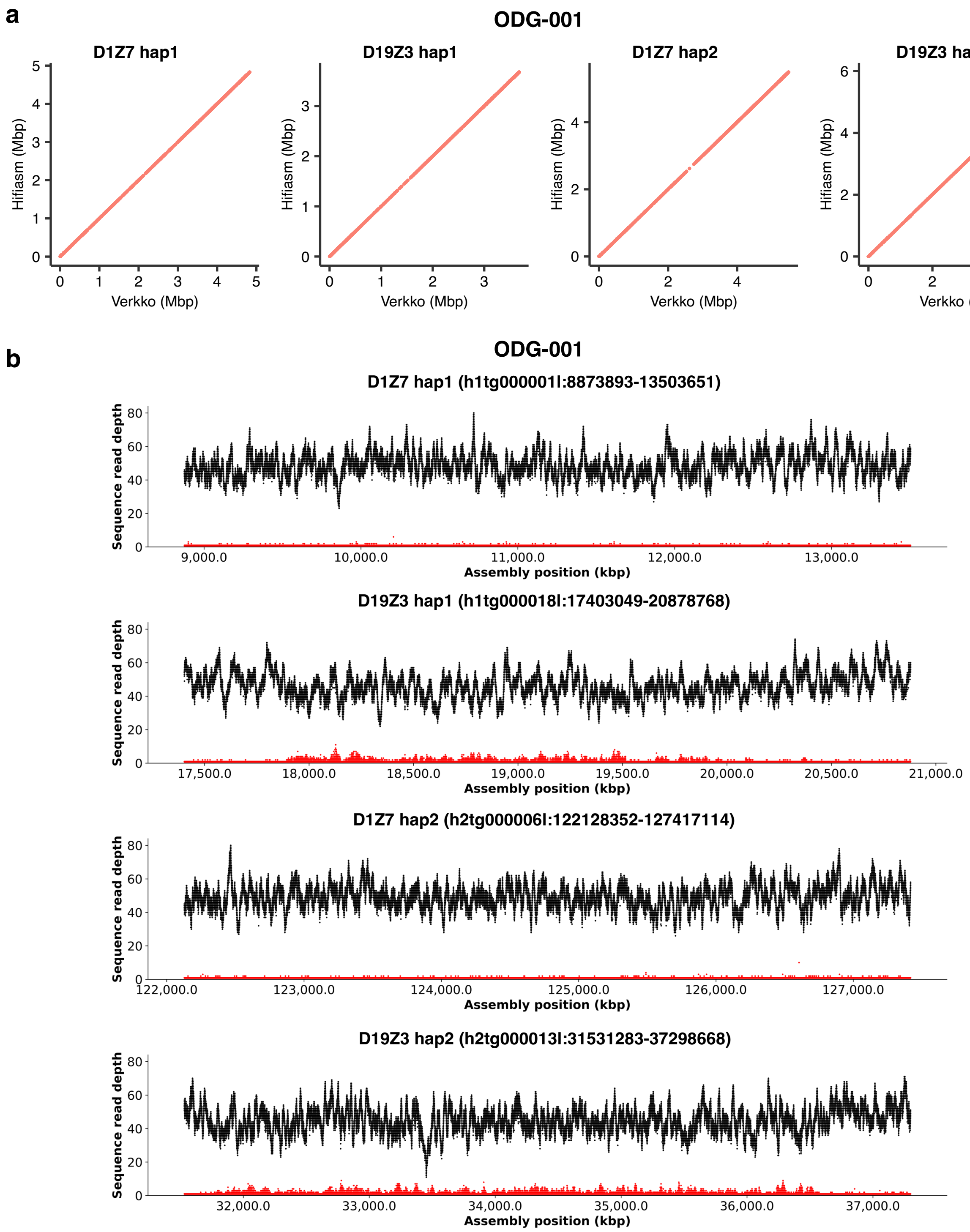


**Supplementary Figure 8: Assessment of centromere sequence assemblies for ODG-001.**

**(**a) Dot plots comparing Hifiasm and Verkko assemblies for the centromere sequences of chromosomes 1 (*D1Z7*) and 19 (*D19Z3*) in the sample ODG-001. Each dot represents shared unique k-mer (*k* = 27) between assemblies, demonstrating highly concordant assembly quality. (b) Sequence read depth across assembled centromeric haplotypes (*D1Z7* hap1, *D19Z3* hap1, *D1Z7* hap2, and *D19Z3* hap2). Black lines represent read depth, and red dots indicate the count of mismatch at each position. Genomic coordinates and contig identifiers are provided above each plot.

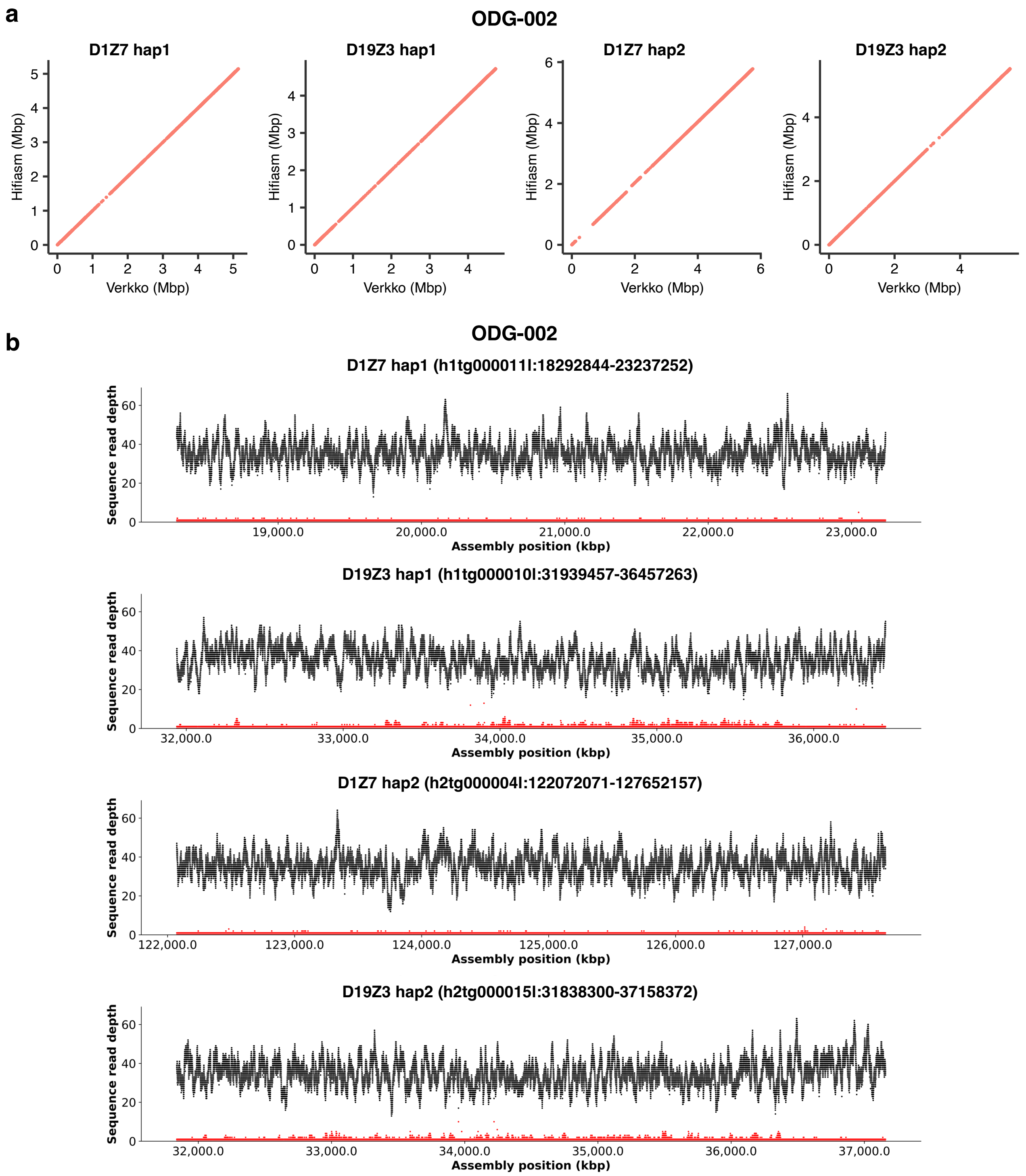


**Supplementary Figure 9: Assessment of centromere sequence assemblies for ODG-002.**

**(**a) Dot plots comparing Hifiasm and Verkko assemblies for the centromere sequences of chromosomes 1 (*D1Z7*) and 19 (*D19Z3*) in the sample ODG-002. Each dot represents shared unique k-mer (*k* = 27) between assemblies, demonstrating highly concordant assembly quality. (b) Sequence read depth across assembled centromeric haplotypes (*D1Z7* hap1, *D19Z3* hap1, *D1Z7* hap2, and *D19Z3* hap2). Black lines represent read depth, and red dots indicate the count of mismatch at each position. Genomic coordinates and contig identifiers are provided above each plot.

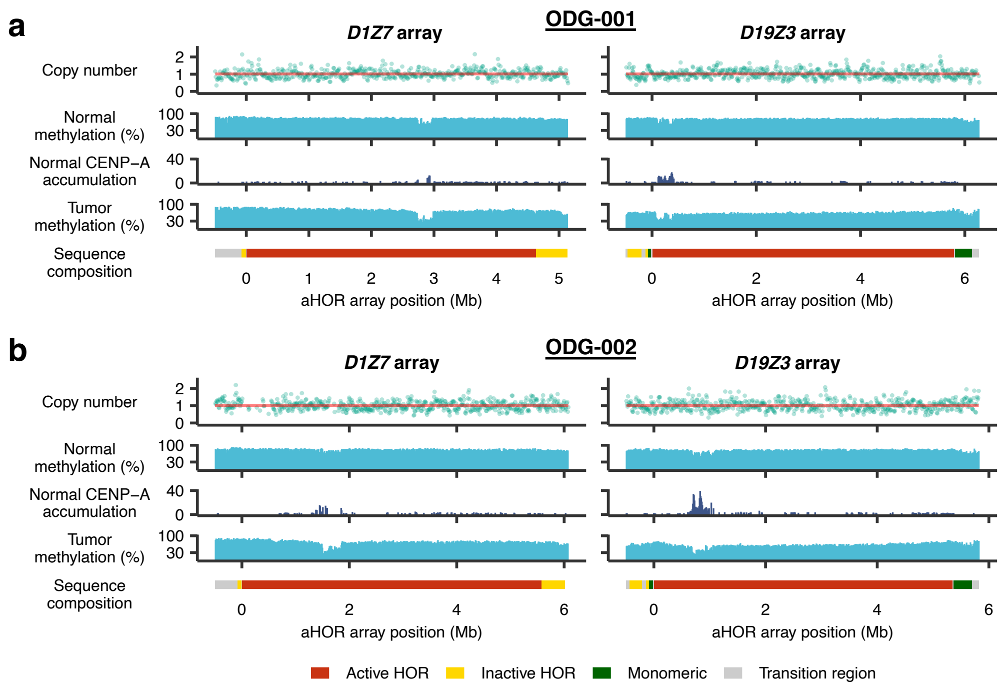


**Supplementary Figure 10: Rearrangement breakpoint mapping using long-read oligodendroglioma samples.** (a, b) Copy number profiles (normalized by ploidy), DNA methylation levels, and CENP-A concentrations at the *D1Z7* and *D19Z3* arrays in alleles not harboring translocation breakpoints in (a) ODG-001 and (b) ODG-002. DNA methylation is presented separately for normal and tumor tissues. CENP-A concentrations are shown in normal tissues. Sequence compositions indicating active α-satellite HOR arrays (Active HOR; red), inactive α-satellite HOR array (Inactive HOR; yellow), monomeric α-satellite (Monomeric; green) and transition region (Transition region; gray) are provided below. See also Fig. 5d, e.
